# Spatial transcriptomics of pediatric tuberculosis lymph nodes reveals a conserved hypoxia–SPP1 niche

**DOI:** 10.64898/2026.10.28.762471

**Authors:** S. Odunola Adebiyi, Pierre Goussard, Daniël R. De Wet, Jacques T. Janson, Tracey Jooste, Ayanda Shabangu, Anca Dorhoi, Stephen Bondoc, Michael Nekorchuk, Michael C. Chao, Sarah M. Fortune, Kathleen Busman-Sahay, Jacob D. Estes, Elya A. Shamskhou, Kevin B. Urdahl, Michael Y. Gerner, Gerhard Walzl, Nelita Du-Plessis

## Abstract

Pediatric tuberculosis (TB) often presents as primary disease involving the mediastinal lymph nodes; however, its granuloma microenvironment remains poorly characterized because these population patient tissues are rarely available for study. Here, we use spatial transcriptomics to profile 60 granulomas from four children aged 4 to 36 months. We identified nine recurrent transcriptional microenvironments shared across granulomas, despite their marked histological heterogeneity. One macrophage-rich microenvironment showed strong hypoxia-associated activity and *secreted phosphoprotein 1 (SPP1)* enrichment. This microenvironment was present in both non-necrotic and necrotic lesions and overlapped with a Mtb-persistence macrophage signature previously identified in murine TB lymph nodes. Pathway analysis revealed the enrichment of both antimicrobial and regulatory programs within this microenvironment, suggesting partial immune control rather than uniform inflammatory or suppressive activity in this region. SPP1 expression was associated with regions enriched for both M1- and M2-like macrophages, suggesting a macrophage program that extends beyond conventional polarization classifications and may be influenced by the local granuloma microenvironment. We further observed a similar spatial association between hypoxia-associated activity and SPP1 expression in an independent adult TB dataset (pooled Spearman ρ=0.65) and validated this association at protein level by multiplex immunofluorescence in our pediatric TB granuloma sample. Overall, this study identified a hypoxia-associated *SPP1*-enriched microenvironment that may contribute to incomplete bacterial clearance and could provide a framework for evaluating host-directed strategies to enhance sterilizing immunity without exacerbating tissue damage.

## Background

Pediatric tuberculosis (TB) accounts for 12% of the global TB burden, with over 175, 000 deaths annually^1^. More than 50% of pediatric TB cases remain undiagnosed thereby increasing the risk of severe complications such as airway compression, disseminated TB, and TB meningitis^2–4^. TB in children is not a scaled-down version of adult TB. It has distinct clinical and biological features that differ from those of adults. Pediatric TB is predominantly a primary TB disease that often involves the mediastinal lymph nodes. Adult TB, on the other hand, often presents as a post-primary disease of the lungs, reflecting years of chronic host-pathogen interactions at these sites^5–7^. These distinct clinical presentations suggest important differences in the underlying biology of the disease, further supported by the markedly greater protective efficacy of the BCG vaccine in children than in adults^8,9^. Despite these fundamental differences, mechanisms underlying childhood TB remains understudied, largely due to difficulty in accessing diagnostic samples, ethical constraints surrounding tissue collection in children, and the paucibacillary nature of pediatric TB samples^10^. Thus, our understanding of the cellular organization and tissue microenvironment underlying pediatric TB relies on histological observation and extrapolation from molecular studies of adult TB tissue.

A defining feature of TB is the ability of the pathogen *Mycobacterium tuberculosis* (Mtb) to persist despite a robust systemic immune response^11^. This suggests that protection against TB is largely determined at the site of infection, rather than by systemic immunity alone^12^. The granuloma, the primary host-pathogen interaction site in TB, is established by the organized aggregation of immune and stromal cells. It functions to contain and eliminate Mtb but can also provide a niche that permits bacterial persistence when protective immunity fails^13–15^. Studies from adult human samples and non-human primates (NHP) have shown that individual granulomas within a host differ markedly in terms of their sterilizing capacity, bacterial burden, cellular composition, reactivation, and inflammatory state^16–20^. The cumulative effect of granulomas in each host determines the clinical outcome in such an individual^21^. Granuloma trajectory is highly dependent on the spatial organization of cells than on their bulk composition, as the same cell types organized differently can have different functional consequences^18,22–24^. With recent advances in transcriptomic technology, there has been an increasing number of studies evaluating the spatial transcriptomic organization in granulomas to identify specific tissue microenvironments and cellular states^23,25–27^. However, nearly all studies have focused on adult pulmonary tissue, reflecting post-primary disease dynamics. Whether the spatial principles governing adult granulomas are conserved in pediatric, primary TB remains unknown.

Macrophages are the central players in TB granulomas. They are both the primary cellular target of Mtb and the principal effector cells of granuloma-based immunity^28,29^. Over the years, multiple functionally distinct macrophage states have been described in TB. Histologically, granulomas are known to be composed of untransformed macrophages, foamy macrophages (often associated with Mtb persistence), epithelioid macrophages, and multinucleated giant cells^15,30,31^. However, transcriptomic studies have revealed additional macrophage states in granulomas. This includes TREM2+ macrophages, GRN+ macrophages, Arg1+ macrophages, IDO1+ macrophages and recently SPP1+ macrophages, which were identified as a prominent feature of adult TB granulomas^23,27,32–36^. All these macrophage states have been characterized in bulk- and single-cell profiling, but their spatial organization within granulomas is only beginning to be resolved. The tissue microenvironment and metabolic context that shape where and when each macrophage state emerges, remain incompletely defined. Understanding this spatial-environmental logic is important because different macrophage states have distinct implications for bacterial control and persistence. This knowledge gap is pronounced in pediatric TB, where no spatial characterization of granuloma organization currently exists.

To address this knowledge gap, we applied the 10x Genomics Visium platform to profile the spatial organization of granulomas from surgically resected mediastinal lymph nodes of 4 pediatric TB patients. We performed histological annotation and profiling of 60 granulomas across 7 tissue sections. We identified nine recurrent transcriptional microenvironments present across lesions, including a distinct hypoxia-associated niche present in both necrotic and non-necrotic granulomas. This niche was enriched for *SPP1* expression, which is localized to predominantly monocyte-derived macrophage-rich spots in the granulomas. Multiplex immunofluorescence confirmed this enrichment of SPP1 in hypoxia region at protein level and showed that SPP1 track with M1 and M2 macrophage phenotype spatial location. The hypoxia-SPP1 axis replicated in an independent adult TB cohort spanning pulmonary, pleural, and lymphoid sites and converged with a persistence-associated macrophage signature from murine TB lymph node infection. To our knowledge, these findings provide the first spatial transcriptomic characterization of pediatric TB granulomas, identifying a conserved hypoxia-SPP1 axis in monocyte-derived macrophage-rich regions as a candidate feature of local immune permissiveness that spans pediatric primary and adult post-primary human TB.

## Results

### Histopathology of pediatric mediastinal lymph node TB granulomas

To examine the spatial biology of primary TB, we obtained archival formalin-fixed paraffin-embedded (FFPE) mediastinal lymph node samples from four pediatric patients aged 4 to 36 months. All underwent surgical resection for severe airway compression caused by intrathoracic tuberculous lymphadenitis. All patients were HIV-negative, and all samples tested culture and GeneXpert-positive for Mtb by sputum, bronchoalveolar lavage (BAL) or gastric washing (supplementary Table 1). Histological review of hematoxylin and eosin-(H&E) and ziehl-neelsen-(ZN) stained sections by a qualified pathologist identified 66 histologically distinct granulomas across the four tissue blocks (Fig.1B). six granulomas were excluded from the current analysis as these were classified as very early micro-granulomas. The remaining 60 were profiled using 10x Genomics Visium Spatial Transcriptomics (Fig.1A). Acid-fast bacilli were sparse across all blocks, consistent with the current literature on pediatric TB tissue^37^ and the initiation of anti-tuberculous therapy prior to surgery.

**Fig. 1.**
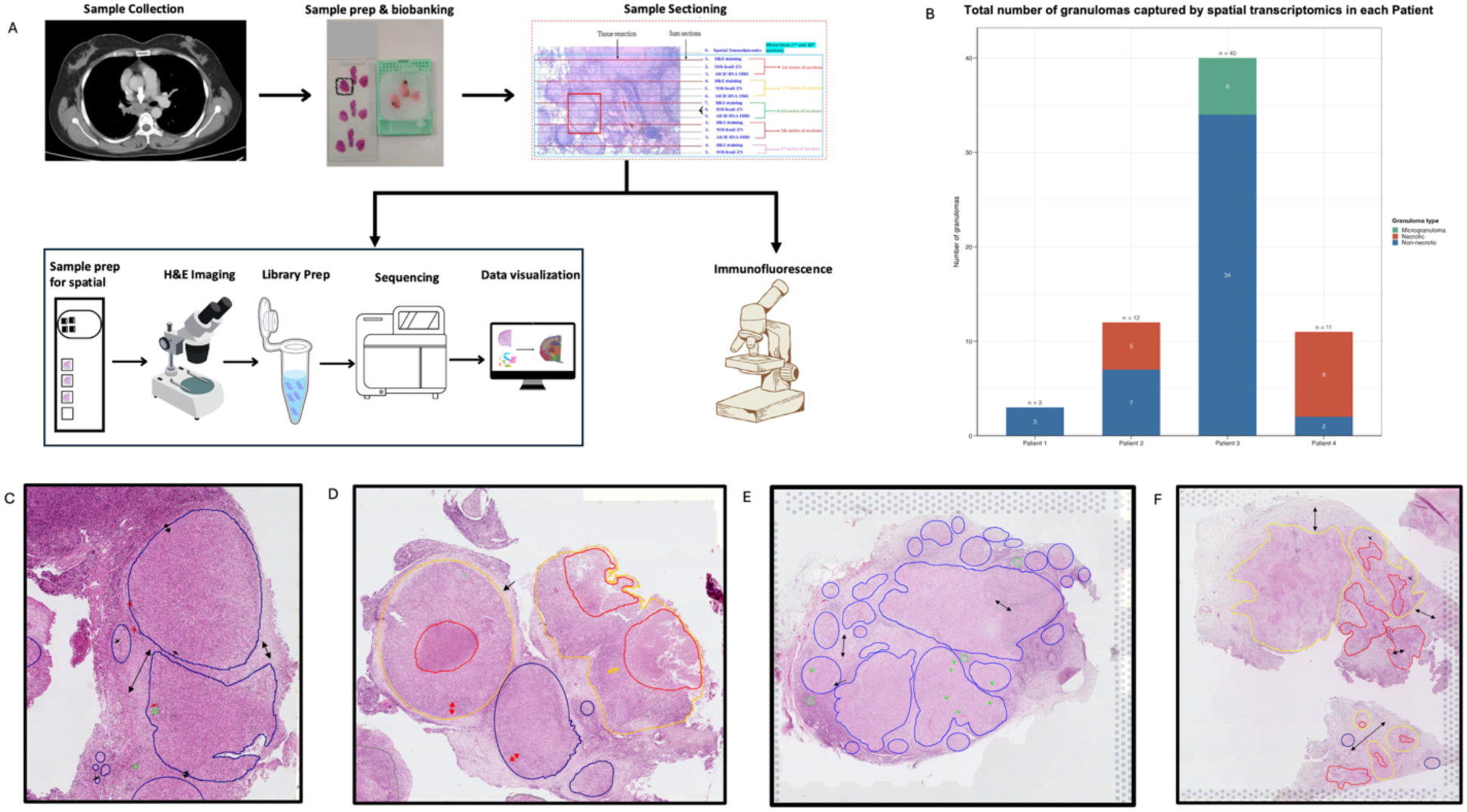
Study design and granuloma annotation. **(A)** Overview of the study workflow. Archival FFPE mediastinal lymph node tissue was collected from four paediatric TB patients aged 4–36 months. Sections were processed for spatial transcriptomics (10x Genomics Visium), and adjacent sections were used for H&E staining, Ziehl–Neelsen (ZN) staining, and immunofluorescence. **(B)** Number of granulomas captured by Visium in each patient, coloured by granuloma type (non-necrotic, necrotic, and microgranuloma); totals are shown above each bar. **(C–F)** H&E-stained sections from Patient 1 **(C)**, Patient 2 **(D)**, Patient 3 **(E)**, and Patient 4 **(F)** as placed on the Visium capture area, with pathologist-annotated granulomas and tissue structures. Blue outlines indicate non-necrotic granulomas, yellow outlines indicate necrotic granulomas, red outlines indicate necrotic cores, and green circles indicate microgranulomas. Red arrows indicate lymphocytic zones, green arrows indicate multinucleated giant cells, and black arrows indicate fibrosis.

Granuloma morphology varied substantially across the cohort (Fig. 1C-E) with both necrotic and non-necrotic granulomas represented within and across tissue blocks. These lesions varied in the degree of fibrosis, lymphocyte infiltration, macrophage organization, and multinucleated giant cell formation. Patient_1 tissue was dominated by lymphocyte-rich non-necrotic granulomas with extensive fibrosis (Fig. 1C). Patient_2 contained both classical zonated necrotic granulomas and less circumcised necrotic lesions, alongside non-necrotic lymphocyte rich granulomas embedded in mature fibrous stoma (Fig. 1D). Patient_3 tissue was made up of multifocal, non-necrotic granulomas rich in epithelioid macrophages and Langhans-type giant cells (Fig. 1E). Patient_4 tissue contained coalescing necrotizing granulomas with poorly demarcated caseous zones, little fibrotic encapsulation, and several non-necrotic granulomas (Fig. 1F). These samples span the histological range already described in TB lymphadenitis, thereby providing a suitable basis for spatial transcriptomic analysis.

### Cellular composition of pediatric mediastinal lymph node TB granulomas is dominated by inter-individual variability rather than necrotic status

Given the histological diversity across this cohort, we asked whether granuloma cellular composition differed by necrotic status. We deconvolved sixty histologically annotated granulomas from four patients and seven sections (two sections from each sample, one sample section was excluded due to lifting during processing) using Robust Cell Type Decomposition (RCTD) against a NHP TB lymph node scRNA-seq reference^17,38^ (Fig. 2A). Macrophages and fibroblasts together accounted for 56% of deconvolved cell-type proportions across granuloma spots. Other immune cell populations were present at lower and variable abundances (Fig. 2C).

**Fig 2.**
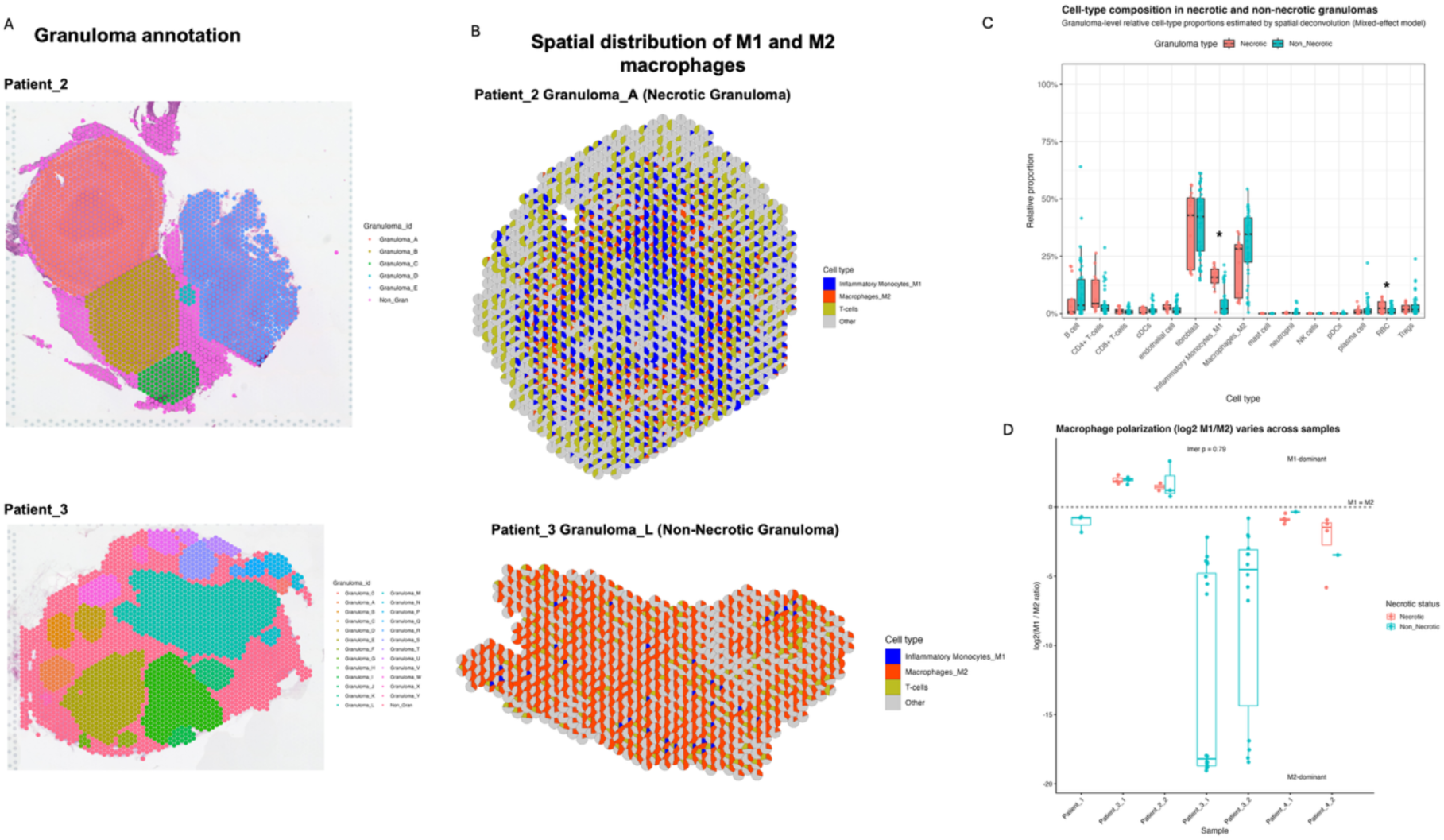
Cellular composition of necrotic and non-necrotic pediatric mediastinal lymph node TB granulomas is dominated by inter-individual variability. **(A)** Representative Visium spatial transcriptomics sections from Patient_2 and Patient_3 showing histology-guided granuloma annotations.Each colour represents an individual granuloma. The two sections are representative of seven sections analysed across four patients (2 sections each from 3 individual and a section from a fourth). **(B)** Spatial distribution of inflammatory monocytes (M1-like, blue), M2-like macrophages (orange), and T cells (yellow) within a representative necrotic granuloma (Patient_2 Granuloma_A) and non-necrotic granuloma (Patient_3 Granuloma_L) granulomas inferred by RCTD spatial deconvolution. Each Visium spot is represented as a pie chart showing the estimated cell-type proportions; other cell types are grouped as other(grey). **(C)** Granuloma-level cell-type proportions across necrotic (n = 14, salmon) and non-necrotic (n = 46, teal) granulomas.Each point represents one granuloma. Box plots show median and interquartile range. Significance reflects linear mixed-effects modelling with patient and section-within-patient as nested random effects (* FDR < 0.05). **(D)** Granuloma-level macrophage polarisation, expressed as log_2_(M1/M2) ratio and stratified by patient sample. Each point represents one granuloma; box plots show median and interquartile range. The dashed line indicates equal M1 and M2 contributions. Necrotic and non-necrotic granulomas are shown in salmon and teal, respectively.

Cohort-level Wilcoxon testing suggested that inflammatory monocytes (M1-skewed; Supplementary method) were enriched in necrotic granulomas (FDR= 3.3 x 10^-4^; Supplementary Fig. 1B). However, because granulomas from the same patient are not statistically independent, we repeated the analysis using a linear mixed-effects model that accounted for patient and tissue section^39^. Under this model, M1 enrichment was markedly reduced in necrotic granulomas compared to what was observed with the uncorrected Wilcoxon test (FDR = 0.026). In addition, red blood cells were also enriched in necrotic granulomas (FDR= 0.020), likely reflecting hemorrhage due to small vessel destruction. No differences for other cell types reached statistical significance (Fig. 2C). A within-individual sensitivity analysis restricted to patients containing both granuloma types failed to recover M1 signal, though power was limited by the small number of contributing patients (Supplementary Fig. 1C). A similar pattern was observed for macrophage polarization where the cohort-level log_2_ (M1/M2) ratio per granuloma was significantly higher in necrotic granulomas (Wilcoxon p = 6 x 10^-4^) suggesting an M1-skewed profile in necrotic lesions. However, this difference was not observed with nested mixed-effects analysis (p = 0.79) (Fig. 2D). We found that patient identity explained the largest share of variance (patient variance = 25.6 vs section-within-patient variance = 4.4 and residual variance = 30.4) in the ratio of log_2_ (M1/M2). Macrophage polarization followed patient-specific patterns that were largely preserved across granulomas within the same patient regardless of their necrotic status. These findings indicate that cellular composition is shaped more strongly by inter-individual immune context than granuloma necrotic status. We therefore turned to spatial organization and transcriptional programs within granulomas.

### Integrated analysis identifies nine recurrent transcriptional microenvironments across granulomas

To examine granuloma spatial organization directly, we extracted all 14,226 spatial spots across the sixty granulomas and integrated them across samples. Prior to integration, spots clustered predominantly by patient, reflecting individual-specific profiles (Fig. 3A, left). Batch-aware integration using Reciprocal Principal Component Analysis (rPCA) reduced this separation and identified nine transcriptional microenvironments (MEs) conserved across patients and granulomas (Fig. 3A (left & right) & B). The emergence of these shared MEs despite strong inter-individual variability, suggests that pediatric mediastinal lymph node granulomas converge on a limited number of recurrent spatial programs. This is in line with prior work demonstrating that granulomas are organized into functionally distinct tissue compartments^18,22^.

**Fig 3.**
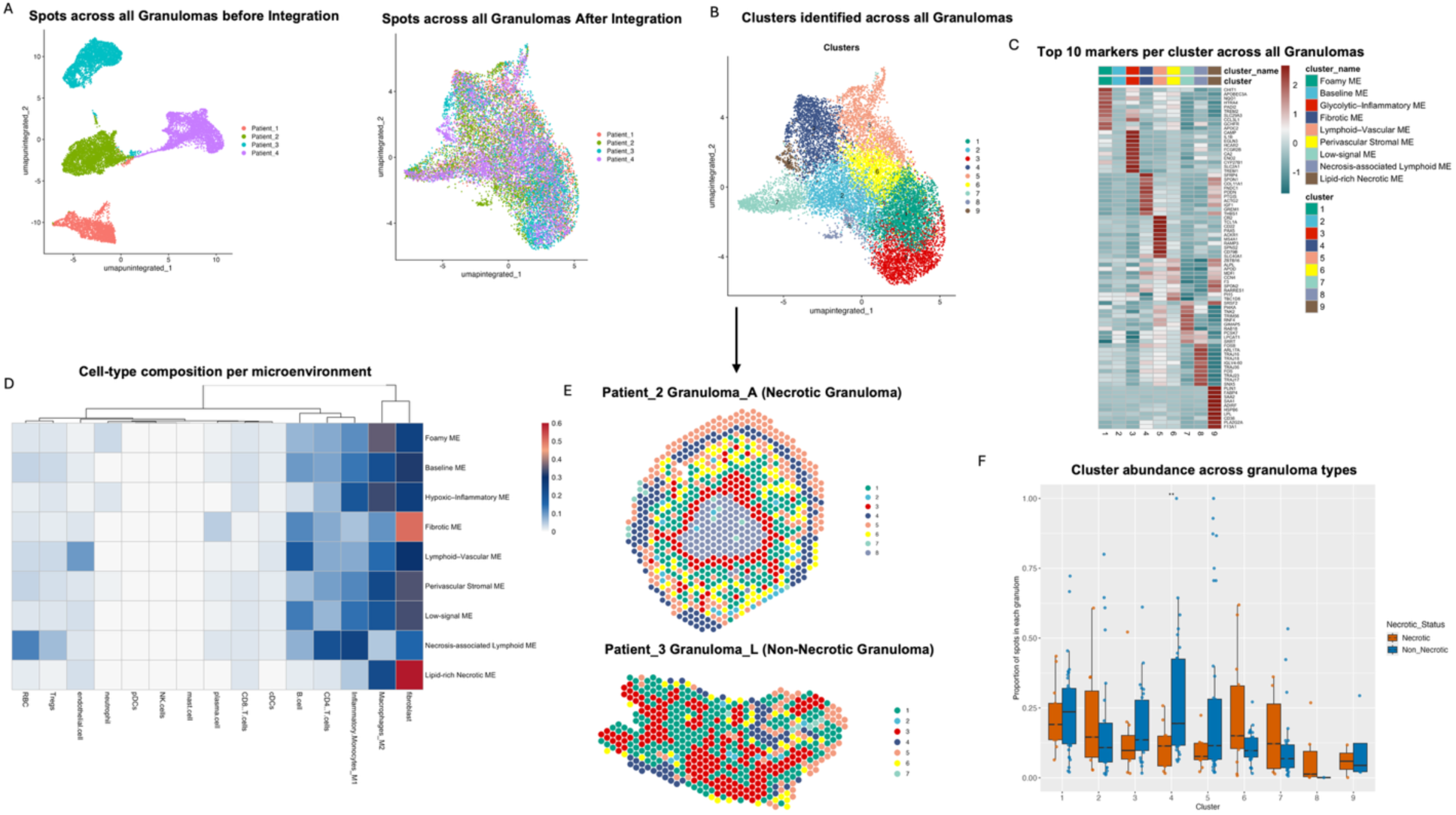
Paediatric mediastinal lymph node TB granulomas are organised into nine recurrent spatial microenvironments. **(A)** UMAP of all 14,226 spatial spots before (left) and after (right) integration with reciprocal PCA (rcPCA), coloured by patient. **(B)** Unsupervised clustering of the integrated UMAP identifies nine transcriptionally distinct microenvironments (MEs). **(C)** Heatmap showing the top 10 marker genes for each ME. Marker genes were identified using the MAST test (log₂FC > 0.35, minimum detection fraction = 0.30). Colour scale represents scaled average expression. Functional annotations assigned to each ME are shown on the right. **(D)** Mean deconvolved cell-type composition of each ME estimated using RCTD with a non-human primate lymph node single-cell reference. Colour intensity indicates the mean estimated proportion of each cell type. Columns are hierarchically clustered according to similarity. **(E)** Spatial projection of ME assignments onto representative necrotic (Patient_2 Granuloma_A) and non-necrotic (Patient_3 Granuloma_L) granulomas. Colours correspond to the nine MEs identified in panel B. (F) Relative abundance of each ME in necrotic (orange, n = 14) and non-necrotic (blue, n = 46) granulomas. Each point represents one granuloma. Box plots show the median and interquartile range. Statistical significance was assessed using linear mixed-effects models with patient and tissue section nested within patient as random effects and BH correction for multiple testing. No ME reached FDR < 0.05; the Fibrotic ME (Cluster 4) showed the strongest trend toward enrichment in necrotic granulomas (FDR = 0.072, mixed effects).

The nine MEs represented distinct combinations of transcriptional programs and cellular composition rather than discrete cell types^38,40^ (Fig. 3C & D). Three were macrophage-associated, namely, a foamy macrophage ME (cluster 1) marked by lipid-laden macrophage genes, a baseline macrophage ME (cluster 2) which lacked a dominant transcriptional program, and the glycolytic-inflammatory macrophage ME (cluster 3) co-expressing inflammatory mediators and glycolytic genes. Two stromal MEs were also identified including a fibrotic stromal ME (cluster 4) marked by extracellular matrix gene expression and a perivascular stromal ME (cluster 6) expressing perivascular fibroblast markers. A lymphoid-vascular ME (cluster 5) co-expressing B cell and endothelial markers was also present.

Three MEs (clusters 7 – 9) localized preferentially to histologically necrotic regions (Fig. 3E). The low signal ME (cluster 7) showed weak, non-specific expression, likely reflecting reduced transcriptional output in necrotic tissue. The necrosis-associated lymphoid ME (cluster 8) co-expressed T cell receptor and immunoglobulin genes, and because these granulomas originated from mediastinal lymph nodes, this ME likely reflects the underlying tissue architecture and not direct lymphocyte infiltration of the necrotic core. The final ME localized preferentially to histologically necrotic regions, is a lipid-rich necrotic ME (cluster 9) defined by lipid metabolism, macrophages and acute-phase response genes. This matches the lipid remodeling typical of caseous necrosis in granuloma ^30,41^. This ME appeared in only a subset of the granulomas. Marker gene expression and RCTD deconvolution supported the annotation of each microenvironment (Fig. 3E; Supplementary Fig 2 & 3B; Supplementary Note 1).

Projection of the MEs back onto tissue sections showed that each occupied spatially coherent domains within granulomas (Fig. 3E) but that their arrangement varied across lesions. Some granulomas displayed concentric organization resembling classical granuloma architecture^18^, whereas others showed more diffuse distributions of the same MEs.

We next asked whether any ME was associated with necrotic status (Fig. 3F). Cohort-level analysis identified the fibrotic stromal ME (cluster 4) as differentially represented (FDR = 0.035). However, this significance was lost when applying nested mixed modelling with patient and section-within-patient as random effects (FDR = 0.072, estimate = 0.167). No other ME reached significance and as seen with cellular composition, the dominant axis of ME variation was inter-individual rather than necrotic status (Fig. 3A).

### Convergent identification of a hypoxia-associated microenvironment within TB granulomas

We next investigated whether any of the MEs were associated with specific immunological functional programs. Gene set enrichment analysis of each cluster marker genes identified pathways consistent with the assigned identities of several ME annotations. These included the epithelial-mesenchymal transition pathway in the fibrotic and perivascular stromal MEs, the fatty acid metabolism pathway in the foamy macrophage ME, and the interferon signaling pathway in the lymphoid-vascular ME (Supplementary Fig. 4; Supplementary Table 2). Among all nine MEs, only the glycolytic-inflammatory macrophage ME (cluster 3) showed significant enrichment of the Hallmark Hypoxia pathway (NES = 1.39, FDR = 0.0076) alongside upregulation of glycolysis and TNF-α signaling via NF-κB (Fig. 4A). This co-enrichment pattern points to HIF-associated metabolic adaptation and inflammatory activation^42,43^.

**Fig 4.**
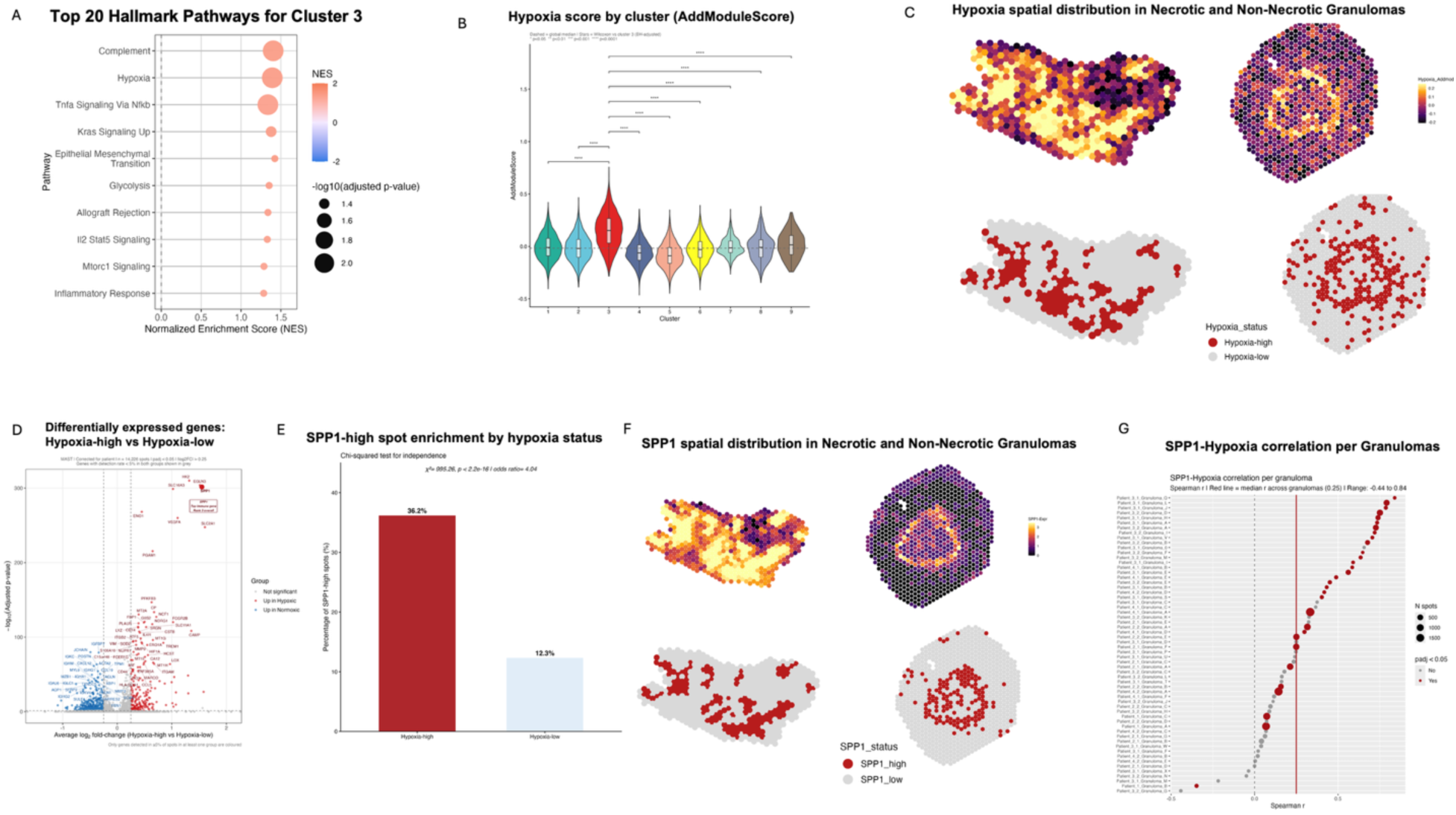
Identification of a hypoxia-associated microenvironment enriched for SPP1 in paediatric TB granulomas. **(A)** Gene set enrichment analysis of Cluster 3 marker genes against the MSigDB Hallmark collection. Dot size indicates statistical significance (−log10 adjusted P value) and colour indicates the normalised enrichment score (NES). **(B)** Hypoxia module scores across the nine microenvironments calculated using AddModuleScore. Cluster 3 showed the highest hypoxia score. Statistical significance was assessed using the Wilcoxon rank-sum test with Benjamini–Hochberg correction (**** adjusted P < 0.0001). Boxes within violins indicate the median and interquartile range. **(C)** Spatial distribution of hypoxia module scores (top) and binary hypoxia classification (bottom; red, hypoxia-high; grey, hypoxia-low) in representative non-necrotic and necrotic granulomas. **(D)** Volcano pl. Volcano plot showing differential gene expression between hypoxia-high (n = 3,394 spots) and hypoxia-low (n = 10,832 spots) regions using MAST with patient identity included as a latent variable. Red, genes upregulated in hypoxia-high regions; blue, genes upregulated in hypoxia-low regions; grey, not significant or lowly detected genes. **(E)** Percentage of SPP1-high spots in hypoxia-high and hypoxia-low regions. Statistical significance was assessed using Pearson’s chi-squared test. **(F)** Spatial distribution of SPP1 expression (top) and binary SPP1 classification (bottom; red, SPP1-high; grey, SPP1-low) in the same representative granulomas shown in panel C. **(G)** Per-granuloma Spearman correlation between hypoxia score and SPP1 expression across all 60 granulomas. Each point represents one granuloma. Red points indicate correlations remaining significant after Benjamini–Hochberg correction.

This finding was unexpected because hypoxia is usually associated with necrotic and caseating TB granulomas^44^. In contrast, cluster 3 was present in both necrotic and non-necrotic granulomas in our data. We therefore sought independent evidence that this ME represented a true hypoxic tissue compartment. Hypoxia was scored across all spatial spots using Seurat’s AddModuleScore function and UCell with canonical HIF-responsive genes^45,46^. Both methods identified Cluster 3 as the most hypoxia-enriched ME with strong global concordance between the methods (Spearman ρ = 0.77, p < 0.001; Supplementary Fig. 5A-B). AddModuleScore produced a more spatially restricted pattern that better matched granuloma architecture and was therefore used for subsequent analyses (Supplementary Fig. 5D-E).

Spots were further classified as hypoxia-high or hypoxia-low using a hybrid per-granuloma thresholding strategy (Methods). Cluster 3 was enriched more than fourfold in hypoxia-high regions compared to hypoxia-low regions (odds ratio= 5.70, 95% CI= [5.16–6.31], adjusted p < 0.001, Supplementary Table 3), This enrichment remained after accounting for patient and granuloma in a mixed-effects logistic model (odds ratio = 7.24, 95% CI= [6.48–8.09]). Spatial mapping showed that hypoxia-high regions formed discrete domains across granulomas from all four patients (Fig. 4B-C).

### SPP1 is the dominant immune-associated transcript of the hypoxia-associated granuloma compartment

Differential expression between hypoxia-high and hypoxia-low spots was performed using MAST with patient identity as a latent variable^47^.The top overexpressed genes in the hypoxia-high regions were dominated by canonical HIF-1α responsive genes such as *SLC2A1*, *VEGFA, SLC16A3, ENO1, PGAM1* and *PFKB3 (*Supplementary Table 4)^48^. Within this validated program, (*SPP1*ranked third overall by significance and second by fold change (log₂FC = 1.55, adjusted p = 2.99 × 10⁻³⁰²) making it the highest-ranked immune-associated transcript enriched in hypoxia-high regions (Fig. 4D). *SPP1* has been implicated in macrophage activation, tissue remodeling, CD44-dependent signaling, and immunoregulatory myeloid states across multiple disease contexts ^49–52^.

To examine *SPP1* distribution, spots were classified as *SPP1*-high or SPP1-low using the same thresholding strategy applied for hypoxia (Methods). *SPP1*-high spots were enriched in hypoxia-high regions compared to hypoxia-low regions (36.2% vs 12.3%). This corresponds to hypoxia-high spots showing 4.04-fold greater odds of SPP1-high classification (odds ratio = 4.04, 95% CI = [3.69–4.42], Fisher exact test p < 0.001; Supplementary Table 5; Fig. 4E). Spatial mapping confirmed that *SPP1*-high regions co-localized with hypoxia-high regions across granulomas and patients (Fig. 4C & F). The 63.8% of hypoxia-high spots that remained *SPP1*-low indicates that *SPP1* is enriched in a subset of hypoxic regions rather than uniformly across all regions. This suggests that *SPP1* enrichment likely depends on local cell composition and cell-state context in addition to hypoxia.

Per-granuloma analysis showed a positive correlation between continuous hypoxia and *SPP1* gene expression in most granulomas (Fig. 4G; Supplementary Table 6). To account for the nested structure of the data, we fitted a linear mixed-effects model with patient and granulomas as random effects. Hypoxia score remained positively associated with *SPP1* expression (β = 2.381, 95% CI = [2.30 - 2.47], t = 55.25, p < 0.001). Hypoxia score alone explained 9.6% of *SPP1* variance (marginal R² = 0.096), with the full model accounting for 65.0% of total variance (conditional R² = 0.650). Patient identity contributed the largest share of variance beyond hypoxia (Supplementary Note 2).

Because hypoxia and inflammatory pathways such as NF-κB signaling are known to interact extensively and NF-κB signaling pathays was also enriched in cluster 3. We tested whether the Hypoxia-SPP1 association in our data could be explained by NF-κB-associated transcriptional activity. We scored all spots using 12 canonical NF-κB target genes using the same method used for hypoxia (Supplementary Methods), Although both programmes were enriched in cluster 3 (Fig. 4A, Supplementary Fig. 6A), NF-κB-associated activity was weakly correlated with hypoxia (median per-granuloma Spearman ρ = 0.14; adjusted p < 0.05 in 13 of 60 granulomas) and with SPP1 expression (median ρ = 0.15; adjusted p < 0.05 in 14 of 60 granulomas) (Supplementary Fig. 6B-C).To determine whether hypoxia-SPP1 association persist after accounting for NF-κB-associated activity, We fitted a linear mixed-effects model with SPP1 expression as the outcome, hypoxia and NF-κB scores as fixed effects, and patient and granuloma as random intercepts. Hypoxia remain strongly associated with SPP1 expression (β = 2.31, 95% CI [2.23, 2.40], t = 53.43, p < 0.001), it is only 3% lower than in the model without NF-κB (β = 2.38). NF-κB-associated activity was also positive associated SPP1 (β = 0.67, 95% CI [0.61, 0.75], t = 17.92, p < 0.001), but this association was reduced by approximately one-third after adjusting for hypoxia (β = 0.43, 95% CI [0.36, 0.50], t = 12.48, p < 0.001) (Supplementary Fig. 6D). When both predictors were Standardized, the association of hypoxia with SPP1 was approximately 4.5-fold stronger than that of NF-κB (Hypoxia B= 0.32 vs 0.07). This clearly shows the hypoxia-SPP1 association is not explained by NF-κB-associated Inflammation, although the two programmes co-occur in some granuloma regions.

Comparing hypoxia-high versus hypoxia-low regions revealed two contrasting immune programs running in parallel with the same hypoxic compartment (Fig. 5A). Hypoxia-high regions were enriched for pro-inflammatory and antimicrobial pathways. These included IFN-y signaling, antigen cross-presentation, ROS production, TNF-, IL-12-, IL-1- and NF-κB signaling. The same regions were also enriched for regulatory pathways such as IL-10-, IL-4-, IL-13 signaling. Importantly, immunosuppressive genes such as *IDO1*, *WARS*, and *VSIG*4 were highly expressed in *SPP1*-high, hypoxia-high regions (Supplementary Table 3).

**Fig 5.**
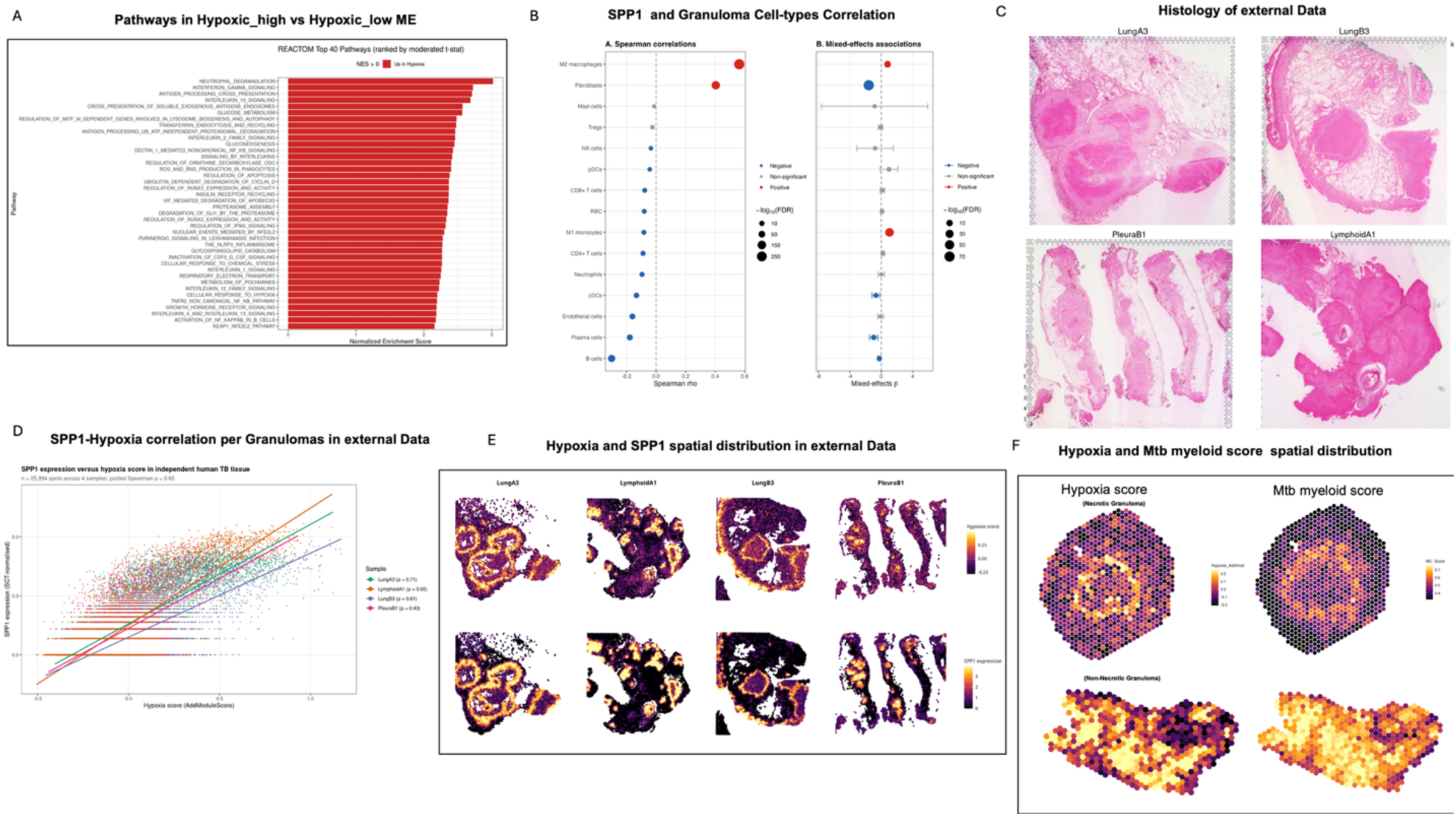
SPP1 is associated with monocyte-derived macrophage-rich regions and the hypoxia– SPP1 relationship replicates in independent human TB tissue. **(A)** Reactome pathway enrichment analysis comparing hypoxia-high and hypoxia-low regions. The top 40 pathways ranked by normalised enrichment score (NES) are shown. **(B)**Correlations between SPP1 expression and deconvolved cell-type proportions in hypoxia-high spots (n = 3,394). Left, Spearman ρ; right, fixed-effect estimates (β) from linear mixed-effects models with patient and granuloma as nested random intercepts. Error bars represent 95% confidence intervals. Point size indicates −log₁₀(FDR). **(C)** H&E images of four independent human TB samples (external dataset). **(D)** Relationship between hypoxia score (AddModuleScore) and SPP1 expression across 25,994 spatial spots from the external cohort. Each colour represents one sample. Lines indicate per-sample linear regression; Spearman correlation coefficients are shown in the legend. **(E)** Spatial distribution of hypoxia score (top) and SPP1 expression (bottom) across the four external TB samples. Colour scales are shared within each row. **(F)** Spatial Spatial distribution of hypoxia score (left) and MC macrophage signature score (right) in representative necrotic and non-necrotic granulomas. Colour scales are shared within each column.

### SPP1 expression in hypoxia-associated granuloma regions is associated with monocyte-derived macrophage-rich spots

Having established that *SPP1* expression is enriched in hypoxia-associated granuloma regions, we next sought to identify which deconvolved cell populations were mostly associated with its expression. Spot-level Spearman correlations showed that M2-like macrophages were the strongest correlate of *SPP1* expression within hypoxia-high regions (ρ = 0.563, adjusted p < 0.001), followed by fibroblasts (ρ = 0.404, adjusted p < 0.001). A similar pattern was observed in hypoxia-low regions (Fig. 5B).

To account for the nested structure of the data, we repeated the analysis using linear mixed-effects models with patient and granuloma as random effects. Within hypoxia-high spots, inflammatory monocytes (M1-like macrophages) were the strongest positive predictor of *SPP1* expression (β = 1.04, t= 13.1, 95% C.I = [0.89 – 1.20], adjusted p < 0.001), followed by M2-like macrophages (β = 0.798, t= 7.11, 95% C.I = [0.58 – 1.01], adjusted p < 0.001). In contrast, fibroblast showed a negative association under the mixed-effects model (β = −1.59, t = −18.5, 95% C.I = [−1.76 −1.42], adjusted p < 0.001). This reversal indicates that the positive Spearman correlation reflected between-patient covariation rather than within-tissue co-localization^53^. A multivariate mixed-effects model including all deconvolved cell types simultaneously, showed substantial collinearity, limiting cell-type discrimination.

*SPP1* expression in pediatric mediastinal lymph node TB granulomas is therefore most strongly associated with monocyte-derived macrophage-rich spots, including both inflammatory monocytes (M1-like) and M2-like macrophages. This resonates with the literature describing SPP1+ monocyte-derived macrophages across inflammatory, tissue-remodeling, and immunoregulatory states in chronic disease contexts ^27,54,55^.

### The hypoxia–SPP1 association replicates in an independent human TB cohort across pulmonary, pleural, and lymphoid tissue

To determine whether the hypoxia-SPP1 relationship extended beyond the pediatric cohort, we analyzed a published adult human TB spatial transcriptomic dataset ^23^. Four samples were randomly selected from this publicly available dataset, to span the context of major TB tissue types. This included two pulmonary, one pleural, and one lymphoid sample comprising 25,994 spatial spots in total. Hypoxia score and *SPP1* expression were quantified using the same methods applied to the primary in-house dataset.

Hypoxia score positively correlated with *SPP1* expression across all the adult TB samples included. Individual sample Spearman correlation ranged from ρ = 0.431 to 0.705 (all FDR-adjusted p < 0.05) and the pooled correlation across all spots was ρ = 0.648 (Fig. 5D). Spatial mapping showed consistent co-localization of hypoxia-associated regions and *SPP1* expression across all tissue types (Fig. 5C & E).

Linear mixed-effects modelling with sample as a random effect further confirmed hypoxia score as a strong positive predictor of *SPP1* expression (β = 4.26, 95% C.I = [4.22 - 4.31], t=179.1, p < 0.001). Hypoxia explained 54.6% of the variation observed in *SPP1* expression (marginal R² = 0.546), while sample identity contributed relatively little additional variance (conditional R² = 0.581). These findings thus indicate that hypoxia–*SPP1* association is conserved across independent human TB cohorts, and anatomical sites.

### The hypoxia–SPP1 niche aligns with a persistence-associated macrophage signature

To ask whether the hypoxia-SPP1 niche converges with macrophage states linked to bacterial persistence, we applied a signature derived from MC macrophages in murine lymph nodes. MC cells are a monocyte-derived macrophage population identified in murine mediastinal lymph nodes that expanded during persistent infection and accumulated intracellular bacilli over time^56^. This MC macrophage signature was used as a proxy for a persistence-associated myeloid state in our data as direct Mtb quantification in our tissue was not feasible due to the paucibacillary nature of TB pediatric tissue and discordance between the Mtb detection approaches we used in this study (Supplementary Fig. 7 Supplementary Note 3). Of the 47 genes comprising the MC macrophage signature, 46 were detected in our human dataset. MC scores were applied to all spots using AddModuleScore.

Our correlation analyses demonstrated that the hypoxia score (developed from our in-house pediatric granuloma dataset) and the MC score were positively correlated across spots (Spearman ρ = 0.557 in hypoxia-high spots, ρ = 0.405 in hypoxia-low spots; both p < 0.001). A linear mixed-effects model with patient and granuloma as nested random intercepts confirmed hypoxia score as a positive predictor of MC score (β = 0.228, 95% C.I = [0.22 – 0.24], t= 36.16, p < 0.001). To determine whether this association reflected macrophage abundance alone, M2-like macrophage and inflammatory monocyte proportions were included as covariates. Hypoxia remained independently associated with MC score (β = 0.114, 95% C.I = [0.10 – 0.12], t=21.43, p < 0.001), although the effect size was reduced. This indicated that part of the association was explained by macrophage composition but not entirely.

Finally, hypoxia score and *SPP1* expression were modelled together while adjusting for macrophage abundance. Both remained independent positive predictors of MC score (hypoxia β = 0.087, 95% C.I = [0.076-0.098], t=15.31, *SPP1* β = 0.013, 95% C.I = [0.011 - 0.015], t=13.04; both p < 0.001). To visualize the joint contribution, spots were classified by hypoxia and SPP1 status. Mean MC scores rose monotonically from the reference category (Hyp-low/SPP1-low, MC = 0.539) to the joint positive category (Hyp-high/SPP1-high, MC = 0.658). A mixed-effects model with macrophage proportions as covariates confirmed that the joint coefficient (β = 0.045, 95% C.I = [0.041 - 0.05], t = 18.69, p < 0.001) approximated the sum of the two single-positive coefficients (Hyp-high/SPP1-low β = 0.019; Hyp-low/SPP1-high β = 0.023). This result demonstrates that the contribution of hypoxia and *SPP1* to MC signature enrichment is additive rather than synergistic (Supplementary Note 4). Spatial mapping showed co-localization of hypoxia-SPP1 regions and MC scores across the granulomas (Fig. 5F).

Given that the macrophage population from which this signature was derived expanded and accumulated intracellular Mtb during murine infection, the enrichment of this signature within the pediatric hypoxia–SPP1 niche suggests that this microenvironment resembles a macrophage state previously linked to bacterial persistence. While our data do not directly demonstrate persistence in human tissue, they suggest that hypoxia-associated, SPP1-rich macrophage niches may contribute to Mtb persistence in pediatric lymph node TB.

### Multiplex immunofluorescences validate hypoxia-associated SPP1 macrophage gene program at the protein level

To determine whether the hypoxia-associated SPP1 niche identified by spatial transcriptomics is present at the protein level, we performed multiplex immunofluorescence on pediatric TB granulomas (Fig. 6). Since tissue sections from the sample originally selected for spatial transcriptomics were not available, we performed immunofluorescence on granulomas from a different patient within the broader pediatric sample cohort. Our panel included antibodies to detect SPP1 and SPP1-expressing myeloid cells; CD64 as a marker of pro-inflammatory macrophage activation; CD163 which is associated with an alternatively activated phenotype; and IDO1 as an immunoregulatory macrophage marker. Because tissue oxygenation could not be measured directly in archival FFPE material, we selected established downstream markers as a proxy of hypoxia. Our panel included proteins for glucose transporter member 1 (GLUT1); carbonic anhydrase IX (CA9); and solute carrier family 16 (SLC16A3). We also attempted to stain for hypoxia inducible factor-1α (HIF-1α), but the antibody did not perform reliably and was therefore excluded from quantification.

**Fig 6.**
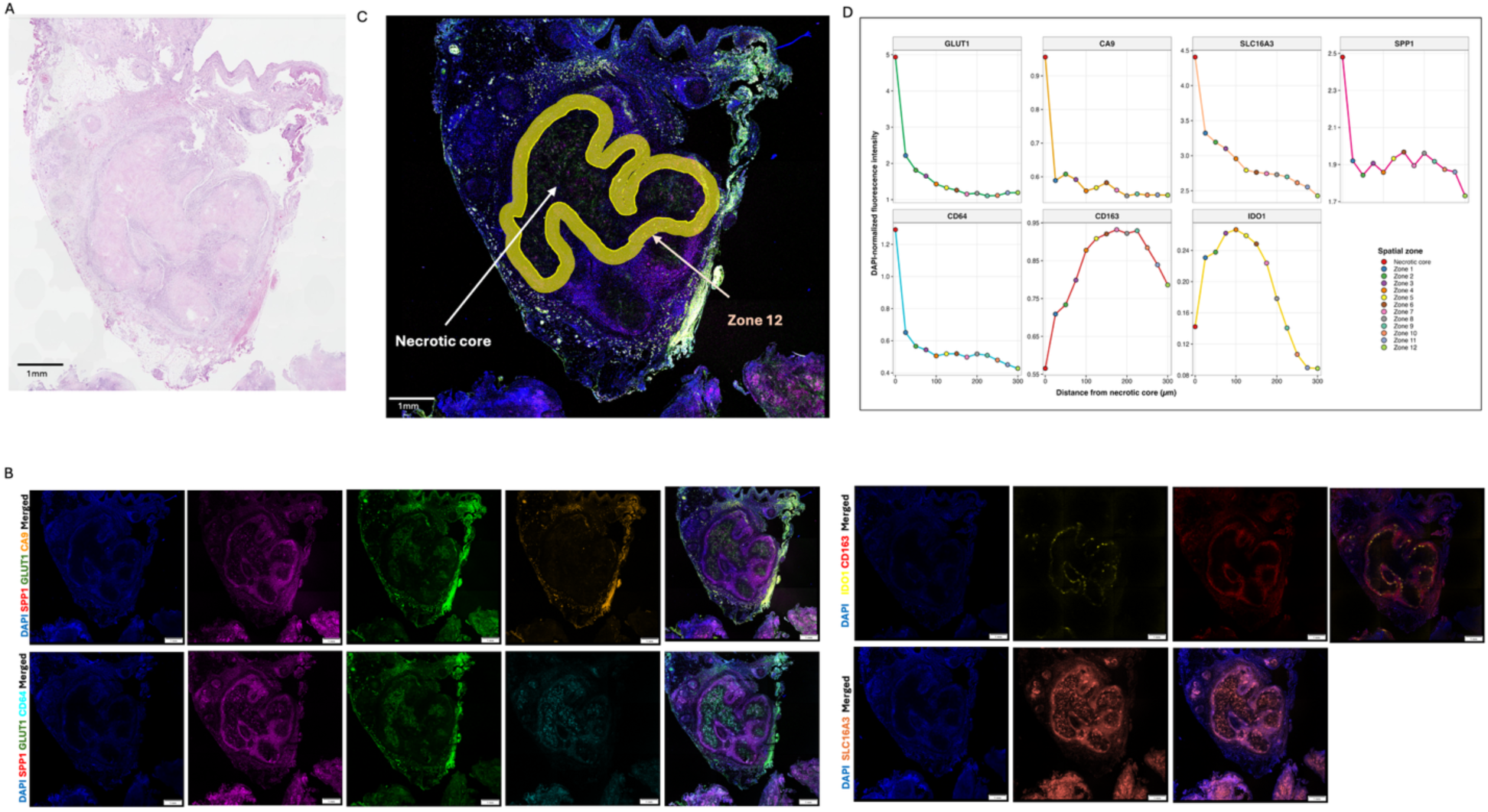
Spatial protein validation of a hypoxia –associated SPP1 macrophage compartment in a pediatric TB granuloma. **(A)** H&E staining of the granuloma used for this validation. **(B)** Multiplex immunofluorescence performed on serial sections from the same granuloma using four staining panels: DAPI, SPP1, GLUT1 and CA9 (top left); DAPI, SPP1, GLUT1 and CD64 (bottom left); DAPI, IDO1 and CD163 (top right); and DAPI and SLC16A3 (bottom right). Individual channels and the corresponding merged images are shown. GLUT1, CA9 and CD64 are enriched within the necrotic core, whereas CD163 and IDO1 are largely confined to a peripheral rim. SPP1 and SLC16A3 are detected across both compartments. **(C)**Representative merged immunofluorescence image illustrating the spatial quantification strategy. The necrotic core was outlined, followed by 12 successive 25-µm zones extending 300 µm outward from the core. The white arrows indicate the necrotic core, and the peach arrow indicates to the outermost quantified zone (Zone 12); yellow contours indicate the spatial zones used for quantification. **(D)** DAPI-normalized fluorescence intensity of GLUT1, CA9, SLC16A3, SPP1, CD64, CD163 and IDO1 across the necrotic core and successive spatial zones. Distance 0 represents the necrotic core, and colored points denote the corresponding zones shown in the spatial-zone key. Lines connect successive measurements to illustrate spatial trends. Data are from the representative tissue section shown; y-axis scales are independently adjusted for each marker. GLUT1, CA9, SLC16A3, and SPP1 are highest at or near the core and decline with distance. CD64 shows the same pattern. CD163 and IDO1 instead peak at an intermediate distance from the core before declining toward the periphery.

Tissues were stained, imaged and quantified using QuPath. For each granuloma, we annotated the (necrotic) core and generated concentric distance bands extending outward from the core at a fixed increment. Mean fluorescence intensity was measured for each marker within each band. The mean intensity was then corrected for background using tissue-free regions on the same section and normalized to DAPI intensity within the same band to account for differences in cell density across tissue region (Fig. 6C).

GLUT1, CA9, SLC16A3 and SPP1 intensity were all highest per cell within the necrotic core and declined with increasing distance from it (Fig. 6D top). SLC16A3 and SPP1 were detectable throughout the granuloma rather than restricted to necrotic region, but their strongest signal after DAPI normalization remained within the necrotic core (Fig. 6B). This pattern is similar to what we observed in the spatial transcriptomics data where regions with highest hypoxia expression, also display the highest *SPP1* expression.

The expression of the macrophage markers revealed two spatially distinct compartments. CD64 followed the same core-restricted distribution as the hypoxia markers, whereas CD163 and IDO1 were largely confined to a peripheral rim, with IDO1 occupying a narrower band than CD163 (Fig. 6B, D). SPP1 was detected in regions marked by both CD64 and CD163, suggesting that SPP1 expressing macrophages are not confined to a single, discretely polarized phenotype. They may instead represent a distinct state that does not map distinctly into the conventional M1-M2 framework. This is in line with what other studies have shown in other fields and supports the need to revisit and reclassify this distinct macrophage subset^57,58^.

## Discussion

Pediatric TB has been extensively characterized clinically and to some extent, histologically^59–61^. However, the molecular and spatial aspect of pediatric TB granulomas remain largely unexplored. This represents an important gap as much of our current mechanistic understanding of granuloma biology has been informed by experimental animal models, which often recapitulate early or primary tuberculosis^62,63^. In contrast, human tissue-based molecular work has focused more on adult pulmonary tuberculosis, which often reflects a different stage of infection and disease evolution^18,22,25,64^. It has therefore remained unclear whether the tissue-level features described in animal models apply to human primary TB disease and how the pediatric primary disease relates to better-studied adult post-primary disease. By constructing the first transcriptional map of pediatric TB granulomas, our study provides a molecular framework for examining this early disease state. Our data revealed that pediatric granulomas are organized around a limited number of recurrent transcriptional microenvironments. One of the macrophage-rich microenvironments is characterized by hypoxia and enriched for SPP1, at both gene and protein level, across different lesion states. This microenvironment is also conserved in the adult TB granuloma cohort and converges with the Mtb-permissive macrophage signature from murine lymph nodes.

A major finding of this study is that one of the recurrent microenvironments, cluster 3, which showed the strongest hypoxia-associated transcriptional activity, was present in both necrotic and non-necrotic lesions. Hypoxia has been repeatedly implicated in TB pathogenesis at lesional level^65,66^. It can result from high local metabolic demand, impaired vascular perfusion, and progressive tissue destruction that disrupt local oxygen delivery^67–71^. In animal experimental models, it has often been highlighted in the context of caseation and necrosis, where central necrosis is thought to be both a consequence and driver of oxygen deprivation^44^. In humans, direct measurement of hypoxia has reported pulmonary lesions to be severely hypoxic, although these studies did not systematically compare across necrotic and non-necrotic granulomas^72^. Our data extends this literature by showing that hypoxia-associated transcriptional programs are features of both necrotic and non-necrotic pediatric TB granulomas. This suggests that hypoxia may represent a broader feature of the human TB granuloma microenvironment rather than a downstream consequence of necrosis alone. This finding in human granulomas is in line with animal studies that have reported the stabilization of HIF-1α, reduced vascularization and activation of hypoxia-responsive pathways in granulomas, well before overt necrosis develops^73,74^.

Comparison of the pathway activity between the hypoxia-high (termed hypoxic region going forward) and hypoxia-low regions in our data reveals concurrent inflammatory and regulatory/suppressive programs within the hypoxic region. The pro-inflammatory and antimicrobial arm of this hypoxic region is consistent with prior evidence that HIF-1Α, the master regulator of hypoxia response, together with IFN-y, enhances Mtb-killing capacity of phagocytes^75,76^ through mechanisms such as nitric oxide production and induction of IL-1a, IL-1B, and IL-6^43,77,78^. In contrast, several prior studies have also described hypoxic regions as immunosuppressive niches linked to lymphocyte exclusion and bacterial persistence^56,66^. In myeloid cells specifically, hypoxia has been shown to result in the induction, recruitment, and accumulation of myeloid-derived suppressor cells (MDSC)^79–81^. While proteomic mapping of human TB granulomas shows that these opposing pro and anti-inflammatory signatures often occupy distinct compartments, with proinflammatory, antimicrobial signals at the centre and anti-inflammatory signature sin the surrounding caseum^18,65^, the co-enrichment of both programs within our hypoxic data suggest a more complex dynamic.. This finding could indicate that our Visium spots capture a mixture of neighboring inflammatory and suppressive cells, or alternatively, represent a single coordinated program in which antimicrobial activity is maintained in the region while counter-regulatory suppressive mechanism constrains it to limit collateral tissue damage. Such a response may be sufficient to restrict bacillary expansion but insufficient to achieve a sterilizing clearance in a granuloma. A further, non-exclusive explanation is that granuloma maturity contributes to the signature observed in our data. HIF-1α has been shown to play opposing roles across the course of murine TB infection. It is host protective in early infection through macrophage activation alongside IFN-y and detrimental later through inhibition of apoptosis in foamy macrophages^82^. Our cohort likely spans granulomas of varying maturity. It is possible that the co-enrichment we observe partly reflects an aggregate signal across granulomas at different stages of this trajectory rather than a single uniform state within each one. Ultimately, while our cross-sectional, spot-level data cannot fully disentangle these biologically plausible interpretations, our findings argue against the common literature simplification of viewing hypoxic regions as either purely pro-inflammatory or anti-inflammatory

Within the hypoxic regions in our study, *SPP1* emerged as the dominant immune transcript. Its cellular source in our data traced to both Inflammatory M1 and M2 macrophages. *SPP1* encodes osteopontin (OPN), an immunomodulatory protein with established roles in macrophage polarization, tissue remodeling, T cell regulation, and immune suppressive/regulatory macrophage biology^57,83,84^. Across cancer, fibrosis and chronic infection contexts, osteopontin has been variably associated with pro-inflammation, pro-fibrosis and immunosuppressive function depending on the context^49,85–87^. Recently, a SPP1+ macrophage population co-expressing VSIG4, CSF1R, CD163 and IL6R, were shown to cause inhibition of CD8 T cell responses and tumor metastases^88^. SPP1 secreted from tumor-associated macrophages, has been shown to bind directly to CD44 on T cells, directly inducing T cell exhaustion and dampening anti-tumor immunity^89^. This protein has also been well investigated in the TB context^52,90–92^. Recent spatial studies have consistently identified SPP1+ macrophages as a dominant feature of TB granulomas^23,26,27^. However, the microenvironment supporting these cells is unclear. By integrating our dataset with a publicly available spatial transcriptomic dataset^23^, our data refine this picture by showing that *SPP1* expression is highly enriched in hypoxic microenvironment within granulomas. This finding aligns with experimental studies in cancer and other diseases showing that reduced oxygen tension directly induces *SPP1* expression in macrophages^93–95^.

An unexpected finding was that *SPP1* expression in our data correlates with both M1 and M2-like macrophages in our data. At first glance, this appears inconsistent with the commonly proposed immunosuppressive role of *SPP1* in TB. However, the macrophage populations identified in our analysis do not represent classical textbook M1 and M2 subsets^96^ (Supplementary Fig. 1A). The M1 population retains features of inflammatory monocytes, whereas the M2 population is enriched for tissue resident, lipid associated markers. Both populations therefore likely represent macrophage states shaped by the granuloma microenvironment, rather than terminally polarized phenotypes. This interpretation is supported by previous studies showing that both monocyte-derived and tissue-resident macrophage can express SPP1 under inflammatory conditions^97–99^. Broadly speaking, increasing evidence suggests that the M1/M2 classification does not adequately capture macrophage diversity *in vivo*^100,101^. It is therefore possible that both populations identified here belong to a broader SPP1-associated macrophage program instead of being different functional states.

Despite the strong association between hypoxia and *SPP1* expression, after accounting for patient and sample effects, hypoxia only explained ∼10-55% of variation in *SPP1* expression in our data. This indicates that hypoxia is important, but not an exclusive regulator of *SPP1* expression in TB granulomas. Studies in cancer have shown that lactate can induce *SPP1* expression through histone acetylation, representing a mechanism downstream of hypoxia but distinct from direct HIF-1α transcriptional regulation^102^. Additional metabolic or inflammatory cues also likely contribute. Identifying these signals and determining whether they influence the functional transition of *SPP1*-expressing macrophages will require mechanistic work and single-cell approaches.

This study has several limitations that should be considered when interpreting the results. First, the cohort is small, containing tissue from only 4 patients. Although this yielded 60 granulomas for analysis, patient-level statistical power remains limited. This reflects the limited availability of pediatric TB tissue, which has been a persistent challenge for molecular studies in this field. In addition, all samples were collected after initiation of TB treatment and represent a single timepoint in the disease progression. We were therefore unable to track how hypoxic microenvironments or SPP1+ macrophages evolve during granuloma maturation or treatment. The paucibacillary nature of the samples further prevented us from directly assessing the relationship between local bacterial burden and our findings.

Technical considerations also influence interpretation of our data. The spatial transcriptomic platform used (Visium, 10x Genomics) does not provide single-cell resolution and cellular composition was therefore estimated by computational deconvolution. For this step, we used a NHP TB mediastinal lymph node reference as no comparable pediatric human single-cell dataset was available at the time of analysis. Although this reference closely matches the tissue and disease context, future studies using human pediatric scRNA-seq data or alternative single-cell spatial transcriptomics platforms would further refine cell-state annotation.

Finally, our conclusions are based primarily on transcriptomic inference. Functional studies are needed to establish the mechanistic contribution of the hypoxia-associated pathways and *SPP1*-expressing macrophages identified here. Future work incorporating richer clinical metadata, longitudinal sampling, single-cell spatial platforms and protein-level validation will further improve the findings in this study and provide a more complete understanding of granuloma heterogeneity and bacterial persistence.

In summary, our findings position the hypoxia-associated SPP1-enriched macrophage microenvironment as a recurrent feature of TB granuloma organization that is not restricted to necrotic lesions. The coexistence of antimicrobial and regulatory programs within these niches, together with its overlap with a murine Mtb persistence macrophage signature, raises the hypothesis that it represents a state of partial immune control in which bacillary expansion is constrained without clearance. Future studies combining cell-resolved spatial profiling with direct localization of viable Mtb and functional perturbation of hypoxia-SPP1 signaling will be required to test this model. Determining whether this microenvironment actively promotes Mtb persistence or arises as a response to it, could reveal features associated with poorly sterilizing lesions and inform spatially targeted host-directed therapies designed to improve bacterial clearance without exacerbating tissue damage.

## Methods

### Ethics statement

All procedures were approved by the Stellenbosch University Health Research Ethics Committee (N23/01/005), the Stellenbosch Research Ethics Committee: Biological and Environmental Safety (BES-2023-28005) and National Health Laboratory Service (NHLS)(reference number: PR2563211). All data was pseudonymized and the study was conducted in compliance with relevant institutional and national ethical regulations governing the use of archived human tissue.

### Sample selection and study design

Four archived formalin-fixed paraffin-embedded (FFPE) mediastinal lymph node tissue blocks were obtained from the National Health Laboratory Service (NHLS). Tissue was collected during routine surgical management of pediatric TB patients with airway compression caused by enlarged mediastinal lymph nodes and subsequently archived. Patients aged 4 to 36 months at the time of surgical resection were HIV-negative, and all samples were culture- and GeneXpert-positive for Mtb. Full clinical and archival details of each block are provided in Supplementary Table 1. Blocks were selected following microscopic review of diagnostic H&E slides, prioritizing those containing multiple well-defined granulomas with minimal necrosis. RNA integrity was assessed prior to library preparation using the RNeasy FFPE kit (Qiagen). Sample size reflected the availability of suitable archival tissue and the cost of Visium Spatial Gene Expression profiling.

### Tissue sectioning, histology, and granuloma annotation

Each block was serially sectioned at 5 µm thickness. Following exposure of tissue, sections one and twenty-one of each block were reserved for spatial transcriptomics. Remaining sections were allocated to a repeating Hematoxylin and Eosin (H&E), Ziehl–Neelsen (ZN), and immunofluorescence (IF) staining cycle for characterization of adjacent tissue. H&E and ZN sections were processed by standard protocols, digitized at 40× magnification on a Grundium Ocus 40 whole-slide scanner (0.25 µm/pixel). The H&E was reviewed and annotated by a qualified pathologist (S.G.). Granulomas were annotated on the spatial transcriptomics sections using QuPath v0.6.0^103^.

### Spatial transcriptomic profiling and sequencing

Spatial transcriptomic profiling was performed using the 10x Genomics Visium Spatial Gene Expression for FFPE (v1 chemistry) workflow according to the manufacturer’s instructions. FFPE sections were mounted, stained, imaged, and processed through the Visium FFPE protocol including probe hybridization, ligation, and library amplification. Libraries were converted for MGI sequencing and sequenced on the DNBSEQ-G400RS platform at the SAMRC Genomics Platform, South Africa. Detailed library preparation, QC metrics, kit specifications, and sequencing parameters are provided in Supplementary Methods.

### Data processing, integration, and quality control

FASTQ files were aligned to the human reference genome (GRCh38_A_2020) using Space Ranger v3.0.0 (10x Genomics). Count matrices were subset to pathologist-annotated granuloma regions using QuPath and Loupe Browser exports. Sixty-five granulomas were identified across 8 tissue sections, of which 60 passed quality control and were retained for downstream analyses. One section from patient_1 was excluded due to tissue lifting. Following quality control, 14,226 spatial spots remained for analysis. Data were normalized using SCTransform v2 and integrated across patients and sections using reciprocal PCA (rcPCA), which was used for all downstream analyses^104^.

### Spatial clustering and marker gene identification

Spatial neighbors were identified using FindNeighbors, and graph-based clustering was performed with FindClusters (resolution = 0.6), yielding nine reproducible transcriptional microenvironments (MEs). Cluster marker genes were identified using FindAllMarkers with the MAST test^47^.

### Cell-type deconvolution

Spot-level cell-type composition was estimated using RCTD v2.2.1 in full mode, which assigns fractional proportions of multiple cell types to each spot^38^. A non-human primate (NHP) lymph node single-cell RNA-seq dataset (GSE200151) was used as the reference, elected because of its relevance to the tissue type and disease context^17^. The NHP reference macrophage population was sub-clustered into inflammatory monocytes (M1-like) and tissue-resident lipid-associated macrophages (M2-like) prior to deconvolution (Supplementary Methods). Fifteen cell-type populations were quantified per spot: B cells, CD4+ T cells, CD8+ T cells, conventional dendritic cells (cDCs), endothelial cells, fibroblasts, inflammatory monocytes (M1-like), M2-like macrophages, mast cells, neutrophils, NK cells, plasmacytoid dendritic cells (pDCs), plasma cells, red blood cells, and regulatory T cells. Per-spot proportions were used in downstream correlation analyses.

### Pathway enrichment analysis

Gene set enrichment analysis was performed using fgsea v1.28.0 against the MSigDB Hallmark gene set collection (v2023.2)^105–107^. Ranked gene lists were generated from cluster marker genes, and pathways with FDR < 0.05 were considered significant.

### Hypoxia scoring and classification

Hypoxia module activity per spot was quantified using Seurat’s AddModuleScore with a canonical HIF-responsive gene set. UCell was used as a secondary scoring method for concordance verification. Spots were classified as hypoxia-high or hypoxia-low using a hybrid per-granuloma thresholding strategy that combined a within-granuloma 75th percentile with a global expression floor yielding 3,394 hypoxia-high spots (23.9%) and 10,832 hypoxia-low spots (76.1%) across the cohort (Supplementary Method). Hypoxia-low denotes spots that did not meet the hypoxia-high transcriptional threshold and does not imply normoxia.

### SPP1 quantification and classification

SPP1 expression was extracted from the SCT-normalized data. Spots were classified as SPP1-high or SPP1-low using a hybrid per-granuloma thresholding strategy similar to the hypoxia classification (Supplementary Methods).

### Differential expression analysis

Differential gene expression between hypoxia-high and hypoxia-low spots was performed using MAST v1.28.0 with patient identity as a latent variable to account for inter-individual variation (Supplementary Methods for full parameters).

### Hypoxia–SPP1 association analyses

The spatial and transcriptional relationship between hypoxia and *SPP1* expression was assessed using: Fisher’s exact test for spot-level enrichment; per-granuloma Spearman correlations with FDR correction; and linear mixed-effects models with SPP1 expression as outcome, hypoxia score as fixed effect, and patient and granuloma as nested random intercepts^108^. Marginal and conditional R² were computed using the `r2()` function from the performance R package ^109^, following the method of Nakagawa and Schielzeth^110^. Cluster-level context of hypoxia-SPP1 co-occurrence was tested with Fisher’s exact test across microenvironments (Supplementary Method).

### Identification of cellular sources of SPP1 expression

To identify cell populations associated with SPP1 expression, Spearman rank correlations were computed between SPP1 expression and each of the fifteen cell-type proportions. The analysis was run across all spots and separately within hypoxia-high and hypoxia-low spots. To account for nested data structure, the analysis was repeated using linear mixed-effects models with patient and granuloma included as nested random intercepts. Both univariate (one cell-type predictor per model) and multivariate (all fourteen non-reference cell-type proportions entered simultaneously) strategies were applied. Univariate models were used to interpret cellular associations. Multivariate results are reported for methodological transparency given the inherent collinearity of compositional cell-type data.

### MC macrophage signature scoring and convergence analysis

A 47-gene MCMC macrophage signature derived from published scRNA-seq data of murine Mtb-infected lymph node tissue was applied to the integrated human dataset^56^. Mouse genes were mapped to human’s orthologues where possible, and 46 of the 47 genes were detected in the human dataset. MC module scores were calculated for each spatial spot using Seurat AddModuleScore. The relationship between MC score, hypoxia, and *SPP1* expression was evaluated using spearman correlation and linear mixed-effects models^108^. Details of the modeling strategy are provided in the Supplementary Methods.

### External validation in an independent human TB cohort

A pre-processed integrated Seurat object containing TB granuloma spatial transcriptomic data from Pyle et al 2025 was downloaded^23^. Four samples were used for external validation: two pulmonary TB samples (LungA3, LungB3), one pleural TB sample (PleuraB1), and one lymphoid TB sample (LymphoidA1), comprising 25,994 spatial spots in total. To ensure direct comparability with the primary cohort analysis, hypoxia and SPP1 quantification in this dataset was done with the same methods used in the primary study. AddModuleScore with canonical hypoxia gene marker, and SPP1 expression extracted from the SCT-normalized data slot. The hypoxia–SPP1 relationship was assessed using per-sample Spearman correlations with Benjamini–Hochberg FDR correction, pooled Spearman correlation across all spots, and linear mixed-effects modelling with sample identity as a single random intercept.

### Immunofluorescence

Immunofluorescence (IF) staining was performed on FFPE sections to validate the spatial patterns of hypoxia-associated, and SPP1 protein expression observed in the spatial transcriptomics data. Sections were baked at 60 °C for 1 hour, deparaffinized in xylene, rehydrated through a graded ethanol series, and subjected to antigen retrieval by heating in citrate buffer (pH 6.0) or Tris-EDTA (pH 8.0) at 95 °C for 30 minutes. Autofluorescence was quenched with Sudan Black B, and sections were permeabilized with 0.2% Triton X-100 for intracellular targets prior to blocking in PBS + 5% BSA + 0.05% Tween-20. Primary antibodies were applied overnight at 4 °C and detected using species-specific Alexa Fluor-conjugated secondary. Nuclei were counterstained with DAPI, and slides mounted in fluorescence mounting medium. Two panels were used: a hypoxia panel (CA9, GLUT1/SLC2A1, SLC16A3, HIF1A, SPP1/Osteopontin) and a myeloid panel (IDO1, CD64, CD163). Full antibody details with catalogue numbers and dilutions are provided in Supplementary Table 7. Following staining, tissue sections were imaged using an Olympus (Evident) FLUOVIEW FV4000 confocal microscope (FLUOVIEW software v4.1.1.134). Each granuloma was then quantified using Qupath(v0.6.0). Details on the quantification approach is in supplementary method.

## Data availability

Raw and processed spatial transcriptomics data generated in this study will be deposited at NCBI Gene Expression Omnibus (GEO) and the accession number will be made available to the public after publication. The external validation spatial dataset and murine dataset are publicly available at <u>GSE296400</u> and <u>GSE309019</u> respectively. Raw and processed data for multiplex immunofluorescence is in IF_supplenmentary_file.

## Code availability

Custom scripts used to perform the analyses and generate the figures are available to editors and reviewers upon request. The final code will be made publicly available through GitHub and archived in Zenodo upon publication.

## Acknowledgements

We thank Reggie Williams, Jeffery Pietersen and Vusi Ntywankile from the Clinical Anatomy Laboratory, Stellenbosch University, for assistance with tissue sectioning, histological staining, and imaging. We thank Lize Engelbrecht of the Stellenbosch University Central Analytical Facilities for assistance with immunofluorescence of image acquisition. We thank Sophete Gers for assistance with tissue annotation and G.W. McElfresh, for expert advice during the initial computational analysis. We thank the South African Medical Research Council (SAMRC) Genomics Platform for providing sequencing resources and infrastructure. This work was supported by the Cascade IMPAc-TB study funded by the National Institute of Allergy and Infectious Diseases (NIAID), NIH, through contract #75N93019C0070 (issued to K.B.U. at Seattle Children’s Research Institute), sub-award issued to G.W. at Stellenbosch University, and also the HI-IMPAcTB study funded by the NIAID, NIH, through contract#75N9301900071 (issued to S.F at Harvard T.H. Chan School of Public Health), sub-award issued to G.W at Stellenbosch University.

## Author Contributions

S.O.A.: Conceptualization, Methodology, Investigation, Formal analysis, Data curation, Visualization, Validation, Writing – original draft, Writing – review & editing, P.G.: Investigation, Resources, Writing – review & editing, D.D.W.: Investigation, Resources, Writing – review & editing. J.T.J: Resources, Writing – review & editing, T.J.: Investigation, Methodology, Supervision, Writing – review & editing. A.S.: Investigation, Writing – review & editing, A.D.: Writing – review & editing. S.B.: Investigation, Writing-review & editing. M.N.: Investigation, Writing – review & editing, M.C.C: Writing – review & editing, S.F.: Writing – review & editing, K.B.S.: Investigation, Writing – review & editing, J.D.E: Investigation, Writing – review & editing, E.S.: Writing – review & editing, K.U.: Writing – review & editing, M.G.: Writing – review & editing, G.W.: Conceptualization, Methodology, Funding acquisition, Supervision, Writing – review & editing, N.D.P.: Conceptualization, Methodology, Funding acquisition, Supervision, Writing – review & editing. All authors reviewed and approved the final manuscript.

## Supplementary Figures

Supplementary Methods, Notes and Tables are provided in a separate file

**Supplementary Fig. 1.**
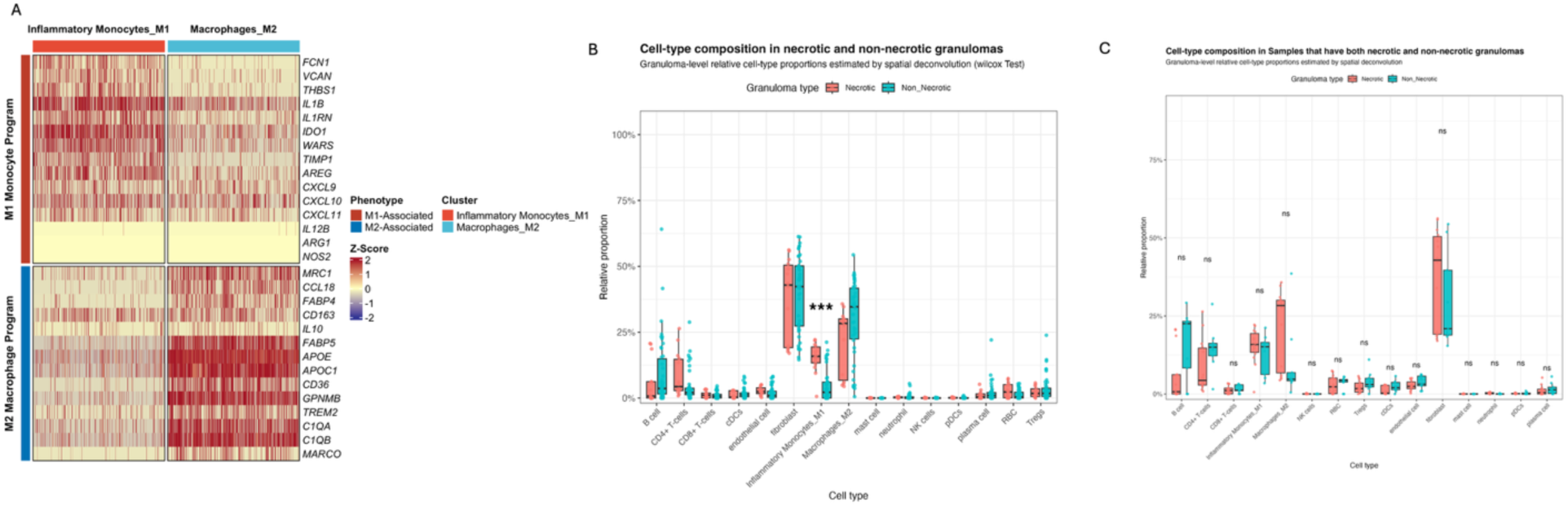
Annotation of macrophage subpopulations in the NHP reference and sensitivity analysis of granuloma cell composition. **(A)** Heatmap showing z-score-normalised expression of selected marker genes across the two macrophage subpopulations identified in the non-human primate (NHP) lymph node single-cell RNA-seq reference used for RCTD deconvolution. Each column represents one cell. Z-scores were calculated for each gene across all cells. (**B**) Granuloma-level cell-type proportions across necrotic (n = 14, salmon) and non-necrotic (n = 46, teal) granulomas. Each point represents one granuloma. Box plots show median and interquartile range. Significance reflects wilcox test (*** FDR < 0.0001). **(C)** Granuloma-level relative cell-type proportions in patients containing both necrotic and non-necrotic granulomas. Each point represents one granuloma; box plots show the median and interquartile range. Statistical significance was assessed using the Wilcoxon rank-sum test with Benjamini–Hochberg. No cell type remained significant after multiple-testing correction.

**Supplementary Fig. 2.**
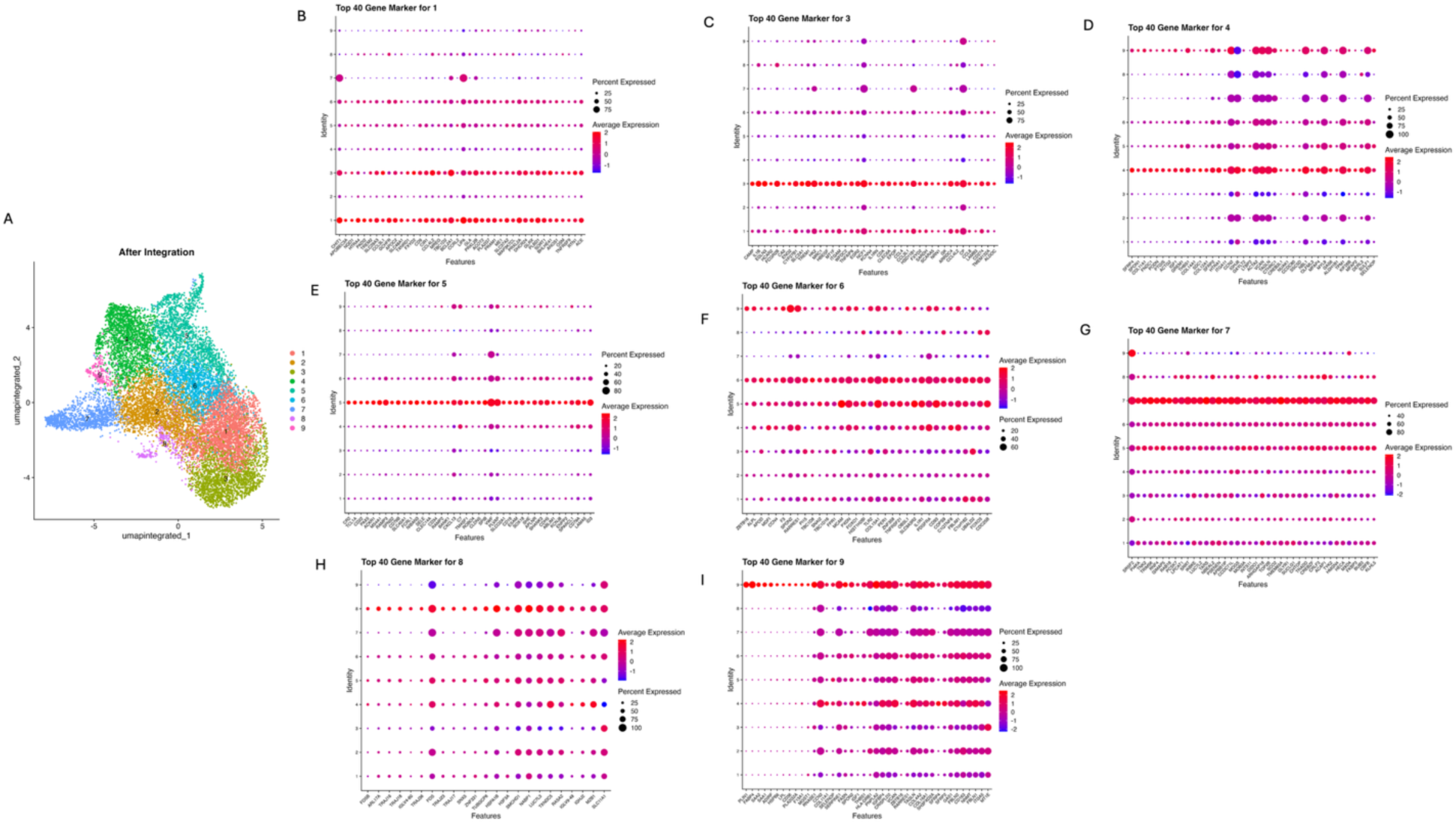
Marker gene expression defining the nine transcriptional microenvironments. **(A)** Integrated UMAP of all 14,226 spatial spots following reciprocal PCA (rcPCA) integration, coloured by cluster assignment. **(B–I)** Dot plots showing the top 40 marker genes for each microenvironment identified using the MAST test. Dot size indicates the percentage of spots expressing each gene, and colour indicates the scaled average expression. Marker genes were used together with RCTD deconvolution results (Supplementary Fig. 3) to assign the biological identities of the nine microenvironments described in Supplementary Note 1.

**Supplementary Fig. 3.**
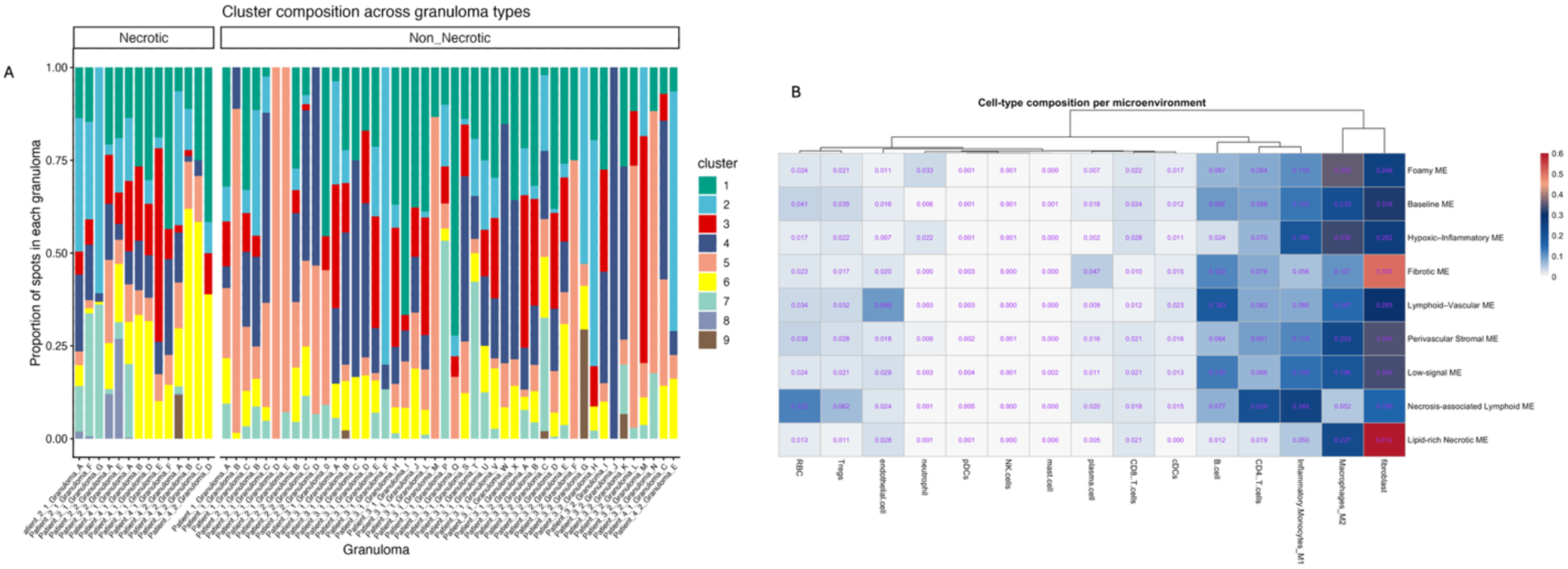
Distribution of microenvironments and their inferred cellular composition. **(A)** Relative abundance of the nine transcriptional microenvironments within each granuloma. Each stacked bar represents one granuloma, with colours indicating the proportion of spots assigned to each microenvironment. Granulomas are grouped by histological classification (necrotic and non-necrotic). **(B)** Mean deconvolved cell-type composition of each microenvironment estimated by RCTD using the non-human primate lymph node single-cell RNA-seq reference. Values represent the average estimated cell-type proportion within each microenvironment. Hierarchical clustering was performed on the cell-type dimension.

**Supplementary Fig. 4.**
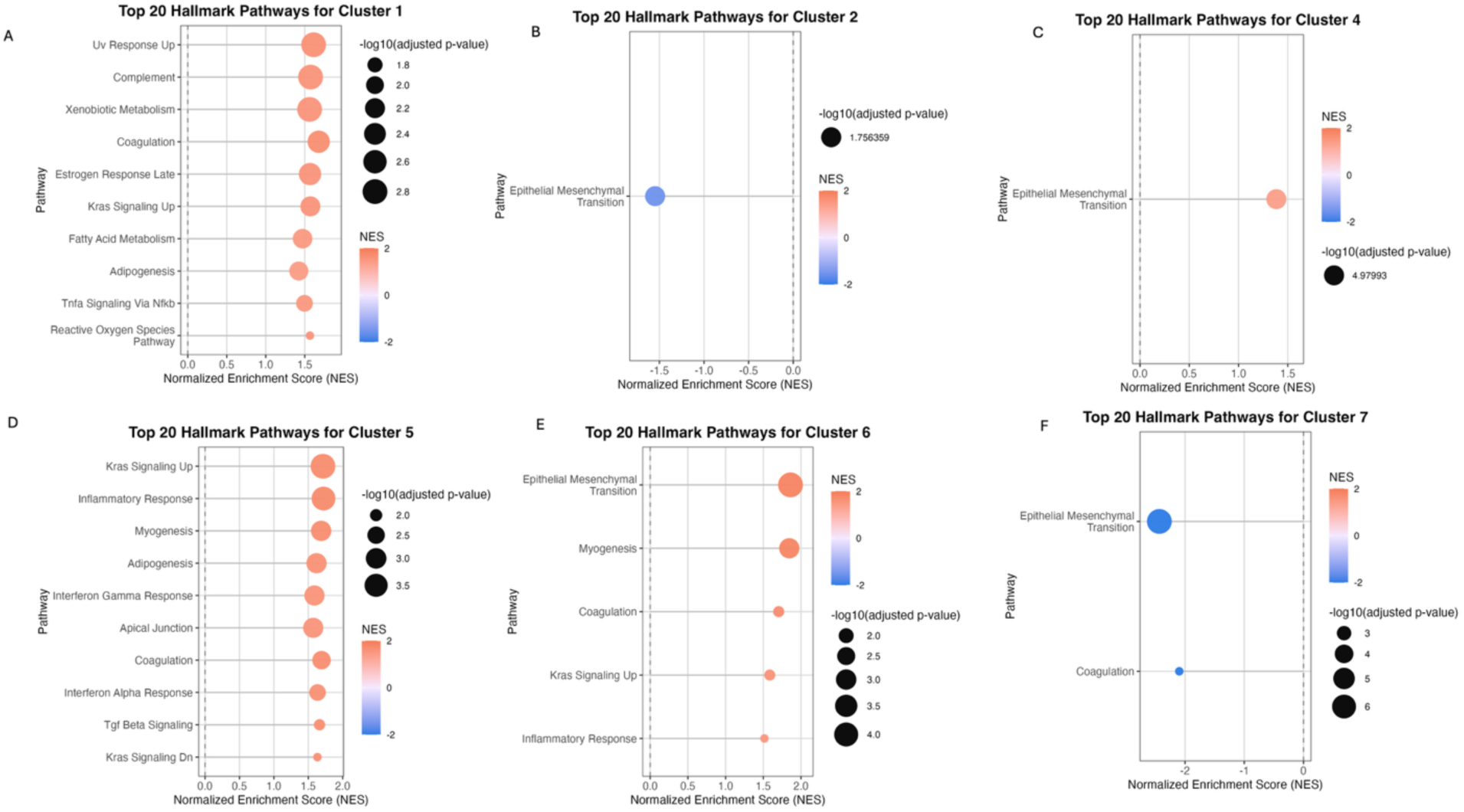
Hallmark pathway enrichment across the remaining transcriptional microenvironments. Gene set enrichment analysis (GSEA) of marker genes from the remaining microenvironments against the MSigDB Hallmark gene set collection. **(A)** Foamy ME (Cluster 1). **(B)** Baseline ME (Cluster 2). **(C)** Fibrotic ME (Cluster 4). **(D)** Lymphoid–Vascular ME (Cluster 5). **(E)** Perivascular Stromal ME (Cluster 6). **(F)** Low-signal ME (Cluster 7). The x-axis shows the normalized enrichment score (NES). Dot size represents −log10(adjusted *P* value), and dot colour represents the normalized enrichment score. Only significantly enriched pathways (FDR < 0.05) are shown. Cluster 3 is presented in the main text (Fig. 4A). Clusters 8 and 9 did not yield significant Hallmark pathway enrichment after multiple-testing correction.

**Supplementary Fig. 5.**
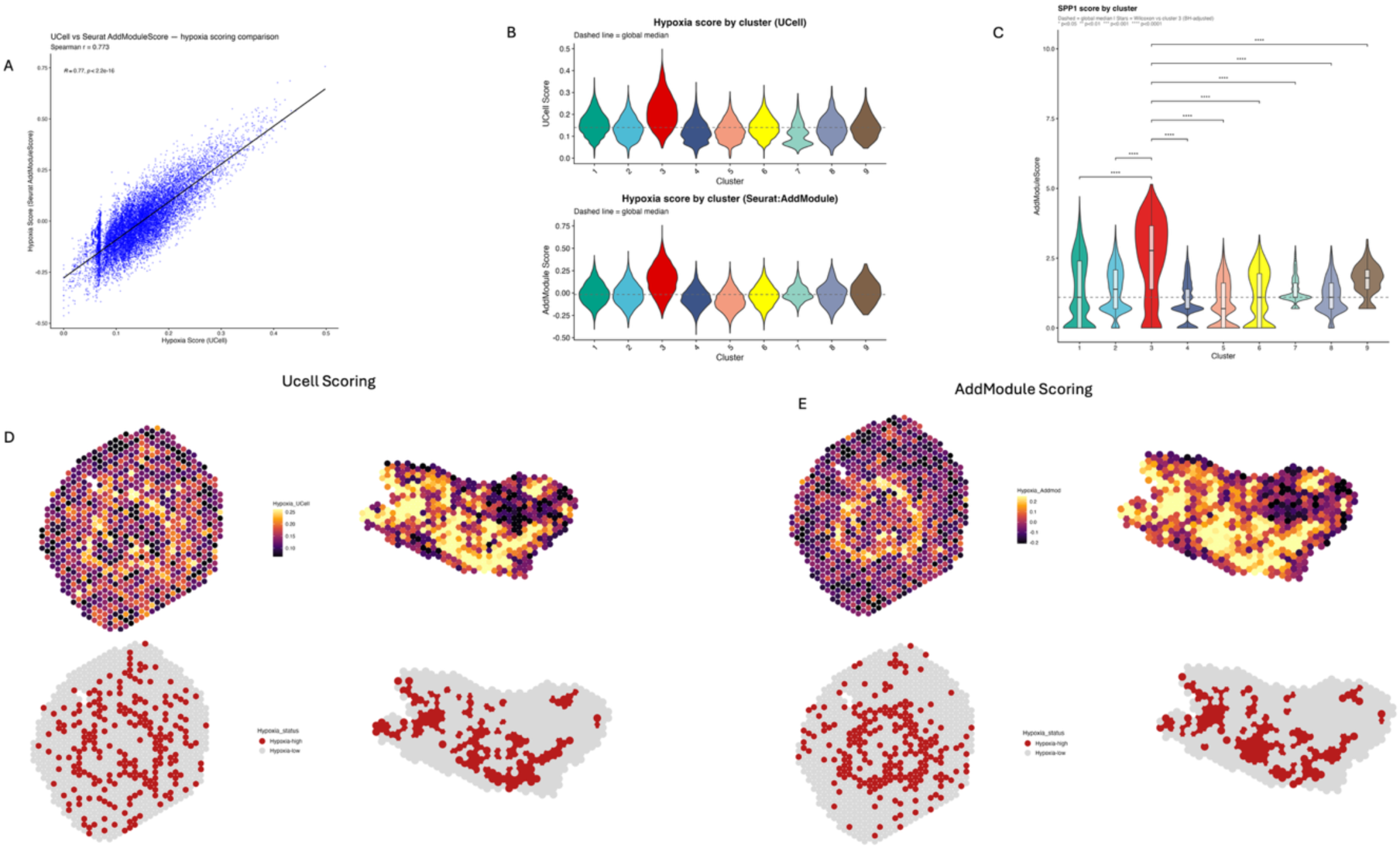
Comparison of hypoxia scoring methods and validation of hypoxia classification. **(A)** Correlation between hypoxia scores generated using UCell and Seurat AddModuleScore across all 14,226 spatial spots (Spearman ρ = 0.77, *P* < 0.001). **(B)** Distribution of hypoxia scores across the nine transcriptional microenvironments calculated using UCell (top) and AddModuleScore (bottom). Dashed lines indicate the global median score for each method. **(C)** Distribution of SPP1 expression across the nine microenvironments. Boxplots within violins show the median and interquartile range. Statistical significance was assessed using pairwise Wilcoxon rank-sum tests with Benjamini–Hochberg correction. **(D)** Spatial distribution of hypoxia scores generated using UCell (top) and the corresponding binary hypoxia classification (bottom) in representative necrotic and non-necrotic granulomas. **(E)** Spatial distribution of hypoxia scores generated using AddModuleScore (top) and the corresponding binary hypoxia classification (bottom) in the same representative granulomas. AddModuleScore produced more spatially restricted hypoxia domains that better matched granuloma architecture and was therefore used for all downstream analyses.

**Supplementary Fig. 6.**
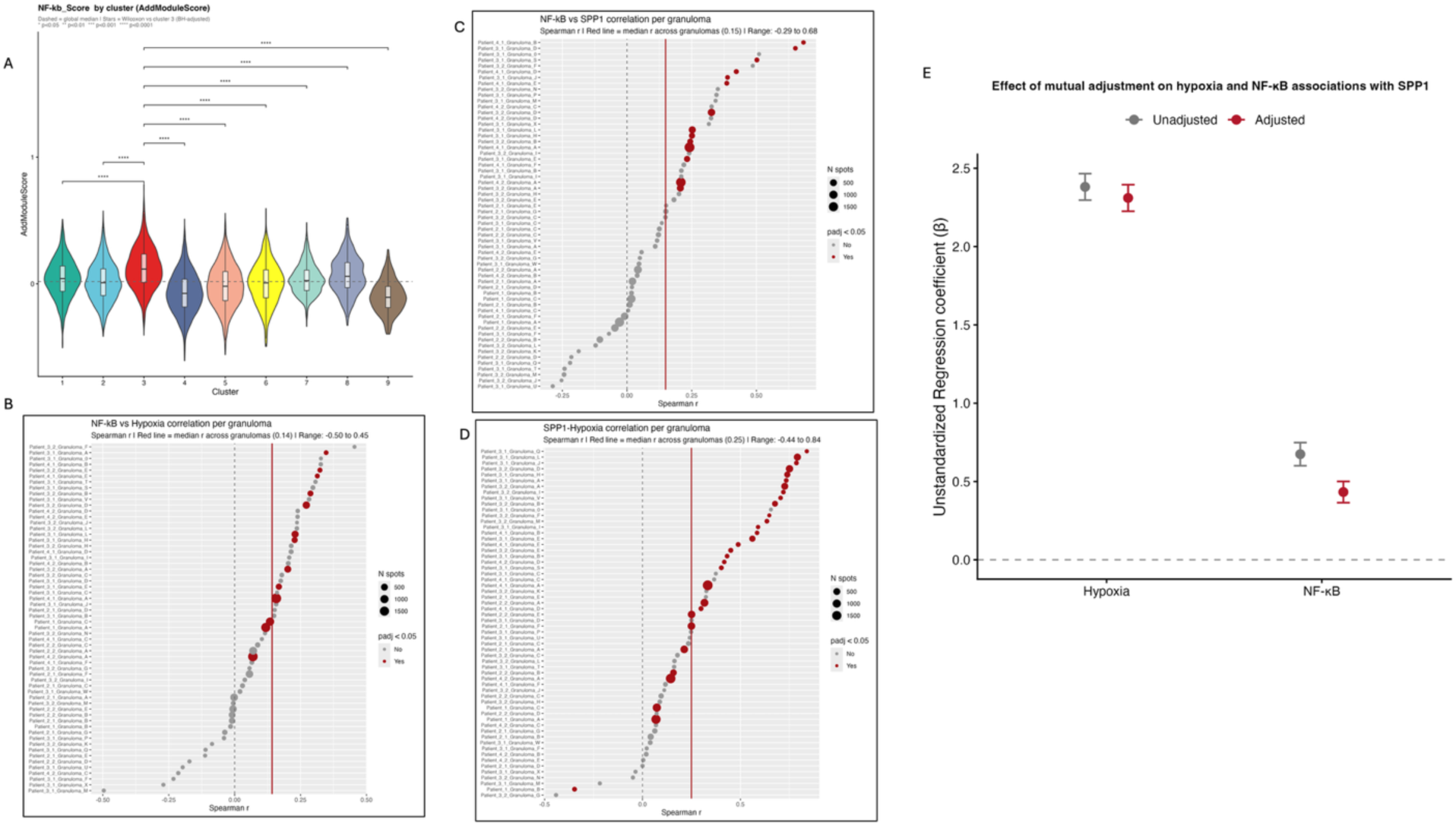
Relationship between NF-κB-associated activity, hypoxia and SPP1 expression across granulomas. **(A)** Distribution of NF-κB expression across the nine microenvironments. Boxplots within violins show the median and interquartile range. Statistical significance was assessed using pairwise Wilcoxon rank-sum tests with Benjamini–Hochberg correction. **(B-D)** Per-granuloma Spearman correlations across 60 granulomas. Each point represents one granuloma; point size indicates the number of spots. Red points indicate correlations significant after Benjamini–Hochberg correction (adjusted P < 0.05), grey points are not significant. The dashed line marks ρ = 0 and the red line the median ρ across granulomas. (**B)** NF-κB score versus hypoxia score (median ρ = 0.14, significant in 13 granulomas). **(C)** NF-κB score versus SPP1 expression (median ρ = 0.15, significant in 14 granulomas). **(D)** Hypoxia score versus SPP1 expression (median ρ = 0.25, significant in 31 granulomas, 30 positively correlated). **(E)** Coefficients from linear mixed-effects models of SPP1 expression, with hypoxia and NF-κB scores fitted alone (unadjusted) or together (adjusted). Points show unstandardised regression coefficients (β) and error bars 95% confidence intervals. All models included patient and granuloma as random intercepts. Adjustment for NF-κB-associated activity reduced the hypoxia coefficient by approximately 3% (β = 2.38 to 2.31), whereas adjustment for hypoxia reduced the NF-κB coefficient by approximately 36% (β = 0.67 to 0.43).

**Supplementary Fig. 7.**
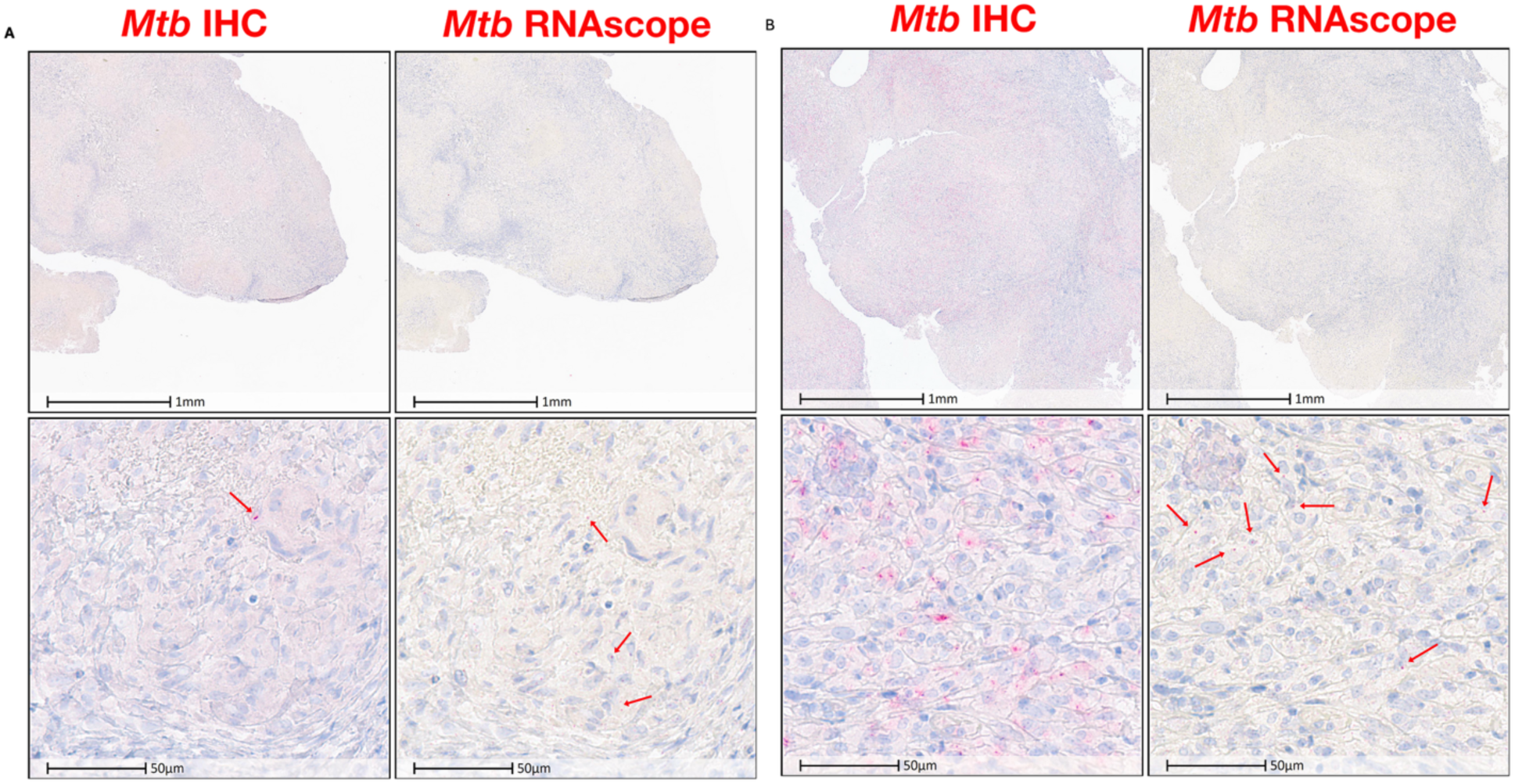
Detection of Mtb in Pediatric TB Granuloma tissue. Mtb was detected in across Lymph nodes samples from the pediatric cohort using IHC, with anti-PPD antibody and RNAscope. The upper panels show whole-section overview, The lower panel show high-magnification images (scale bar=50um). Left panels show IHC and right panels show RNAscope for each sample **(A)** A sample representative of most of the cohort showing very low Mtb signal by both IHC and RNAcope. Red arrows indicate detected Mtb antigen and RNA detected. **(B)** The only sample in the cohort that show high Mtb signal by both methods. Some discordance was observed between the IHC and RNAscope across samples.

## References

1. World Health Organization: Roadmap towards ending…- Google Scholar. Accessed July 21, 2026. https://scholar.google.com/scholar_lookup?title=Roadmap%20Towards%20Ending%20Tb%20in%20Children%20and%20Adolescents&publication_year=2023&

2. Marais BJ, Gie RP, Schaaf HS, et al. The natural history of childhood intra-thoracic tuberculosis: a critical review of literature from the pre-chemotherapy era. Int J Tuberc Lung Dis Off J Int Union Tuberc Lung Dis. 2004;8(4):392–402.

3. Huynh J, Abo YN, du Preez K, Solomons R, Dooley KE, Seddon JA. Tuberculous Meningitis in Children: Reducing the Burden of Death and Disability. Pathogens. 2021;11(1):38. doi:10.3390/pathogens11010038

4. Yerramsetti S, Cohen T, Atun R, Menzies NA. Global estimates of paediatric tuberculosis incidence in 2013–19: a mathematical modelling analysis. Lancet Glob Health. 2022;10(2):e207–e215. doi:10.1016/S2214-109X(21)00462-9

5. Seddon JA, Chiang SS, Esmail H, Coussens AK. The Wonder Years: What Can Primary School Children Teach Us About Immunity to Mycobacterium tuberculosis? Front Immunol. 2018;9. doi:10.3389/fimmu.2018.02946

6. Mehrian P, Moghaddam AM, Tavakkol E, et al. Determining the lymphadenopathy characteristics of the mediastinum in lung CT scan of children with tuberculosis. Int J Mycobacteriology. 2016;5(3):306–312. doi:10.1016/j.ijmyco.2016.06.015

7. Chen Q, Huang T, Zou L, et al. Differences in epidemiological and clinical features between adult and pediatric tracheobronchial tuberculosis patients in Southwest China. Front Public Health. 2023;11. doi:10.3389/fpubh.2023.1225267

8. Martinez L, Cords O, Liu Q, et al. Infant BCG vaccination and risk of pulmonary and extrapulmonary tuberculosis throughout the life course: a systematic review and individual participant data meta-analysis. Lancet Glob Health. 2022;10(9):e1307–e1316. doi:10.1016/S2214-109X(22)00283-2

9. Mangtani P, Abubakar I, Ariti C, et al. Protection by BCG Vaccine Against Tuberculosis: A Systematic Review of Randomized Controlled Trials. Clin Infect Dis. 2014;58(4):470–480. doi:10.1093/cid/cit790

10. Basile FW, Nabeta P, Ruhwald M, Song R. Pediatric Tuberculosis Diagnostics: Present and Future. J Pediatr Infect Dis Soc. 2022;11(Supplement_3):S85–S93. doi:10.1093/jpids/piac082

11. Ernst JD. The immunological life cycle of tuberculosis. Nat Rev Immunol. 2012;12(8):581–591. doi:10.1038/nri3259

12. Gideon HP, Phuah J, Myers AJ, et al. Variability in Tuberculosis Granuloma T Cell Responses Exists, but a Balance of Pro-and Anti-inflammatory Cytokines Is Associated with Sterilization. PLOS Pathog. 2015;11(1):e1004603. doi:10.1371/journal.ppat.1004603

13. Cronan MR. In the Thick of It: Formation of the Tuberculous Granuloma and Its Effects on Host and Therapeutic Responses. Front Immunol. 2022;13. doi:10.3389/fimmu.2022.820134

14. Pagán AJ, Ramakrishnan L. The Formation and Function of Granulomas. Annu Rev Immunol. 2018;36(Volume 36, 2018):639–665. doi:10.1146/annurev-immunol-032712-100022

15. Ramakrishnan L. Revisiting the role of the granuloma in tuberculosis. Nat Rev Immunol. 2012;12(5):352–366. doi:10.1038/nri3211

16. Lin PL, Ford CB, Coleman MT, et al. Sterilization of granulomas is common in active and latent tuberculosis despite within-host variability in bacterial killing. Nat Med. 2014;20(1):75–79. doi:10.1038/nm.3412

17. Gideon HP, Hughes TK, Tzouanas CN, et al. Multimodal profiling of lung granulomas in macaques reveals cellular correlates of tuberculosis control. Immunity. 2022;55(5):827–846.e10. doi:10.1016/j.immuni.2022.04.004

18. Marakalala MJ, Raju RM, Sharma K, et al. Inflammatory signaling in human tuberculosis granulomas is spatially organized. Nat Med. 2016;22(5):531–538. doi:10.1038/nm.4073

19. Subbian S, Tsenova L, Kim MJ, et al. Lesion-Specific Immune Response in Granulomas of Patients with Pulmonary Tuberculosis: A Pilot Study. PLoS ONE. 2015;10(7):e0132249. doi:10.1371/journal.pone.0132249

20. Abengozar-Muela M, Esparza MV, Garcia-Ros D, et al. Diverse immune environments in human lung tuberculosis granulomas assessed by quantitative multiplexed immunofluorescence. Mod Pathol Off J U S Can Acad Pathol Inc. 2020;33(12):2507–2519. doi:10.1038/s41379-020-0600-6

21. Cadena AM, Fortune SM, Flynn JL. Heterogeneity in tuberculosis. Nat Rev Immunol. 2017;17(11):691–702. doi:10.1038/nri.2017.69

22. McCaffrey EF, Donato M, Keren L, et al. The immunoregulatory landscape of human tuberculosis granulomas. Nat Immunol. 2022;23(2):318–329. doi:10.1038/s41590-021-01121-x

23. Pyle CJ, Wang L, Beerman RW, et al. Paired single-cell and spatial transcriptional profiling reveals a central osteopontin macrophage response mediating tuberculous granuloma formation. mBio. 16(9):e01559–25. doi:10.1128/mbio.01559-25

24. Gern BH, Klas JM, Foster KA, et al. Early and opposing neutrophil and CD4 T cell responses shape pulmonary tuberculosis pathology. J Exp Med. 2025;222(10):e20250161. doi:10.1084/jem.20250161

25. Qiu X, Zhong P, Yue L, et al. Spatial transcriptomic sequencing reveals immune microenvironment features of Mycobacterium tuberculosis granulomas in lung and omentum. Theranostics. 2024;14(16):6185–6201. doi:10.7150/thno.99038

26. Mbano IM, Liu N, Wadsworth MH, et al. Single-cell and spatial profiling highlights TB-induced myofibroblasts as drivers of lung pathology. J Exp Med. 2026;223(3):e20251067. doi:10.1084/jem.20251067

27. Katano H, Hebisawa A, Sato Y, Hoshino Y. Identification of SPP1-positive macrophages by single-cell spatial analysis in human lung tissues with mycobacterial infection. bioRxiv. Preprint posted online January 20, 2025:2024.09.12.612778. doi:10.1101/2024.09.12.612778

28. Ahmad F, Rani A, Alam A, et al. Macrophage: A Cell With Many Faces and Functions in Tuberculosis. Front Immunol. 2022;13. doi:10.3389/fimmu.2022.747799

29. Cohen SB, Gern BH, Delahaye JL, et al. Alveolar Macrophages Provide an Early Mycobacterium tuberculosis Niche and Initiate Dissemination. Cell Host Microbe. 2018;24(3):439–446.e4. doi:10.1016/j.chom.2018.08.001

30. Russell DG, Cardona PJ, Kim MJ, Allain S, Altare F. Foamy macrophages and the progression of the human TB granuloma. Nat Immunol. 2009;10(9):943–948. doi:10.1038/ni.1781

31. Chen Z, Kong X, Ma Q, et al. The impact of Mycobacterium tuberculosis on the macrophage cholesterol metabolism pathway. Front Immunol. 2024;15. doi:10.3389/fimmu.2024.1402024

32. Dabla A, Liang YC, Rajabalee N, et al. TREM2 Promotes Immune Evasion by Mycobacterium tuberculosis in Human Macrophages. mBio. 2022;13(4):e01456–22. doi:10.1128/mbio.01456-22

33. Shang X, Maimaiti N, Fan J, et al. Triggering Receptor Expressed on Myeloid Cells 2 Mediates the Involvement of M2-Type Macrophages in Pulmonary Tuberculosis Infection. J Inflamm Res. 2024;17:1919–1928. doi:10.2147/JIR.S435216

34. Luo G, Wen Y, Wang M, et al. A granulin-positive macrophage subtype in mycobacterial granulomas alleviates tissue damage by limiting excessive inflammation. Proc Natl Acad Sci U S A. 122(31):e2413946122. doi:10.1073/pnas.2413946122

35. Singh B, Sharan R, Ravichandran G, et al. Indoleamine-2,3-dioxygenase inhibition improves immunity and is safe for concurrent use with cART during Mtb/SIV coinfection. JCI Insight. 9(15):e179317. doi:10.1172/jci.insight.179317

36. Pessanha AP, Martins RAP, Mattos-Guaraldi AL, Vianna A, Moreira LO. Arginase-1 expression in granulomas of tuberculosis patients. FEMS Immunol Med Microbiol. 2012;66(2):265–268. doi:10.1111/j.1574-695X.2012.01012.x

37. SM N, AJ B, S A, E W, B K. Paediatric Tuberculosis. Lancet Infect Dis. 2008;8(8):498–510. doi:10.1016/S1473-3099(08)70182-8

38. Cable DM, Murray E, Zou LS, et al. Robust decomposition of cell type mixtures in spatial transcriptomics. Nat Biotechnol. 2022;40(4):517–526. doi:10.1038/s41587-021-00830-w

39. Zimmerman KD, Espeland MA, Langefeld CD. A practical solution to pseudoreplication bias in single-cell studies. Nat Commun. 2021;12(1):738. doi:10.1038/s41467-021-21038-1

40. Ståhl PL, Salmén F, Vickovic S, et al. Visualization and analysis of gene expression in tissue sections by spatial transcriptomics. Science. 2016;353(6294):78–82. doi:10.1126/science.aaf2403

41. Kim MJ, Wainwright HC, Locketz M, et al. Caseation of human tuberculosis granulomas correlates with elevated host lipid metabolism. EMBO Mol Med. 2010;2(7):258–274. doi:10.1002/emmm.201000079

42. Palsson-McDermott EM, Curtis AM, Goel G, et al. Pyruvate Kinase M2 regulates Hif-1α activity and IL-1β induction, and is a critical determinant of the Warburg Effect in LPS-activated macrophages. Cell Metab. 2015;21(1):65–80. doi:10.1016/j.cmet.2014.12.005

43. Tannahill GM, Curtis AM, Adamik J, et al. Succinate is an inflammatory signal that induces IL-1β through HIF-1α. Nature. 2013;496(7444):238–242. doi:10.1038/nature11986

44. Via LE, Lin PL, Ray SM, et al. Tuberculous Granulomas Are Hypoxic in Guinea Pigs, Rabbits, and Nonhuman Primates. Infect Immun. 2008;76(6):2333–2340. doi:10.1128/IAI.01515-07

45. Andreatta M, Carmona SJ. UCell: Robust and scalable single-cell gene signature scoring. Comput Struct Biotechnol J. 2021;19:3796–3798. doi:10.1016/j.csbj.2021.06.043

46. Tirosh I, Izar B, Prakadan SM, et al. Dissecting the multicellular ecosystem of metastatic melanoma by single-cell RNA-seq. Science. 2016;352(6282):189–196. doi:10.1126/science.aad0501

47. Finak G, McDavid A, Yajima M, et al. MAST: a flexible statistical framework for assessing transcriptional changes and characterizing heterogeneity in single-cell RNA sequencing data. Genome Biol. 2015;16:278. doi:10.1186/s13059-015-0844-5

48. Semenza GL. Hypoxia-Inducible Factors in Physiology and Medicine. Cell. 2012;148(3):399–408. doi:10.1016/j.cell.2012.01.021

49. Lang F, Li Y, Yao R, Jiang M. Osteopontin in Chronic Inflammatory Diseases: Mechanisms, Biomarker Potential, and Therapeutic Strategies. Biology. 2025;14(4):428. doi:10.3390/biology14040428

50. Lund SA, Giachelli CM, Scatena M. The role of osteopontin in inflammatory processes. J Cell Commun Signal. 2009;3(3):311–322. doi:10.1007/s12079-009-0068-0

51. Castello LM, Raineri D, Salmi L, et al. Osteopontin at the Crossroads of Inflammation and Tumor Progression. Mediators Inflamm. 2017;2017:4049098. doi:10.1155/2017/4049098

52. Nau GJ, Liaw L, Chupp GL, Berman JS, Hogan BLM, Young RA. Attenuated Host Resistance againstMycobacterium bovis BCG Infection in Mice Lacking Osteopontin. Infect Immun. 1999;67(8):4223–4230. doi:10.1128/iai.67.8.4223-4230.1999

53. Fisher AJ, Medaglia JD, Jeronimus BF. Lack of group-to-individual generalizability is a threat to human subjects research. Proc Natl Acad Sci. 2018;115(27):E6106–E6115. doi:10.1073/pnas.1711978115

54. Sun L, Chu X, Kong T, et al. An SPP1-SOCS1 pathway constrains interferon responses in tumor-associated macrophages and shapes an immunosuppressive tumor microenvironment. Immunity. 2026;59(5):1422–1437.e9. doi:10.1016/j.immuni.2026.04.001

55. Mantel I, Zhang H, Vargas J, et al. SPP1hi macrophages in fibrin niches promote hyperplastic tissue remodeling in rheumatoid arthritis synovium. Sci Transl Med. 2026;18(855):eaed3087. doi:10.1126/scitranslmed.aed3087

56. Shamskhou EA, Duffy FJ, Cross LM, et al. Monocytic niches escape T cell surveillance and promote *Mycobacterium tuberculosis* persistence in lymph nodes. Immunity. 2026;59(8):2122–2141.e9. doi:10.1016/j.immuni.2026.05.017

57. Bill R, Wirapati P, Messemaker M, et al. CXCL9:SPP1 macrophage polarity identifies a network of cellular programs that control human cancers. Science. 2023;381(6657):515–524. doi:10.1126/science.ade2292

58. Reggio A, Fuoco C, Deodati R, Palma A. SPP1 macrophages across diseases: A call for reclassification? FASEB J. 2025;39(5):e70448. doi:10.1096/fj.202403227R

59. Perez-Velez CM, Marais BJ. Tuberculosis in Children. N Engl J Med. 2012;367(4):348-361. doi:10.1056/NEJMra1008049

60. Lyu J, Narum DE, Baldwin SL, et al. Understanding the development of tuberculous granulomas: insights into host protection and pathogenesis, a review in humans and animals. Front Immunol. 2024;15:1427559. doi:10.3389/fimmu.2024.1427559

61. Purbaningsih W, Rachmawati M, Triyani Y, Rahmi FS. Histopathological Review of Granuloma in Diagnosis of Tuberculosis Lymphadenitis (TBL). Glob Med Health Commun GMHC. 2023;11(3):171–175. doi:10.29313/gmhc.v11i3.12742

62. Cohen SB, Gern BH, Urdahl KB. The Tuberculous Granuloma and Preexisting Immunity. Annu Rev Immunol. 2022;40(Volume 40, 2022):589–614. doi:10.1146/annurev-immunol-093019-125148

63. Hunter L, Ruedas-Torres I, Agulló-Ros I, Rayner E, Salguero FJ. Comparative pathology of experimental pulmonary tuberculosis in animal models. Front Vet Sci. 2023;10. doi:10.3389/fvets.2023.1264833

64. Sawyer AJ, Patrick E, Edwards J, et al. Spatial mapping reveals granuloma diversity and histopathological superstructure in human tuberculosis. J Exp Med. 2023;220(6):e20221392. doi:10.1084/jem.20221392

65. Prosser G, Brandenburg J, Reiling N, Barry CE, Wilkinson RJ, Wilkinson KA. The bacillary and macrophage response to hypoxia in tuberculosis and the consequences for T cell antigen recognition. Microbes Infect. 2017;19(3):177–192. doi:10.1016/j.micinf.2016.10.001

66. McCaffrey EF, Delmastro AC, Fitzhugh I, et al. The immunometabolic topography of cellular organization and bacterial control in tuberculosis granulomas. Nat Immunol. 2026;27(4):867–880. doi:10.1038/s41590-026-02431-8

67. Shi L, Salamon H, Eugenin EA, Pine R, Cooper A, Gennaro ML. Infection with Mycobacterium tuberculosis induces the Warburg effect in mouse lungs. Sci Rep. 2015;5(1):18176. doi:10.1038/srep18176

68. Oehlers SH, Cronan MR, Scott NR, et al. Interception of host angiogenic signalling limits mycobacterial growth. Nature. 2015;517(7536):612–615. doi:10.1038/nature13967

69. Datta M, Via LE, Kamoun WS, et al. Anti-vascular endothelial growth factor treatment normalizes tuberculosis granuloma vasculature and improves small molecule delivery. Proc Natl Acad Sci. 2015;112(6):1827–1832. doi:10.1073/pnas.1424563112

70. Datta M, Kennedy M, Siri S, et al. Mathematical model of oxygen, nutrient, and drug transport in tuberculosis granulomas. PLOS Comput Biol. 2024;20(2):e1011847. doi:10.1371/journal.pcbi.1011847

71. Datta M, Via LE, Dartois V, Xu L, Barry CE, Jain RK. Leveraging insights from cancer to improve tuberculosis therapy. Trends Mol Med. 2025;31(1):11–20. doi:10.1016/j.molmed.2024.07.011

72. Belton M, Brilha S, Manavaki R, et al. Hypoxia and tissue destruction in pulmonary TB. Thorax. 2016;71(12):1145–1153. doi:10.1136/thoraxjnl-2015-207402

73. Makafe GG, Low-Beer T, Travis K, et al. Mouse models of tuberculosis uniformly featuring hypoxic necrotic lesions and poor response to chemotherapy. Infect Immun. 94(3):e00574–25. doi:10.1128/iai.00574-25

74. Cardoso MS, Silva TM, Resende M, Appelberg R, Borges M. Lack of the Transcription Factor Hypoxia-Inducible Factor 1α (HIF-1α) in Macrophages Accelerates the Necrosis of Mycobacterium avium-Induced Granulomas. Infect Immun. 2015;83(9):3534–3544. doi:10.1128/IAI.00144-15

75. Zenk SF, Hauck S, Mayer D, Grieshober M, Stenger S. Stabilization of Hypoxia-Inducible Factor Promotes Antimicrobial Activity of Human Macrophages Against Mycobacterium tuberculosis. Front Immunol. 2021;12. doi:10.3389/fimmu.2021.678354

76. Braverman J, Sogi KM, Benjamin D, Nomura DK, Stanley SA. HIF-1α is an essential mediator of IFN-γ dependent immunity to Mycobacterium tuberculosis. J Immunol Baltim Md 1950. 2016;197(4):1287-1297. doi:10.4049/jimmunol.1600266

77. Braverman J, Stanley SA. Nitric oxide modulates macrophage responses to M. tuberculosis infection through activation of HIF-1α and repression of NF-kB. J Immunol Baltim Md 1950. 2017;199(5):1805-18161. doi:10.4049/jimmunol.1700515

78. Zhu T, Liu H, Su L, et al. MicroRNA-18b-5p Downregulation Favors Mycobacterium tuberculosis Clearance in Macrophages via HIF-1α by Promoting an Inflammatory Response. ACS Infect Dis. 2021;7(4):800–810. doi:10.1021/acsinfecdis.0c00650

79. Hypoxia induces myeloid-derived suppressor cell…: Hepatology. doi:10.1002/hep.28655

80. Corzo CA, Condamine T, Lu L, et al. HIF-1α regulates function and differentiation of myeloid-derived suppressor cells in the tumor microenvironment. J Exp Med. 2010;207(11):2439–2453. doi:10.1084/jem.20100587

81. Chiu DKC, Tse APW, Xu IMJ, et al. Hypoxia inducible factor HIF-1 promotes myeloid-derived suppressor cells accumulation through ENTPD2/CD39L1 in hepatocellular carcinoma. Nat Commun. 2017;8(1):517. doi:10.1038/s41467-017-00530-7

82. Baay-Guzman GJ, Duran-Padilla MA, Rangel-Santiago J, Tirado-Rodriguez B, Et. A. Dual Role of Hypoxia-Inducible Factor 1 α in Experimental Pulmonary Tuberculosis: Its Implication as a New Therapeutic Target. Future Microbiol. Published online 2018. doi:10.2217/fmb-2017-0168

83. Lund SA, Giachelli CM, Scatena M. The role of osteopontin in inflammatory processes. J Cell Commun Signal. 2009;3(3-4):311–322. doi:10.1007/s12079-009-0068-0

84. Wang KX, Denhardt DT. Osteopontin: Role in immune regulation and stress responses. Cytokine Growth Factor Rev. 2008;19(5):333–345. doi:10.1016/j.cytogfr.2008.08.001

85. Karasalih B, Duman H, Bechelany M, Karav S. Osteopontin: Its Properties, Recent Studies, and Potential Applications. Int J Mol Sci. 2025;26(12):5868. doi:10.3390/ijms26125868

86. Pardo A, Gibson K, Cisneros J, et al. Up-Regulation and Profibrotic Role of Osteopontin in Human Idiopathic Pulmonary Fibrosis. PLoS Med. 2005;2(9):e251. doi:10.1371/journal.pmed.0020251

87. Shurin MR. Osteopontin controls immunosuppression in the tumor microenvironment. J Clin Invest. 128(12):5209–5212. doi:10.1172/JCI124918

88. Trehan R, Huang P, Zhu XB, et al. SPP1 + macrophages cause exhaustion of tumor-specific T cells in liver metastases. Nat Commun. 2025;16(1):4242. doi:10.1038/s41467-025-59529-0

89. Hou L, Jiang M, Li Y, et al. Targeting SPP1+ macrophages via the SPP1-CD44 axis reveals a key mechanism of immune suppression and tumor progression in ovarian cancer. Int Immunopharmacol. 2025;160:114906. doi:10.1016/j.intimp.2025.114906

90. Koguchi Y, Kawakami K, Uezu K, et al. High Plasma Osteopontin Level and Its Relationship with Interleukin-12–mediated Type 1 T Helper Cell Response in Tuberculosis. Am J Respir Crit Care Med. 2003;167(10):1355–1359. doi:10.1164/rccm.200209-1113OC

91. Motaung B, Holtzhausen AR, Stanley K, van Rensburg I, Snyders CI, Loxton AG. The value of the monocyte-to-lymphocyte ratio and osteopontin (SPP1) in tuberculosis treatment response monitoring. Sci Rep. 2026;16(1):15390. doi:10.1038/s41598-026-46176-8

92. Van Der Windt GJW, Wieland CW, Wiersinga WJ, Florquin S, Van Der Poll T. Osteopontin is not crucial to protective immunity during murine tuberculosis. Immunology. 2009;128(1pt2):e766-e776. doi:10.1111/j.1365-2567.2009.03081.x

93. Wei J, Chen Z, Hu M, et al. Characterizing Intercellular Communication of Pan-Cancer Reveals SPP1+ Tumor-Associated Macrophage Expanded in Hypoxia and Promoting Cancer Malignancy Through Single-Cell RNA-Seq Data. Front Cell Dev Biol. 2021;9. doi:10.3389/fcell.2021.749210

94. Wang L, Chen B, He J, et al. ZFP36L1 promotes non-small cell lung cancer progression under hypoxia by modulating CXCL9:SPP1 polarity: A single-cell transcriptomic study. Clin Transl Med. 2026;16(4):e70642. doi:10.1002/ctm2.70642

95. Yang J, Li X, Zhu X, et al. Hypoxia-Induced Osteopontin-Positive Glioma-Associated Macrophages Facilitate Glioma Mesenchymal Transition via NF-κB Pathway Activation. Cancer Commun. 46:0007. doi:10.34133/cancomm.0007

96. Martinez FO, Gordon S, Locati M, Mantovani A. Transcriptional Profiling of the Human Monocyte-to-Macrophage Differentiation and Polarization: New Molecules and Patterns of Gene Expression. J Immunol. 2006;177(10):7303–7311. doi:10.4049/jimmunol.177.10.7303

97. Morse C, Tabib T, Sembrat J, et al. Proliferating SPP1/MERTK-expressing macrophages in idiopathic pulmonary fibrosis. Eur Respir J. 2019;54(2):1802441. doi:10.1183/13993003.02441-2018

98. Duan X, Zhang L, Liu K, et al. Macrophage-derived SPP1 exacerbate myocardial injury by interacting with fibroblasts in viral myocarditis. Biol Direct. 2025;20(1):30. doi:10.1186/s13062-025-00621-2

99. He C, Coarfa C, Lebimoyo O, et al. D29-10 Targeting Macrophage Secreted Phosphoprotein 1 in Pulmonary Fibrosis. Am J Respir Crit Care Med. 2026;212(Supplement_1):aamag162.2870. doi:10.1093/ajrccm/aamag162.2870

100. Murray PJ, Allen JE, Biswas SK, et al. Macrophage activation and polarization: nomenclature and experimental guidelines. Immunity. 2014;41(1):14–20. doi:10.1016/j.immuni.2014.06.008

101. Xue J, Schmidt SV, Sander J, et al. Transcriptome-Based Network Analysis Reveals a Spectrum Model of Human Macrophage Activation. Immunity. 2014;40(2):274–288. doi:10.1016/j.immuni.2014.01.006

102. Zhang J, Li Z, Yin J, et al. SPP1+ macrophages polarized by lactate confer the progression of hypoxic adaptive tumor cells in brain. Neuro-Oncol. 2026;28(1):70–85. doi:10.1093/neuonc/noaf208

103. Bankhead P, Loughrey MB, Fernández JA, et al. QuPath: Open source software for digital pathology image analysis. Sci Rep. 2017;7:16878. doi:10.1038/s41598-017-17204-5

104. Hao Y, Stuart T, Kowalski MH, et al. Dictionary learning for integrative, multimodal and scalable single-cell analysis. Nat Biotechnol. 2024;42(2):293–304. doi:10.1038/s41587-023-01767-y

105. Korotkevich G, Sukhov V, Budin N, Shpak B, Artyomov M, Sergushichev A. Fast gene set enrichment analysi. Preprint posted online 2016. doi:10.1101/060012

106. Subramanian A, Tamayo P, Mootha VK, et al. Gene set enrichment analysis: A knowledge-based approach for interpreting genome-wide expression profiles. Proc Natl Acad Sci U S A. 2005;102(43):15545–15550. doi:10.1073/pnas.0506580102

107. Liberzon A, Birger C, Thorvaldsdóttir H, Ghandi M, Mesirov JP, Tamayo P. The Molecular Signatures Database (MSigDB) hallmark gene set collection. Cell Syst. 2015;1(6):417–425. doi:10.1016/j.cels.2015.12.004

108. Bates D, Mächler M, Bolker B, Walker S. Fitting Linear Mixed-Effects Models Using lme4. J Stat Softw. 2015;67:1–48. doi:10.18637/jss.v067.i01

109. Lüdecke D, Ben Shachar M, Patil I, Waggoner P, Makowski D. performance: An R Package for Assessment, Comparison and Testing of Statistical Models. J Open Source Softw. 2021;6:3139. doi:10.21105/joss.03139

110. Nakagawa S, Schielzeth H. A general and simple method for obtaining R2 from generalized linear mixed-effects models. Methods Ecol Evol. 2013;4(2). Accessed August 6, 2026. https://pub.uni-bielefeld.de/record/2565368

