## Supplementary Methods,Note and Tables for "Spatial transcriptomics of pediatric tuberculosis lymph nodes reveals a conserved hypoxia–SPP1 niche"

#### Spatial transcriptomic library preparation and sequencing.

Each Visium capture area (6.5 × 6.5 mm, ~5,000 spatially barcoded spots at 55 µm diameter) was processed following the manufacturer's Visium FFPE User Guide (CG000407, Rev D). Sections were deparaffinized, H&E-stained using an in-house established protocol, imaged, decrosslinked, hybridized with the Visium FFPE human probe set, and processed through probe ligation, extension, and library amplification. Libraries were indexed using the Dual Index Plate TS Set A (10x Genomics) and subjected to quality control with the Qubit dsDNA High Sensitivity Assay and TapeStation High Sensitivity D5000 Assay, respectively. Linear libraries were converted using the MGIEasy Universal Library Conversion Kit (App-A, MGI) and sequenced using a custom cycling program with 28 cycles for Read 1 (spatial barcode), 100 cycles for Read 2 (cDNA insert), and 10 cycles for dual index sequencing.

#### Data processing and clustering

Spots were retained if they had at least 200 detected genes, at least 400 total UMI counts, and fewer than 30% mitochondrial reads. Genes detected in fewer than three spots across the merged dataset were excluded. SCTransform v2 was applied with default parameters, regressing out the number of detected features (nFeature_Spatial) to account for variation in spot-level RNA capture efficiency. Reciprocal PCA (rcPCA) was applied to the first 30 principal components of the SCTransform-normalized data. rcPCA-corrected embeddings were used for all downstream dimensionality reduction, clustering, and spatial analyses. Spatial neighbors were computed using FindNeighbors on rcPCA-corrected embeddings (dims = 1:30). Spots were clustered using FindClusters at resolution 0.6, yielding nine transcriptional microenvironments. FindAllMarkers was run with the MAST test, restricting to genes with $\log_2$ fold-change $\geq 0.35$, minimum detection fraction $\geq 0.30$ in the expressing cluster, and minimum difference in detection fraction $\geq 0.15$ between clusters. For each cluster, a ranked gene list was constructed from the full FindAllMarkers output run without fold-change or detection thresholds, using the signed product of $\log_2$ fold-change and $-\log_{10}$ raw p-value as the ranking metric. GSEA was run with 10,000 permutations, minimum gene set size of 15, and maximum of 500. Pathways reaching FDR $< 0.05$ were reported.

#### NHP reference sub-clustering.

The NHP lymph node reference dataset (GSE200151) originally annotated myeloid cells as a single broad "Macrophage" population. To enable finer-resolution deconvolution of Visium spots, this population was sub-clustered using Seurat FindNeighbors and FindClusters resolving two transcriptionally distinct subpopulations. The first (n = 10,008 cells; labelled "Inflammatory Monocytes_M1") showed the highest expression of monocyte recruitment markers (FCN1, VCAN, THBS1), pro-inflammatory transcripts (IL1B, IL1RN, IDO1), and the IFN-γ-induced chemokines CXCL9, CXCL10, and CXCL11, indicating IFN-γ-driven M1-skewed activation. Canonical M1 effectors (NOS2, IL12B, IL6) were minimally expressed. The second subpopulation (n = 16,100 cells; labelled "Macrophages_M2") co-expressed alternatively activated macrophage markers (MRC1, CD163, CCL18, IL10) and a tissue-resident lipid-associated macrophage signature (APOE, APOC1, C1QA/B/C, TREM2, MARCO, CD36, FABP4/5, GPNMB, PPARG) characteristic of foamy or LAM-type macrophages in chronic granulomatous inflammation including TB. This subpopulation showed lower-level CXCL9, CXCL10, and CXCL11 expression and minimal canonical M2 effectors (ARG1). Reduced NOS2 and ARG1 induction is consistent with primate compared to murine macrophages. The co-expression of polarization and IFN-γ-response programs across both subclusters aligns with reports that TB granuloma macrophages do not occupy pure M1 or M2 states but carry features of multiple activation programs. Full transcriptional profiles supporting these annotations are shown in Supplementary Fig S1. M1/M2 labels are retained in figures for traceability but should be interpreted as inflammatory monocytes and tissue-resident alternatively activated / lipid-associated macrophages, respectively.

#### Molecular scoring and statistical analyses

**Hypoxia scoring.** The canonical hypoxia gene set comprised *HIF1A, EGLN3, CA9, VEGFA, BNIP3, SLC2A1, PDK1, ENO1, SLC16A3, PFKFB3, ANGPTL4, HK2, LOX* and *PGAM1*. All fourteen genes were confirmed present in the SCT assay. AddModuleScore was run with nbin = 12. UCell v2.6.2 was applied as a secondary scoring method

AddModuleScore was selected as the primary scoring and classification method based on spatial specificity. Spatial visualization demonstrated that AddModuleScore produced hypoxia-enriched regions concentrated at anatomically consistent granuloma locations co-localizing with expected necrotic architecture, whereas UCell produced more diffuse signal extending beyond biologically expected hypoxic boundaries (Supplementary Fig. 5D–E). This difference reflects the methodological distinction that UCell ranks genes within each spot independently, such that spots with modest co-elevation of hypoxia genes receive intermediate scores regardless of absolute expression levels. AddModuleScore uses background subtraction, which subtraction penalizes spots where hypoxia gene expression is only marginally above expression-matched control genes, yielding more spatially restricted enrichment. Despite this difference in spatial behavior, the two methods showed strong global concordance (Spearman $\rho = 0.77$, $p < 2.2 \times 10^{-16}$; Cohen's $\kappa = 0.593$ applied to binary classifications), confirming that the underlying hypoxia signal was robust to methodological choice.

#### Biological rationale for the hypoxia classification threshold

The global AddModuleScore distribution across all 14,226 spots showed a median of −0.017, with 44.8% of spots scoring above zero. This distribution shape indicates that above-background hypoxia gene activity is spatially restricted to a minority of spots rather than being a universal feature of granuloma tissue. Zero was therefore selected as the primary biological floor for hypoxia-high classification, as it represents the point at which hypoxia gene expression equals that of expression-matched control genes, a threshold with intrinsic biological meaning independent of the data distribution. This contrasts with percentile-based absolute cut-offs, which would shift with the size and composition of the cohort and would not have a direct biological interpretation.

Within this absolute biological floor, a hybrid per-granuloma thresholding strategy was applied to preserve within-lesion resolution across granulomas with heterogeneous absolute hypoxia levels. For each granuloma independently, the 75th percentile of AddModuleScore values was computed.

Where the within-granuloma 75th percentile exceeded zero, it was used as the classification threshold, identifying the most hypoxically active spots within an already above-background distribution. In granulomas where the within-granuloma 75th percentile fell at or below zero, indicating that the upper quarter of spots in that lesion showed no above-background hypoxia activity, zero was applied as the absolute classification floor and only spots with positive AddModuleScore values were classified as hypoxia-high.

**SPP1 quantification and thresholding.** SPP1 expression was extracted from the SCT-normalized data slot (global median = 1.099, 80.9% of spots expressing SPP1 above zero). Spots were classified using an absolute expression floor of 1.099 normalized units (the global median of SPP1-expressing spots and the inflection point of the expression distribution) combined with a within-granuloma 75th percentile threshold. Where the within-granuloma 75th percentile exceeded the absolute floor, it served as the threshold; otherwise, the floor was applied.

**NF-κB scoring.** The canonical NF-κB gene set used comprised *NFKBIA, NFKBIE, TNFAIP3, NFKB1, NFKB2, RELB, BIRC3, TNIP1, TRAF1, BCL3, ICAM1* and *CCL20*, all were drawn from the MSigDB Hallmark TNFα signalling via NF-κB gene set. The panel included NF-κB family members, regulators of NF-κB signalling and downstream NF-κB-responsive genes. The panel was selected to represent NF-κB-associated transcriptional responses while avoiding direct gene overlap with the predefined 14-gene hypoxia signature. All 12 genes were detected in the SCT assay. Module scores were calculated using Seurat's AddModuleScore function with nbin = 12, using the same parameters as for the hypoxia score.

**Hypoxia-SPP1 association pipeline details.** Enrichment of SPP1-high spots within hypoxia-high versus hypoxia-low regions was tested using Pearson's chi-squared test with Yates' continuity correction across all 14,226 spots. Effect size was quantified using the odds ratio with 95% CI via Fisher's exact test (epitools package). Per-granuloma Spearman correlations were computed for granulomas with at least ten spots and were stratified by patient identity. Linear mixed-effects models used lme4 v1.1-35 and lmerTest v3.1-3 with maximum likelihood estimation. Marginal and conditional $R^2$ were computed using MuMIn following Nakagawa and Schielzer.

**MAST differential expression parameters.** Pre-filtering used $\log_2$ fold-change ≥ 0.25 and minimum detection fraction ≥ 0.10 in at least one group. A separate unfiltered FindMarkers call was run with logfc.threshold = 0 and min.pct = 0 to generate the ranked gene list for pathway analysis.

**MC Scoring.** To test whether the hypoxia–SPP1 compartment of pediatric mediastinal lymph node TB granulomas corresponds to a transcriptional state previously characterized in murine TB infection, a published 47-gene MC macrophage signature derived from single-cell RNA-seq of *Mtb*-infected murine lymph node tissue was applied verbatim as published (ref Collaborator paper). The MC population in the murine study represents a monocyte-derived macrophage state that expands during persistent infection and accumulates an increasing proportion of intracellular bacilli at later time points. Mouse gene symbols were converted to uppercase as a first-pass mouse-to-human orthologues approximation, intersected with the gene names present in the integrated human Seurat object, and only genes detected in the human dataset were retained.

The relationship between MC score, spatial hypoxia, and SPP1 expression was assessed using per-spot Spearman correlations and three nested linear mixed-effects models (lme4, lmerTest; maximum likelihood estimation) with patient and granuloma as nested random intercepts [1]. The basic model specified MC score as a function of hypoxia score alone:

$$MC\_Score \sim Hypoxia\_Score + (1 \mid Patient / Granuloma)$$

Because the MC signature is derived from a macrophage population and the MC score is expected to track with macrophage abundance, an adjusted model added M2-like macrophage and M1-like inflammatory monocyte deconvolved proportions as fixed effects to isolate the hypoxia-specific effect:

$$MC\_Score \sim Hypoxia\_Score + Macrophages\_M2 + Inflammatory\_Monocytes\_M1 + (1 \mid Patient / Granuloma)$$

A full model added SPP1 expression to test independent contributions of hypoxia and SPP1 to MC score:

$$MC\_Score \sim Hypoxia\_Score + SPP1\_expression + Macrophages\_M2 + Inflammatory\_Monocytes\_M1 + (1 \mid Patient / Granuloma)$$

To visualize the joint contribution of hypoxia and SPP1 to MC score, each spot was classified into one of four categories based on binarized hypoxia and SPP1 status (Hyp-low/SPP1-low as reference, Hyp-high/SPP1-low, Hyp-low/SPP1-high, Hyp-high/SPP1-high), a four-category mixed-effects model was fitted with macrophage proportions as covariates. Marginal and conditional $R^2$ were computed using MuMIn[2]. Models were compared by AIC and likelihood ratio test.

#### Immunofluorescence Quantification

Serial sections were imaged with an Olympus microscope at 4x magnification as a tile scan. Raw, uncompressed images ( .oir format) were used for all quantification. Image quantifications were performed in QuPath (V0.6.0) and the data generated was imported to R for analysis.
In QuPath, for each granuloma, the necrotic center was manually annotated as a polygon following the boundary visible on the DAPI channel. Concentric distance bands (Zones) were then generated by expanding the core annotation outward at 25um increments to a maximum of 300um (Objects > Annotations> Expanded annotations, with remove interior and constraint to the parent object boxes checked). This produced 12 cumulative zones per granuloma. Three tissue-free background regions were annotated per granuloma.

Mean fluorescence intensity per channel was measured for the necrotic core, each zone and each background region using QuPath's pixel intensity features (pixel size was set to 3um). Background intensity was calculated as the mean of the three background regions per channel and subtracted from all core and zone measurements for that channel.

Because each zone included the preceding one, exclusive band intensities for non-overlapping 25um interval were calculated by area-weighted subtraction of consecutive zones.

Formula used:

Preceding zone Area: $A_1$; Preceding zone Mean Intensity: $I_1$

Following zone Area: $A_2$; Following zone Mean Intensity: $I_2$

The exclusive band intensity was then calculated as

$$[(A_2 * I_2) - (A1 * I_1)] / (A_2 - A_1)$$

The innermost zone and the necrotic core need no correction, as it was already exclusive by construction.

Given that cell density varied with distance from the necrotic core, band intensities for each marker were then normalized to the corresponding exclusive DAPI intensity with the same band as proxy for signal per cell. The DAPI normalized intensity for each marker was then plotted against distance.

### References:

1. Bates D, Mächler M, Bolker B, Walker S. Fitting Linear Mixed-Effects Models Using lme4. *J Stat Softw*. 2015;67:1-48. doi:10.18637/jss.v067.i01

2. Nakagawa S, Schielzeth H. A general and simple method for obtaining R2 from generalized linear mixed-effects models. *Methods Ecol Evol*. 2013;4(2). Accessed August 6, 2026. https://pub.uni-bielefeld.de/record/2565368

### Supplementary Notes

### Supplementary Note 1: Cluster Annotation

Using unsupervised clustering, we identified nine reproducible transcriptional microenvironments across the dataset. Cluster-defining marker genes (Fig. 3C, Supplementary Fig. 2) and deconvolved cell-type composition per ME (Fig. 3D, Supplementary Fig. 3B) were used to assign functional identity[1,2]. Because Visium spots capture multiple neighbouring cells, each ME represents a tissue programme rather than a single cell type.

Three MEs were macrophage-associated. The Foamy ME (Cluster 1) was defined by expression of *TREM2, LIPA, APOC2,* and *CD9*. Deconvolution showed M2-like macrophages as the dominant population (~0.36 of total signal) in this ME. These markers align with foamy macrophage states described in TB[3–5].The Baseline ME (Cluster 2) lacked strong programmatic enrichment. Deconvolution showed an even composition across macrophage subsets, T cells, and fibroblasts in this ME. The Glycolytic-Inflammatory ME (Cluster 3) co-expressed inflammatory genes (*IL1B, TREM1, CCL4, CD274*) and glycolytic genes (*SLC2A1, ALDOC, HK2*). This co-expression matches the signature of cells adapting to reduced oxygen availability[6,7]. This ME had elevated proportions of M2-like macrophages (~0.34), M1-skewed inflammatory monocytes (~0.20), and fibroblasts (~0.26).

Two MEs were stromal. The Fibrotic ME (Cluster 4) was characterized by expression of extracellular matrix and myofibroblast genes (*COL11A1, ACTA2, TAGLN*). Fibroblasts made up ~0.51 of the deconvolved composition. The Perivascular Stromal ME (Cluster 6) expressed *PDGFRA, COL15A1, TLR2,* and *IL1R1*, matching perivascular fibroblast populations in inflamed tissue. This ME comprised ~0.34 fibroblasts and ~0.23 M2-like macrophages, with other cell types at lower proportions. The Lymphoid-Vascular ME (Cluster 5) co-expressed B cell markers (CD19, MS4A1) and endothelial markers (VWF, CD34).

Three MEs were associated with necrotic regions. The Low-signal ME (Cluster 7) showed weak, non-specific gene expression, likely reflecting reduced transcriptional output in necrotic tissue. The Necrosis-associated Lymphoid ME (Cluster 8) co-expressed T cell receptor and immunoglobulin genes (TRAJ family genes, IGLV, IGHJ, MZB1). The Lipid-rich Necrotic ME

(Cluster 9) was defined by lipid metabolism genes (PLIN1, FABP4, CD36, LPL), CD163, and acute-phase response genes (SAA1, SAA2). This matches the lipid accumulation typical of caseous necrosis[4,8].

#### Supplementary Note 2: SPP1-hypoxia association across granulomas

Per-granuloma Spearman correlations between continuous hypoxia score and SPP1 expression showed a median positive association ($\rho = 0.25$, range -0.44 to 0.84) across the 60 granulomas. Thirty-one granulomas (51.7%) reached statistical significance after FDR correction. Median per-patient correlations ranged from $\rho = 0.07$ to 0.42.

A single granuloma (Sample_39, Granuloma_B) showed a statistically significant negative correlation ($\rho = -0.347$, FDR = 0.012). Inspection showed that this lesion lacked Cluster 3 representation, had no SPP1 expression, and had uniformly low hypoxia scores. This negative correlation therefore resulted from minimal biological variation rather than a true inverse association.

Variance decomposition from the linear mixed-effects model showed that patient identity accounted for 47.8% of the total variance in SPP1 expression ($\sigma^2 = 0.506$), while granuloma identity explained a further 13.5% ($\sigma^2 = 0.143$). Hypoxia score accounted for 9.6% of the variance as a fixed effect (marginal $R^2 = 0.096$), and the full model explained 65.0% of the total variance (conditional $R^2 = 0.650$).

#### Supplementary Note 3: MTB detection in Pediatric TB Granuloma Tissue.

Pediatric TB is widely reported to be paucibacillary, although the reasons for this remain debated. The reasons that have been raised so far include both the biology of the disease and the limited sensitivity of the available Mtb detection method in human tissue. To investigate the relationship between the Hypoxia-SPP1 niche identified in our spatial transcriptomics data and Mtb burden, we attempted to detect and quantify the Mtb in TB pediatric lymph node tissue from samples within the same pediatric cohort as the samples we used for spatial transcriptomics. Immunohistochemistry (IHC) was performed using Mtb-specific anti-PPD antibody and

RNAscope was done to target Mtb RNA, with both methods applied to enhance sensitivity detection. Across the cohort, most samples showed very low to rare MTB signals by both approaches. One sample showed a comparatively high Mtb burden. RNAscope consistently yielded lower signals than IHC across all samples tested (Supplementary Fig. 6). This discordance between the two methods made reliable quantification unfeasible. These findings confirm the paucibacillary nature of the pediatric tissues and motivated our use of a murine Mtb persistence-associated macrophage signature as a proxy for bacterial burden in downstream analysis.

### Supplementary Note 4: Persistence-associated macrophage signature analysis

MC scores ranged from -0.11 to 0.95 (median = 0.57). The unadjusted mixed-effects model estimated hypoxia score as a positive predictor of MC score ($\beta = 0.228$, SE = 0.006, marginal $R^2 = 0.043$, conditional $R^2 = 0.635$). The high conditional $R^2$ reflects the contribution of patient and granuloma identity to MC score variance.

The macrophage-adjusted model included M2-like macrophage and inflammatory monocyte (M1-like) proportions as additional fixed effects. Under this model, hypoxia score remained a positive predictor ($\beta = 0.114$, SE = 0.005). Inflammatory monocyte proportion contributed positively ($\beta = 0.195$, $p < 2 \times 10^{-16}$). Marginal $R^2$ increased from 0.043 to 0.363. The adjusted model outperformed the basic model by AIC ($\Delta$AIC $\approx$ 5,959) and likelihood ratio testing ($\chi^2 = 5{,}962.8$, $p < 2.2 \times 10^{-16}$).

In the full model with hypoxia, SPP1, and macrophage proportions as fixed effects, the hypoxia coefficient was further attenuated ($\beta = 0.114 \rightarrow 0.087$), indicating that part of the macrophage-adjusted hypoxia effect was shared with SPP1 expression.

For the joint visualization, spots were classified into four categories:

Hyp-low/SPP1-low (n = 9,495; reference),

Hyp-high/SPP1-low (n = 2,164)

Hyp-low/SPP1-high (n = 1,334)

Hyp-high/SPP1-high (n = 1,227)

The joint category coefficient (β = 0.045) approximated the sum of the two single-positive coefficients (Hyp-high/SPP1-low β = 0.019; Hyp-low/SPP1-high β = 0.023), consistent with additive rather than synergistic contribution to MC score.

### References:


1. Cable DM, Murray E, Zou LS, et al. Robust decomposition of cell type mixtures in spatial transcriptomics. *Nat Biotechnol*. 2022;40(4):517-526. doi:10.1038/s41587-021-00830-w

2. Ståhl PL, Salmén F, Vickovic S, et al. Visualization and analysis of gene expression in tissue sections by spatial transcriptomics. *Science*. 2016;353(6294):78-82. doi:10.1126/science.aaf2403

3. Peyron P, Vaubourgeix J, Poquet Y, et al. Foamy Macrophages from Tuberculous Patients' Granulomas Constitute a Nutrient-Rich Reservoir for M. tuberculosis Persistence. *PLoS Pathog*. 2008;4(11):e1000204. doi:10.1371/journal.ppat.1000204

4. Russell DG, Cardona PJ, Kim MJ, Allain S, Altare F. Foamy macrophages and the progression of the human TB granuloma. *Nat Immunol*. 2009;10(9):943-948. doi:10.1038/ni.1781

5. Pisu D, Huang L, Narang V, et al. Single cell analysis of M. tuberculosis phenotype and macrophage lineages in the infected lung. *J Exp Med*. 2021;218(9):e20210615. doi:10.1084/jem.20210615

6. Shi L, Salamon H, Eugenin EA, Pine R, Cooper A, Gennaro ML. Infection with Mycobacterium tuberculosis induces the Warburg effect in mouse lungs. *Sci Rep*. 2015;5(1):18176. doi:10.1038/srep18176

7. Zhong WJ, Liu T, Yang HH, et al. TREM-1 governs NLRP3 inflammasome activation of macrophages by firing up glycolysis in acute lung injury. *Int J Biol Sci*. 2023;19(1):242-257. doi:10.7150/ijbs.77304

8. Kim MJ, Wainwright HC, Locketz M, et al. Caseation of human tuberculosis granulomas correlates with elevated host lipid metabolism. *EMBO Mol Med*. 2010;2(7):258-274. doi:10.1002/emmm.201000079

9. Bates D, Mächler M, Bolker B, Walker S. Fitting Linear Mixed-Effects Models Using lme4. *J Stat Softw*. 2015;67:1-48. doi:10.18637/jss.v067.i01

10. Nakagawa S, Schielzeth H. A general and simple method for obtaining R2 from generalized linear mixed-effects models. *Methods Ecol Evol*. 2013;4(2). Accessed August 6, 2026. https://pub.uni-bielefeld.de/record/2565368

### Supplementary Tables

**Supplementary Table 1.** Clinical and sample characteristics of TB lymph node donors included in spatial transcriptomics analysis.

| Patient ID | Sex | Age (months) | HIV status | TB treatment status | Sections (n) | Granulomas annotated (n) | Spots after QC (n) | Necrotic granulomas (n) | Non-necrotic granulomas (n) | ZN status |
|---|---|---|---|---|---|---|---|---|---|---|
| Patient_1 | Female | 4 | Negative | Treated | 1 | 3 | 2,623 | 0 | 3 | Positive |
| Patient_2 | Female | 18 | Negative | Treated | 2 | 12 | 4,178 | 5 | 7 | Negative |
| Patient_3 | Male | 36 | Negative | Treated | 2 | 34 | 3,428 | 0 | 34 | Negative |
| Patient_4 | Male | 6 | Negative | Treated | 2 | 11 | 3,997 | 9 | 2 | Positive |

QC = quality control; ZN = Ziehl-Neelsen acid-fast staining; n = number.

All patients were HIV-negative and receiving anti-tuberculosis treatment at the time of tissue biopsy. One tissue section from Patient_1 was excluded from analysis due to processing failure. ZN status refers to Ziehl-Neelsen acid-fast bacilli staining of the original biopsy specimen.

Supplementary Table 2

| Cluster | Pathway | NES | P_value | Padj | Gene_set_size | Direction |
|---|---|---|---|---|---|---|
| 1 | Complement | 1.5979 | 0.0001462 | 0.003776 | 62 | Activated |
| 1 | Uv Response Up | 1.6387 | 0.0002575 | 0.003776 | 51 | Activated |
| 1 | Xenobiotic Metabolism | 1.5787 | 0.0001825 | 0.003776 | 70 | Activated |
| 1 | Estrogen Response Late | 1.59 | 0.001095 | 0.01205 | 48 | Activated |
| 1 | Coagulation | 1.7035 | 0.001419 | 0.01248 | 22 | Activated |
| 1 | Adipogenesis | 1.4329 | 0.0032 | 0.01564 | 88 | Activated |
| 1 | Fatty Acid Metabolism | 1.4855 | 0.003 | 0.01564 | 68 | Activated |
| 1 | Kras Signaling Up | 1.5967 | 0.002498 | 0.01564 | 42 | Activated |
| 1 | Tnfa Signaling Via Nfkb | 1.5202 | 0.0023 | 0.01564 | 59 | Activated |
| 1 | Bile Acid Metabolism | 1.5253 | 0.006102 | 0.02685 | 36 | Activated |
| 1 | Androgen Response | 1.5361 | 0.00741 | 0.02841 | 32 | Activated |
| 1 | Reactive Oxygen Species Pathway | 1.5824 | 0.007748 | 0.02841 | 24 | Activated |
| 1 | Mtorc1 Signaling | 1.3744 | 0.009799 | 0.03317 | 82 | Activated |
| 2 | Epithelial Mesenchymal Transition | -1.5712 | 0.01364 | 0.01364 | 15 | Suppressed |
| 3 | Complement | 1.4289 | 0.0005358 | 0.01522 | 56 | Activated |
| 3 | Hypoxia | 1.4094 | 0.001114 | 0.01522 | 58 | Activated |
| 3 | Tnfa Signaling Via Nfkb | 1.3565 | 0.0009296 | 0.01522 | 86 | Activated |
| 3 | Kras Signaling Up | 1.4038 | 0.0025 | 0.02562 | 49 | Activated |
| 3 | Il2 Stat5 Signaling | 1.3449 | 0.0042 | 0.03444 | 62 | Activated |
| 3 | Glycolysis | 1.3698 | 0.005099 | 0.03485 | 52 | Activated |
| 3 | Epithelial Mesenchymal Transition | 1.4506 | 0.006608 | 0.0387 | 28 | Activated |
| 3 | Allograft Rejection | 1.3568 | 0.009999 | 0.04223 | 48 | Activated |
| 3 | Inflammatory Response | 1.2949 | 0.0103 | 0.04223 | 76 | Activated |
| 3 | Mtorc1 Signaling | 1.3079 | 0.009599 | 0.04223 | 69 | Activated |
| 4 | Epithelial Mesenchymal Transition | 1.3834 | 1.309e-06 | 1.047e-05 | 78 | Activated |
| 5 | Kras Signaling Up | 1.7159 | 2.494e-06 | 0.0001097 | 70 | Activated |
| 5 | Inflammatory Response | 1.7253 | 1.174e-05 | 0.0002583 | 60 | Activated |
| 5 | Myogenesis | 1.6912 | 7.389e-05 | 0.001084 | 51 | Activated |
| 5 | Apical Junction | 1.5739 | 0.0002143 | 0.001885 | 76 | Activated |
| 5 | Interferon Gamma Response | 1.59 | 0.0001757 | 0.001885 | 79 | Activated |
| 5 | Adipogenesis | 1.6228 | 0.0003076 | 0.002256 | 69 | Activated |
| 5 | Interferon Alpha Response | 1.6414 | 0.0003987 | 0.002506 | 49 | Activated |
| 5 | Coagulation | 1.7011 | 0.0004817 | 0.002649 | 34 | Activated |
| 5 | Tgf Beta Signaling | 1.669 | 0.002108 | 0.0103 | 28 | Activated |
| 5 | Kras Signaling Dn | 1.6418 | 0.002406 | 0.01058 | 31 | Activated |
| 5 | Estrogen Response Early | 1.4564 | 0.0049 | 0.01737 | 70 | Activated |

| | | | | | | |
|---|---|---|---|---|---|---|
| **5** | Il2 Stat5 Signaling | 1.4309 | 0.005099 | 0.01737 | 82 | Activated |
| **5** | Il6 Jak Stat3 Signaling | 1.6316 | 0.005133 | 0.01737 | 24 | Activated |
| **5** | Uv Response Dn | 1.3982 | 0.008399 | 0.0264 | 84 | Activated |
| **5** | Complement | 1.4364 | 0.0121 | 0.03549 | 55 | Activated |
| **5** | Epithelial Mesenchymal Transition | 1.3985 | 0.0156 | 0.0429 | 66 | Activated |
| **6** | Epithelial Mesenchymal Transition | 1.9049 | 2.15e-06 | 9.458e-05 | 69 | Activated |
| **6** | Myogenesis | 1.8702 | 8.878e-05 | 0.001953 | 40 | Activated |
| **6** | Coagulation | 1.7203 | 0.002718 | 0.02816 | 32 | Activated |
| **6** | Inflammatory Response | 1.5439 | 0.0032 | 0.02816 | 71 | Activated |
| **6** | Kras Signaling Up | 1.6084 | 0.0024 | 0.02816 | 59 | Activated |
| **7** | Epithelial Mesenchymal Transition | -2.6773 | 1.025e-08 | 3.586e-07 | 47 | Suppressed |
| **7** | Coagulation | -2.3076 | 0.0001075 | 0.001881 | 18 | Suppressed |

Supplementary Table 3

| Cluster | Total_spots | % of hypoxia-high spots | % of hypoxia-low spots | Median SPP1 expression | SPP1-high spots in cluster (%) | Both hypoxia-high and SPP1-high (%) | Odds ratio | FDR p-value | Sig | Direction |
|---|---|---|---|---|---|---|---|---|---|---|
| 1 | 2977 | 19.6 | 21.3 | 1.099 | 13.2 | 5.1 | 0.9 | 0.0429 | * | depleted |
| 2 | 2266 | 14.1 | 16.5 | 1.386 | 17.3 | 4.6 | 0.827 | 0.00104 | ** | depleted |
| 3 | 1991 | 33.1 | 8 | 2.773 | 55 | 40.3 | 5.705 | 1.12e-255 | *** | enriched |
| 4 | 1970 | 7.6 | 15.8 | 0.693 | 4.7 | 1 | 0.44 | 2.28e-36 | *** | depleted |
| 5 | 1735 | 5.5 | 14.3 | 0.693 | 7.6 | 1.7 | 0.345 | 5.45e-49 | *** | depleted |
| 6 | 1528 | 7.5 | 11.7 | 1.099 | 7.6 | 1.6 | 0.613 | 2.7e-12 | *** | depleted |
| 7 | 1227 | 7.9 | 8.9 | 1.099 | 15.3 | 3.8 | 0.883 | 0.0968 | | depleted |
| 8 | 322 | 2.9 | 2.1 | 1.099 | 39.1 | 13 | 1.408 | 0.00988 | ** | enriched |
| 9 | 210 | 1.8 | 1.4 | 1.792 | 12.9 | 3.3 | 1.282 | 0.121 | | enriched |

Supplementary Table 4

| gene | avg_log2FC | pct.1 | pct.2 | p_val | p_val_adj | direction | gene_category |
|---|---|---|---|---|---|---|---|
| **HK2** | 1.3193 | 0.714 | 0.433 | 0 | 0 | Up in Hypoxia-high | Canonical hypoxia |
| **EGLN3** | 1.5155 | 0.493 | 0.265 | 5.862e-309 | 1.046e-304 | Up in Hypoxia-high | Canonical hypoxia |
| **SPP1** | 1.5518 | 0.864 | 0.791 | 1.633e-306 | 2.915e-302 | Up in Hypoxia-high | SPP1 |
| **SLC16A3** | 1.0224 | 0.752 | 0.509 | 8.45300000000000e-304 | 1.508e-299 | Up in Hypoxia-high | Canonical hypoxia |
| **ENO1** | 0.4496 | 1 | 0.997 | 4.256e-273 | 7.594e-269 | Up in Hypoxia-high | Canonical hypoxia |
| **VEGFA** | 1.1074 | 0.699 | 0.413 | 6.106e-265 | 1.09e-260 | Up in Hypoxia-high | Canonical hypoxia |
| **SLC2A1** | 1.6072 | 0.481 | 0.213 | 1.642e-252 | 2.931e-248 | Up in Hypoxia-high | Canonical hypoxia |
| **PGAM1** | 0.6485 | 0.897 | 0.726 | 2.303e-220 | 4.109e-216 | Up in Hypoxia-high | Canonical hypoxia |
| **PFKFB3** | 0.6293 | 0.794 | 0.653 | 9.189e-152 | 1.64e-147 | Up in Hypoxia-high | Canonical hypoxia |
| **CP** | 0.675 | 0.842 | 0.727 | 1.709e-138 | 3.049e-134 | Up in Hypoxia-high | Other |
| **MT2A** | 0.3927 | 1 | 0.998 | 2.321e-135 | 4.141e-131 | Up in Hypoxia-high | Other |
| **NCF1** | 0.7157 | 0.788 | 0.683 | 4.261e-132 | 7.603e-128 | Up in Hypoxia-high | Other |
| **G0S2** | 0.5027 | 0.968 | 0.93 | 1.447e-125 | 2.581e-121 | Up in Hypoxia-high | Other |
| **FCGR2B** | 1.0241 | 0.494 | 0.351 | 4.857e-125 | 8.666e-121 | Up in Hypoxia-high | Macrophage/immune |
| **FBP1** | 0.4879 | 0.984 | 0.958 | 1.503e-124 | 2.681e-120 | Up in Hypoxia-high | Other |
| **PLAUR** | 0.3898 | 0.995 | 0.98 | 1.62e-123 | 2.891e-119 | Up in Hypoxia-high | Other |
| **NDRG1** | 0.6425 | 0.793 | 0.718 | 2.546e-122 | 4.543e-118 | Up in Hypoxia-high | Other |
| **LYZ** | 0.3818 | 1 | 1 | 8.443e-119 | 1.507e-114 | Up in Hypoxia-high | Macrophage/immune |
| **SLC11A1** | 0.5295 | 0.86 | 0.745 | 1.817e-118 | 3.242e-114 | Up in Hypoxia-high | Macrophage/immune |
| **CD14** | 0.383 | 0.994 | 0.986 | 8.31e-118 | 1.483e-113 | Up in Hypoxia-high | Macrophage/immune |
| **CAMP** | 1.3577 | 0.268 | 0.171 | 2.553e-113 | 4.555e-109 | Up in Hypoxia-high | Macrophage/immune |
| **SRGN** | 0.4025 | 0.995 | 0.988 | 2.258e-112 | 4.03e-108 | Up in Hypoxia-high | Other |
| **CSTB** | 0.4158 | 0.986 | 0.977 | 7.175e-111 | 1.28e-106 | Up in Hypoxia-high | Other |
| **IL4I1** | 0.4309 | 0.892 | 0.804 | 3.865e-103 | 6.896e-99 | Up in Hypoxia-high | Other |
| **MT1G** | 0.6535 | 0.832 | 0.737 | 4.078e-101 | 7.276e-97 | Up in Hypoxia-high | Other |
| **ITGB2** | 0.342 | 0.999 | 0.996 | 6.998e-100 | 1.249e-95 | Up in Hypoxia-high | Other |
| **VIM** | 0.2949 | 1 | 1 | 2.36e-99 | 4.211e-95 | Up in Hypoxia-high | Other |
| **SOD2** | 0.2872 | 1 | 1 | 6.854e-99 | 1.223e-94 | Up in Hypoxia-high | Macrophage/immune |
| **ATF5** | 0.4464 | 0.883 | 0.829 | 7.774e-98 | 1.387e-93 | Up in Hypoxia-high | Other |
| **TREM1** | 0.8456 | 0.507 | 0.362 | 2.483e-97 | 4.43e-93 | Up in Hypoxia-high | Macrophage/immune |
| **HIF1A** | 0.4555 | 0.834 | 0.746 | 2.065e-96 | 3.684e-92 | Up in Hypoxia-high | Canonical hypoxia |
| **S100A10** | 0.3468 | 0.995 | 0.987 | 1.442e-94 | 2.572e-90 | Up in Hypoxia-high | Other |
| **ERO1A** | 0.55 | 0.775 | 0.695 | 2.024e-94 | 3.611e-90 | Up in Hypoxia-high | Other |
| **FTH1** | 0.2162 | 1 | 1 | 1.726e-91 | 3.08e-87 | Up in Hypoxia-high | Other |
| **NUPR1** | 0.2986 | 0.999 | 0.993 | 4.53e-85 | 8.084e-81 | Up in Hypoxia-high | Macrophage/immune |
| **JCHAIN** | -0.4917 | 0.899 | 0.937 | 9.654e-85 | 1.723e-80 | Up in Hypoxia-low | Other |
| **IGFBP7** | -0.3395 | 0.976 | 0.988 | 3.123e-84 | 5.573e-80 | Up in Hypoxia-low | Other |
| **MMP9** | 0.3889 | 0.984 | 0.97 | 1.758e-82 | 3.138e-78 | Up in Hypoxia-high | Macrophage/immune |
| **C15orf48** | 0.3594 | 0.982 | 0.956 | 3.457e-81 | 6.169e-77 | Up in Hypoxia-high | Other |
| **HCST** | 0.5621 | 0.736 | 0.69 | 7.565e-81 | 1.35e-76 | Up in Hypoxia-high | Other |
| **WARS** | 0.2313 | 1 | 1 | 2.017e-80 | 3.599e-76 | Up in Hypoxia-high | Other |
| **MT1E** | 0.5085 | 0.859 | 0.781 | 2.456e-78 | 4.383e-74 | Up in Hypoxia-high | Other |

| | | | | | | | |
|---|---|---|---|---|---|---|---|
| **B2M** | 0.1347 | 1 | 1 | 7.232e-75 | 1.29e-70 | Up in Hypoxia-high | Other |
| **FCER1G** | 0.2526 | 0.996 | 0.992 | 2.356e-74 | 4.204e-70 | Up in Hypoxia-high | Other |
| **CD44** | 0.2788 | 0.999 | 0.996 | 4.913e-74 | 8.766e-70 | Up in Hypoxia-high | Other |
| **CA12** | 0.5155 | 0.698 | 0.611 | 1.077e-72 | 1.922e-68 | Up in Hypoxia-high | Other |
| **CTSB** | 0.1977 | 1 | 1 | 4.433e-72 | 7.91e-68 | Up in Hypoxia-high | Other |
| **MT1H** | 0.6593 | 0.547 | 0.46 | 9.176e-71 | 1.637e-66 | Up in Hypoxia-high | Other |
| **MARCO** | 0.4911 | 0.724 | 0.657 | 1.298e-70 | 2.315e-66 | Up in Hypoxia-high | Other |
| **ITGAM** | 0.6751 | 0.476 | 0.381 | 3.433e-69 | 6.125e-65 | Up in Hypoxia-high | Other |
| **LOX** | 0.9672 | 0.352 | 0.217 | 7.475e-69 | 1.334e-64 | Up in Hypoxia-high | Canonical hypoxia |
| **POSTN** | -0.3778 | 0.864 | 0.933 | 2.101e-68 | 3.749e-64 | Up in Hypoxia-low | Other |
| **ZNF385A** | 0.4826 | 0.731 | 0.661 | 2.164e-68 | 3.861e-64 | Up in Hypoxia-high | Other |
| **IGKC** | -0.4293 | 0.998 | 0.999 | 3.087e-66 | 5.508e-62 | Up in Hypoxia-low | Other |
| **CCL5** | 0.4173 | 0.96 | 0.948 | 8.825e-65 | 1.575e-60 | Up in Hypoxia-high | Other |
| **MIF** | 0.3777 | 0.944 | 0.926 | 1.199e-64 | 2.139e-60 | Up in Hypoxia-high | Other |
| **CD68** | 0.2446 | 0.999 | 0.997 | 3.953e-63 | 7.054e-59 | Up in Hypoxia-high | Macrophage/immune |
| **HLA-DQA1** | 0.2754 | 0.997 | 0.996 | 8.978e-62 | 1.602e-57 | Up in Hypoxia-high | Other |
| **IDO1** | 0.2916 | 0.948 | 0.92 | 2.106e-61 | 3.759e-57 | Up in Hypoxia-high | Other |
| **ACSL1** | 0.287 | 0.983 | 0.947 | 7.078e-60 | 1.263e-55 | Up in Hypoxia-high | Other |
| **PLTP** | 0.3845 | 0.877 | 0.828 | 9.856e-60 | 1.759e-55 | Up in Hypoxia-high | Other |
| **CXCL12** | -0.3544 | 0.782 | 0.864 | 3.363e-59 | 6E-55 | Up in Hypoxia-low | Other |
| **RNASET2** | 0.278 | 0.986 | 0.977 | 1.009e-58 | 1.801e-54 | Up in Hypoxia-high | Other |
| **SPARC** | -0.2358 | 0.996 | 0.997 | 2.318e-57 | 4.137e-53 | Up in Hypoxia-low | Other |
| **SQOR** | 0.2536 | 0.993 | 0.983 | 7.793e-56 | 1.391e-51 | Up in Hypoxia-high | Other |
| **CCL19** | -0.3149 | 0.983 | 0.991 | 1.185e-55 | 2.115e-51 | Up in Hypoxia-low | Other |
| **GLUL** | 0.1675 | 0.999 | 0.999 | 2.453e-55 | 4.377e-51 | Up in Hypoxia-high | Other |
| **TYROBP** | 0.1967 | 1 | 1 | 1.845e-54 | 3.292e-50 | Up in Hypoxia-high | Other |
| **IGHG1** | -0.4656 | 0.986 | 0.992 | 1.91e-54 | 3.408e-50 | Up in Hypoxia-low | Other |
| **GBP5** | 0.1773 | 0.995 | 0.988 | 4.288e-54 | 7.652e-50 | Up in Hypoxia-high | Other |
| **GPI** | 0.3247 | 0.958 | 0.939 | 1.955e-53 | 3.488e-49 | Up in Hypoxia-high | Other |
| **S100A6** | 0.2312 | 1 | 1 | 7.784e-52 | 1.389e-47 | Up in Hypoxia-high | Other |
| **COL3A1** | -0.1833 | 1 | 1 | 7.962e-52 | 1.421e-47 | Up in Hypoxia-low | Other |
| **CA2** | 1.0445 | 0.225 | 0.122 | 2.01e-51 | 3.587e-47 | Up in Hypoxia-high | Other |
| **TPM1** | -0.3507 | 0.751 | 0.834 | 5.607e-51 | 1E-46 | Up in Hypoxia-low | Other |
| **STOM** | 0.3048 | 0.961 | 0.933 | 1.09e-50 | 1.945e-46 | Up in Hypoxia-high | Other |
| **ACTA2** | -0.3607 | 0.821 | 0.873 | 1.582e-50 | 2.822e-46 | Up in Hypoxia-low | Other |
| **CD74** | 0.1599 | 1 | 1 | 3.277e-50 | 5.847e-46 | Up in Hypoxia-high | Other |
| **HK3** | 0.4222 | 0.743 | 0.656 | 1.012e-49 | 1.806e-45 | Up in Hypoxia-high | Other |
| **IGHM** | -0.4205 | 0.856 | 0.907 | 4.981e-49 | 8.888e-45 | Up in Hypoxia-low | Other |
| **ICAM1** | 0.2662 | 0.984 | 0.969 | 4.717e-48 | 8.416e-44 | Up in Hypoxia-high | Other |
| **COL1A2** | -0.2335 | 0.999 | 1 | 1.2e-47 | 2.142e-43 | Up in Hypoxia-low | Other |
| **XBP1** | -0.2753 | 0.956 | 0.974 | 1.556e-47 | 2.776e-43 | Up in Hypoxia-low | Other |
| **SERPINF1** | -0.2452 | 0.955 | 0.973 | 3.506e-47 | 6.256e-43 | Up in Hypoxia-low | Other |
| **AQP9** | 0.3911 | 0.699 | 0.595 | 4.251e-47 | 7.584e-43 | Up in Hypoxia-high | Other |
| **MZB1** | -0.5307 | 0.563 | 0.661 | 4.43e-47 | 7.905e-43 | Up in Hypoxia-low | Other |

| | | | | | | | |
|---|---|---|---|---|---|---|---|
| **MT1X** | 0.4653 | 0.627 | 0.55 | 5.05e-47 | 9.01e-43 | Up in Hypoxia-high | Other |
| **MYL9** | -0.3947 | 0.774 | 0.842 | 1.041e-46 | 1.857e-42 | Up in Hypoxia-low | Other |
| **CTSS** | 0.1797 | 0.999 | 0.998 | 2.406e-46 | 4.293e-42 | Up in Hypoxia-high | Other |
| **TAGLN** | -0.3919 | 0.746 | 0.814 | 2.477e-46 | 4.421e-42 | Up in Hypoxia-low | Other |
| **DCN** | -0.1906 | 0.983 | 0.988 | 3.075e-46 | 5.486e-42 | Up in Hypoxia-low | Other |
| **SAT1** | 0.1458 | 0.999 | 0.998 | 5.359e-46 | 9.562e-42 | Up in Hypoxia-high | Other |
| **IL1B** | 0.7747 | 0.442 | 0.347 | 6.432e-46 | 1.148e-41 | Up in Hypoxia-high | Macrophage/immune |
| **NINJ1** | 0.298 | 0.894 | 0.841 | 9.207e-46 | 1.643e-41 | Up in Hypoxia-high | Other |
| **VCAN** | -0.3766 | 0.712 | 0.793 | 2.259e-45 | 4.032e-41 | Up in Hypoxia-low | Other |
| **IGHA1** | -0.4629 | 0.972 | 0.986 | 2.267e-45 | 4.045e-41 | Up in Hypoxia-low | Other |
| **CLEC6A** | 0.6088 | 0.377 | 0.263 | 2.697e-45 | 4.813e-41 | Up in Hypoxia-high | Other |
| **MMP2** | -0.352 | 0.869 | 0.899 | 1.369e-44 | 2.442e-40 | Up in Hypoxia-low | Other |
| **HCAR2** | 0.9965 | 0.208 | 0.116 | 1.99e-44 | 3.551e-40 | Up in Hypoxia-high | Canonical hypoxia |
| **IGLC1** | -0.5067 | 0.892 | 0.925 | 5.857e-44 | 1.045e-39 | Up in Hypoxia-low | Other |
| **IFITM3** | -0.2335 | 0.996 | 0.998 | 5.099e-43 | 9.098e-39 | Up in Hypoxia-low | Other |
| **ENO2** | 0.87 | 0.247 | 0.148 | 1.279e-42 | 2.281e-38 | Up in Hypoxia-high | Other |
| **IGHJ6** | -0.7551 | 0.729 | 0.774 | 4.999e-42 | 8.919e-38 | Up in Hypoxia-low | Other |
| **CHI3L1** | 0.2124 | 0.991 | 0.975 | 1.066e-41 | 1.901e-37 | Up in Hypoxia-high | Other |
| **C1QA** | 0.2389 | 0.998 | 0.996 | 2.337e-41 | 4.17e-37 | Up in Hypoxia-high | Other |
| **CCR1** | 0.2964 | 0.813 | 0.734 | 7.49e-41 | 1.336e-36 | Up in Hypoxia-high | Other |
| **FTL** | 0.0836 | 1 | 1 | 8.267e-41 | 1.475e-36 | Up in Hypoxia-high | Other |
| **FBLN1** | -0.3328 | 0.743 | 0.792 | 1.686e-40 | 3.008e-36 | Up in Hypoxia-low | Other |
| **HLA-DRA** | 0.1428 | 1 | 1 | 3.157e-40 | 5.632e-36 | Up in Hypoxia-high | Other |
| **S100A11** | 0.1626 | 0.999 | 0.996 | 3.621e-40 | 6.46e-36 | Up in Hypoxia-high | Other |
| **ALOX5** | 0.3057 | 0.891 | 0.831 | 9.833e-40 | 1.754e-35 | Up in Hypoxia-high | Other |
| **AQP1** | -0.6001 | 0.331 | 0.428 | 1.116e-39 | 1.992e-35 | Up in Hypoxia-low | Other |
| **SNX10** | 0.2358 | 0.922 | 0.864 | 1.177e-39 | 2.101e-35 | Up in Hypoxia-high | Other |
| **COL1A1** | -0.2255 | 0.999 | 1 | 6.49e-39 | 1.158e-34 | Up in Hypoxia-low | Other |
| **TXNDC5** | -0.3997 | 0.636 | 0.728 | 6.776e-39 | 1.209e-34 | Up in Hypoxia-low | Other |
| **BCAT1** | 0.2549 | 0.941 | 0.899 | 1.493e-38 | 2.663e-34 | Up in Hypoxia-high | Other |
| **PGD** | 0.1948 | 0.995 | 0.989 | 1.771e-38 | 3.16e-34 | Up in Hypoxia-high | Other |
| **LAPTM5** | 0.1253 | 1 | 1 | 1.952e-38 | 3.483e-34 | Up in Hypoxia-high | Other |
| **PLD3** | 0.19 | 0.997 | 0.995 | 4.837e-38 | 8.631e-34 | Up in Hypoxia-high | Other |
| **FBN1** | -0.2508 | 0.876 | 0.89 | 5.954e-38 | 1.062e-33 | Up in Hypoxia-low | Other |
| **SERPINA1** | 0.3771 | 0.716 | 0.639 | 8E-38 | 1.427e-33 | Up in Hypoxia-high | Other |
| **IL18BP** | 0.4031 | 0.666 | 0.589 | 8.076e-38 | 1.441e-33 | Up in Hypoxia-high | Other |
| **SFRP2** | -0.5914 | 0.355 | 0.417 | 1.236e-37 | 2.205e-33 | Up in Hypoxia-low | Other |
| **CDA** | 0.6273 | 0.368 | 0.261 | 6.578e-37 | 1.174e-32 | Up in Hypoxia-high | Other |
| **IGHG2** | -0.5779 | 0.913 | 0.934 | 6.797e-37 | 1.213e-32 | Up in Hypoxia-low | Other |
| **SULF1** | -0.3818 | 0.66 | 0.738 | 1.069e-36 | 1.908e-32 | Up in Hypoxia-low | Other |
| **MRC1** | 0.6932 | 0.296 | 0.213 | 2.99e-36 | 5.335e-32 | Up in Hypoxia-high | Other |
| **S100A4** | 0.2662 | 0.957 | 0.93 | 3.625e-36 | 6.468e-32 | Up in Hypoxia-high | Other |
| **IFI6** | 0.1197 | 0.978 | 0.96 | 4.765e-36 | 8.502e-32 | Up in Hypoxia-high | Other |
| **IGKV4-1** | -0.4893 | 0.861 | 0.897 | 4.821e-36 | 8.602e-32 | Up in Hypoxia-low | Other |

| | | | | | | | |
|---|---|---|---|---|---|---|---|
| **CAPG** | 0.165 | 0.999 | 0.995 | 8.01e-36 | 1.429e-31 | Up in Hypoxia-high | Other |
| **CCN2** | -0.3361 | 0.791 | 0.843 | 9.103e-36 | 1.624e-31 | Up in Hypoxia-low | Other |
| **RARRES2** | -0.2917 | 0.852 | 0.88 | 1.192e-35 | 2.128e-31 | Up in Hypoxia-low | Other |
| **C3** | -0.1894 | 0.971 | 0.976 | 1.795e-35 | 3.203e-31 | Up in Hypoxia-low | Other |
| **SLC39A8** | 0.3443 | 0.836 | 0.799 | 6.793e-35 | 1.212e-30 | Up in Hypoxia-high | Other |
| **MS4A6A** | -0.3857 | 0.523 | 0.629 | 1.408e-34 | 2.512e-30 | Up in Hypoxia-low | Other |
| **CCL21** | -0.6231 | 0.733 | 0.768 | 1.664e-34 | 2.969e-30 | Up in Hypoxia-low | Other |
| **P4HA1** | 0.5602 | 0.381 | 0.289 | 2.502e-34 | 4.465e-30 | Up in Hypoxia-high | Other |
| **QPCT** | 0.4143 | 0.609 | 0.529 | 3.172e-34 | 5.661e-30 | Up in Hypoxia-high | Other |
| **VSIG4** | 0.1978 | 0.961 | 0.919 | 3.54e-34 | 6.316e-30 | Up in Hypoxia-high | Other |
| **THY1** | -0.3735 | 0.583 | 0.673 | 1.724e-33 | 3.077e-29 | Up in Hypoxia-low | Other |
| **CALD1** | -0.3762 | 0.555 | 0.649 | 1.878e-33 | 3.35e-29 | Up in Hypoxia-low | Other |
| **TPM2** | -0.2853 | 0.859 | 0.884 | 2.068e-33 | 3.69e-29 | Up in Hypoxia-low | Other |
| **NCF2** | 0.2844 | 0.88 | 0.83 | 2.189e-33 | 3.906e-29 | Up in Hypoxia-high | Other |
| **STX11** | 0.2559 | 0.911 | 0.859 | 2.698e-33 | 4.814e-29 | Up in Hypoxia-high | Other |
| **BGN** | -0.2246 | 0.935 | 0.947 | 6.52e-33 | 1.163e-28 | Up in Hypoxia-low | Other |
| **TNFAIP6** | 0.607 | 0.375 | 0.287 | 1.014e-32 | 1.81e-28 | Up in Hypoxia-high | Other |
| **APLP2** | 0.2305 | 0.95 | 0.927 | 1.998e-32 | 3.565e-28 | Up in Hypoxia-high | Other |
| **TXN** | 0.1704 | 0.996 | 0.993 | 3.053e-32 | 5.448e-28 | Up in Hypoxia-high | Other |
| **POU2AF1** | -0.6267 | 0.276 | 0.377 | 3.418e-32 | 6.098e-28 | Up in Hypoxia-low | Other |
| **MT1F** | 0.5292 | 0.37 | 0.279 | 9.885e-32 | 1.764e-27 | Up in Hypoxia-high | Other |
| **S100A8** | 0.1906 | 0.976 | 0.948 | 1.952e-31 | 3.482e-27 | Up in Hypoxia-high | Other |
| **C7** | -0.8678 | 0.266 | 0.338 | 2.455e-31 | 4.38e-27 | Up in Hypoxia-low | Other |
| **AEBP1** | -0.2755 | 0.937 | 0.949 | 9.934e-31 | 1.772e-26 | Up in Hypoxia-low | Other |
| **CCL8** | 0.5641 | 0.273 | 0.204 | 1.147e-30 | 2.047e-26 | Up in Hypoxia-high | Other |
| **PILRA** | 0.281 | 0.801 | 0.722 | 1.168e-30 | 2.084e-26 | Up in Hypoxia-high | Other |
| **BRI3** | 0.1057 | 0.999 | 0.998 | 1.427e-30 | 2.546e-26 | Up in Hypoxia-high | Other |
| **CD82** | 0.2347 | 0.927 | 0.896 | 1.499e-30 | 2.674e-26 | Up in Hypoxia-high | Other |
| **ID1** | -0.4487 | 0.444 | 0.519 | 2.341e-30 | 4.177e-26 | Up in Hypoxia-low | Other |
| **PLA2G2D** | -0.4588 | 0.509 | 0.594 | 3.014e-30 | 5.379e-26 | Up in Hypoxia-low | Other |
| **EGR2** | 0.4828 | 0.418 | 0.374 | 5.47e-30 | 9.76e-26 | Up in Hypoxia-high | Other |
| **OSCAR** | 0.3748 | 0.618 | 0.528 | 6.448e-30 | 1.151e-25 | Up in Hypoxia-high | Other |
| **IGF1** | -0.737 | 0.166 | 0.238 | 9.827e-30 | 1.753e-25 | Up in Hypoxia-low | Other |
| **UBC** | 0.0994 | 1 | 1 | 9.881e-30 | 1.763e-25 | Up in Hypoxia-high | Other |
| **CTSK** | -0.2444 | 0.879 | 0.901 | 2.104e-29 | 3.754e-25 | Up in Hypoxia-low | Other |
| **ITM2C** | -0.4396 | 0.401 | 0.492 | 2.263e-29 | 4.038e-25 | Up in Hypoxia-low | Other |
| **HLA-C** | 0.1741 | 0.916 | 0.885 | 2.711e-29 | 4.837e-25 | Up in Hypoxia-high | Other |
| **SLC2A3** | 0.5267 | 0.381 | 0.29 | 3.555e-29 | 6.343e-25 | Up in Hypoxia-high | Other |
| **OLFML2B** | -0.3574 | 0.498 | 0.596 | 4.244e-29 | 7.573e-25 | Up in Hypoxia-low | Other |
| **GM2A** | 0.1503 | 0.996 | 0.989 | 7.603e-29 | 1.357e-24 | Up in Hypoxia-high | Other |
| **AKT1S1** | 0.3621 | 0.603 | 0.537 | 8.188e-29 | 1.461e-24 | Up in Hypoxia-high | Other |
| **CTHRC1** | -0.4403 | 0.457 | 0.539 | 9.486e-29 | 1.693e-24 | Up in Hypoxia-low | Other |
| **TYMP** | 0.153 | 1 | 0.998 | 1.684e-28 | 3.005e-24 | Up in Hypoxia-high | Other |
| **CNN3** | -0.4417 | 0.354 | 0.458 | 1.743e-28 | 3.11e-24 | Up in Hypoxia-low | Other |

| | | | | | | | |
|---|---|---|---|---|---|---|---|
| **CTSD** | 0.1126 | 1 | 1 | 2.715e-28 | 4.844e-24 | Up in Hypoxia-high | Other |
| **LITAF** | 0.1815 | 0.996 | 0.993 | 5.842e-28 | 1.042e-23 | Up in Hypoxia-high | Other |
| **PTPN1** | 0.2335 | 0.932 | 0.901 | 6.8e-28 | 1.213e-23 | Up in Hypoxia-high | Other |
| **CXCL9** | 0.1151 | 0.992 | 0.986 | 7.276e-28 | 1.298e-23 | Up in Hypoxia-high | Other |
| **NIBAN2** | 0.361 | 0.638 | 0.573 | 7.53e-28 | 1.343e-23 | Up in Hypoxia-high | Other |
| **BNIP3L** | 0.2584 | 0.822 | 0.772 | 1.161e-27 | 2.072e-23 | Up in Hypoxia-high | Other |
| **GNS** | 0.2355 | 0.844 | 0.794 | 1.187e-27 | 2.118e-23 | Up in Hypoxia-high | Other |
| **TXNIP** | 0.2137 | 0.974 | 0.97 | 1.735e-27 | 3.096e-23 | Up in Hypoxia-high | Other |
| **TMEM132A** | 0.5455 | 0.346 | 0.251 | 2.299e-27 | 4.102e-23 | Up in Hypoxia-high | Other |
| **SYNE2** | -0.3195 | 0.532 | 0.627 | 2.383e-27 | 4.253e-23 | Up in Hypoxia-low | Other |
| **GSN** | 0.2292 | 0.98 | 0.968 | 2.402e-27 | 4.285e-23 | Up in Hypoxia-high | Other |
| **IFITM2** | -0.1996 | 0.966 | 0.978 | 2.709e-27 | 4.834e-23 | Up in Hypoxia-low | Other |
| **COL16A1** | -0.2999 | 0.628 | 0.683 | 4.56e-27 | 8.137e-23 | Up in Hypoxia-low | Other |
| **IL17RA** | 0.298 | 0.771 | 0.709 | 8.465e-27 | 1.51e-22 | Up in Hypoxia-high | Other |
| **UPP1** | 0.3543 | 0.564 | 0.494 | 1.574e-26 | 2.808e-22 | Up in Hypoxia-high | Other |
| **CYP27B1** | 0.5294 | 0.398 | 0.329 | 1.85e-26 | 3.302e-22 | Up in Hypoxia-high | Other |
| **PLEK** | 0.2093 | 0.928 | 0.905 | 3.088e-26 | 5.511e-22 | Up in Hypoxia-high | Other |
| **TNS1** | 0.3093 | 0.669 | 0.616 | 3.786e-26 | 6.756e-22 | Up in Hypoxia-high | Other |
| **SNRNP70** | -0.1726 | 0.915 | 0.948 | 4.676e-26 | 8.344e-22 | Up in Hypoxia-low | Other |
| **GPNMB** | 0.1404 | 0.992 | 0.977 | 5.059e-26 | 9.026e-22 | Up in Hypoxia-high | Other |
| **RNF149** | 0.1956 | 0.97 | 0.938 | 5.75e-26 | 1.026e-21 | Up in Hypoxia-high | Other |
| **LUM** | -0.1818 | 0.945 | 0.949 | 7.157e-26 | 1.277e-21 | Up in Hypoxia-low | Other |
| **FAM124A** | 0.5052 | 0.342 | 0.249 | 9.297e-26 | 1.659e-21 | Up in Hypoxia-high | Other |
| **QSOX1** | 0.3431 | 0.567 | 0.497 | 9.485e-26 | 1.692e-21 | Up in Hypoxia-high | Other |
| **IGFBP4** | -0.2309 | 0.795 | 0.806 | 1.179e-25 | 2.103e-21 | Up in Hypoxia-low | Other |
| **LGALS1** | 0.2007 | 0.939 | 0.905 | 1.511e-25 | 2.696e-21 | Up in Hypoxia-high | Other |
| **MYOF** | 0.2897 | 0.654 | 0.58 | 1.807e-25 | 3.225e-21 | Up in Hypoxia-high | Other |
| **HTRA1** | -0.4959 | 0.326 | 0.401 | 1.976e-25 | 3.526e-21 | Up in Hypoxia-low | Other |
| **HSD11B1** | 0.4234 | 0.579 | 0.509 | 2.823e-25 | 5.037e-21 | Up in Hypoxia-high | Other |
| **CTSZ** | 0.1129 | 1 | 1 | 2.023e-24 | 3.609e-20 | Up in Hypoxia-high | Other |
| **SERPINE1** | 0.2032 | 0.771 | 0.688 | 2.121e-24 | 3.785e-20 | Up in Hypoxia-high | Other |
| **PTMS** | -0.2122 | 0.881 | 0.914 | 2.25e-24 | 4.014e-20 | Up in Hypoxia-low | Other |
| **JUN** | 0.2926 | 0.727 | 0.687 | 2.999e-24 | 5.351e-20 | Up in Hypoxia-high | Other |
| **SEL1L3** | -0.2986 | 0.566 | 0.656 | 3.622e-24 | 6.462e-20 | Up in Hypoxia-low | Other |
| **VWF** | -0.3848 | 0.451 | 0.537 | 3.883e-24 | 6.928e-20 | Up in Hypoxia-low | Other |
| **HLA-DRB1** | 0.3324 | 0.467 | 0.44 | 6.741e-24 | 1.203e-19 | Up in Hypoxia-high | Other |
| **TNS3** | 0.2177 | 0.941 | 0.92 | 7.72e-24 | 1.378e-19 | Up in Hypoxia-high | Other |
| **LOXL2** | 0.3946 | 0.506 | 0.468 | 9.652e-24 | 1.722e-19 | Up in Hypoxia-high | Other |
| **ANXA2** | 0.2073 | 0.937 | 0.92 | 9.89e-24 | 1.765e-19 | Up in Hypoxia-high | Other |
| **NFKBIA** | 0.1472 | 0.994 | 0.986 | 9.983e-24 | 1.781e-19 | Up in Hypoxia-high | Other |
| **EPOP** | 0.5809 | 0.274 | 0.193 | 1.133e-23 | 2.021e-19 | Up in Hypoxia-high | Other |
| **NBL1** | -0.2998 | 0.769 | 0.806 | 2.008e-23 | 3.583e-19 | Up in Hypoxia-low | Other |
| **GK** | 0.596 | 0.265 | 0.182 | 2.282e-23 | 4.071e-19 | Up in Hypoxia-high | Other |
| **THEMIS2** | 0.1895 | 0.954 | 0.932 | 2.942e-23 | 5.25e-19 | Up in Hypoxia-high | Other |

| | | | | | | | |
|---|---|---|---|---|---|---|---|
| **LRRC15** | -0.4093 | 0.343 | 0.408 | 3.102e-23 | 5.535e-19 | Up in Hypoxia-low | Other |
| **PLVAP** | -0.4247 | 0.334 | 0.422 | 3.499e-23 | 6.244e-19 | Up in Hypoxia-low | Other |
| **IGHG3** | -0.3453 | 0.928 | 0.94 | 5.66e-23 | 1.01e-18 | Up in Hypoxia-low | Other |
| **SLAMF8** | 0.1828 | 0.956 | 0.927 | 7.541e-23 | 1.346e-18 | Up in Hypoxia-high | Other |
| **GOLM1** | -0.3238 | 0.539 | 0.608 | 8.38e-23 | 1.495e-18 | Up in Hypoxia-low | Other |
| **DPYSL3** | -0.332 | 0.514 | 0.578 | 1.289e-22 | 2.3e-18 | Up in Hypoxia-low | Other |
| **FGR** | 0.2328 | 0.887 | 0.845 | 1.332e-22 | 2.377e-18 | Up in Hypoxia-high | Other |
| **PTPRE** | 0.1872 | 0.895 | 0.853 | 2.008e-22 | 3.583e-18 | Up in Hypoxia-high | Other |
| **METRNL** | 0.236 | 0.845 | 0.799 | 2.504e-22 | 4.468e-18 | Up in Hypoxia-high | Other |
| **THBS1** | -0.439 | 0.305 | 0.39 | 2.759e-22 | 4.923e-18 | Up in Hypoxia-low | Other |
| **MCL1** | 0.1419 | 0.998 | 0.997 | 2.821e-22 | 5.033e-18 | Up in Hypoxia-high | Other |
| **SDC4** | 0.2682 | 0.815 | 0.753 | 3.148e-22 | 5.618e-18 | Up in Hypoxia-high | Other |
| **PALLD** | -0.3449 | 0.352 | 0.429 | 3.912e-22 | 6.98e-18 | Up in Hypoxia-low | Other |
| **SLC43A3** | 0.2791 | 0.682 | 0.617 | 5.241e-22 | 9.351e-18 | Up in Hypoxia-high | Other |
| **RARRES1** | -0.3346 | 0.377 | 0.423 | 6.812e-22 | 1.215e-17 | Up in Hypoxia-low | Other |
| **PFKP** | 0.2811 | 0.652 | 0.592 | 7.998e-22 | 1.427e-17 | Up in Hypoxia-high | Other |
| **CARD19** | 0.3575 | 0.505 | 0.427 | 9.589e-22 | 1.711e-17 | Up in Hypoxia-high | Other |
| **SH3BP5** | 0.2854 | 0.725 | 0.674 | 1.126e-21 | 2.009e-17 | Up in Hypoxia-high | Other |
| **IL1RN** | 0.5417 | 0.201 | 0.147 | 1.552e-21 | 2.768e-17 | Up in Hypoxia-high | Other |
| **FOXP1** | -0.3508 | 0.478 | 0.565 | 2.023e-21 | 3.609e-17 | Up in Hypoxia-low | Other |
| **ARRDC4** | 0.3929 | 0.473 | 0.407 | 2.309e-21 | 4.121e-17 | Up in Hypoxia-high | Other |
| **IFNGR2** | 0.1392 | 0.963 | 0.947 | 2.447e-21 | 4.366e-17 | Up in Hypoxia-high | Other |
| **TAPBP** | 0.1476 | 0.998 | 0.995 | 2.847e-21 | 5.079e-17 | Up in Hypoxia-high | Other |
| **FBLN2** | -0.2349 | 0.727 | 0.777 | 2.856e-21 | 5.097e-17 | Up in Hypoxia-low | Other |
| **KIF26B** | -0.514 | 0.212 | 0.274 | 3.079e-21 | 5.493e-17 | Up in Hypoxia-low | Other |
| **FCGR3A** | 0.1329 | 0.997 | 0.991 | 3.471e-21 | 6.194e-17 | Up in Hypoxia-high | Other |
| **SLC7A11** | 0.5406 | 0.302 | 0.218 | 3.838e-21 | 6.848e-17 | Up in Hypoxia-high | Other |
| **PDXK** | 0.2328 | 0.79 | 0.736 | 6.295e-21 | 1.123e-16 | Up in Hypoxia-high | Other |
| **GAS6** | -0.6097 | 0.168 | 0.24 | 6.404e-21 | 1.143e-16 | Up in Hypoxia-low | Other |
| **SDC3** | -0.2992 | 0.609 | 0.672 | 7.039e-21 | 1.256e-16 | Up in Hypoxia-low | Other |
| **COL4A1** | -0.1893 | 0.878 | 0.906 | 8.696e-21 | 1.552e-16 | Up in Hypoxia-low | Other |
| **LOXL1** | -0.3812 | 0.36 | 0.426 | 1.07e-20 | 1.909e-16 | Up in Hypoxia-low | Other |
| **BEX3** | -0.3652 | 0.39 | 0.474 | 1.137e-20 | 2.029e-16 | Up in Hypoxia-low | Other |
| **DENND5A** | 0.2396 | 0.769 | 0.715 | 1.423e-20 | 2.539e-16 | Up in Hypoxia-high | Other |
| **BCL2A1** | 0.3492 | 0.522 | 0.444 | 1.432e-20 | 2.556e-16 | Up in Hypoxia-high | Other |
| **EPAS1** | -0.324 | 0.487 | 0.558 | 1.917e-20 | 3.421e-16 | Up in Hypoxia-low | Other |
| **RILPL2** | 0.2246 | 0.838 | 0.775 | 2.162e-20 | 3.858e-16 | Up in Hypoxia-high | Other |
| **ANXA1** | 0.21 | 0.911 | 0.889 | 2.186e-20 | 3.901e-16 | Up in Hypoxia-high | Other |
| **IFI30** | 0.1033 | 1 | 1 | 2.211e-20 | 3.944e-16 | Up in Hypoxia-high | Other |
| **LILRB4** | 0.1894 | 0.88 | 0.827 | 2.248e-20 | 4.012e-16 | Up in Hypoxia-high | Other |
| **NFIB** | -0.4757 | 0.253 | 0.332 | 2.667e-20 | 4.759e-16 | Up in Hypoxia-low | Other |
| **CYBB** | 0.1127 | 0.999 | 0.998 | 2.991e-20 | 5.336e-16 | Up in Hypoxia-high | Other |
| **SHANK3** | -0.5276 | 0.196 | 0.26 | 3.092e-20 | 5.517e-16 | Up in Hypoxia-low | Other |
| **NRN1** | 0.5564 | 0.213 | 0.144 | 3.492e-20 | 6.23e-16 | Up in Hypoxia-high | Other |

| | | | | | | | |
|---|---|---|---|---|---|---|---|
| **PAPSS1** | 0.2416 | 0.817 | 0.767 | 3.498e-20 | 6.241e-16 | Up in Hypoxia-high | Other |
| **MFAP4** | -0.5088 | 0.255 | 0.319 | 3.58e-20 | 6.388e-16 | Up in Hypoxia-low | Other |
| **GRN** | 0.1145 | 0.998 | 0.993 | 3.917e-20 | 6.989e-16 | Up in Hypoxia-high | Other |
| **A2M** | -0.2715 | 0.671 | 0.732 | 6.648e-20 | 1.186e-15 | Up in Hypoxia-low | Other |
| **CDH11** | -0.3973 | 0.377 | 0.445 | 8.075e-20 | 1.441e-15 | Up in Hypoxia-low | Other |
| **RALA** | 0.2482 | 0.722 | 0.674 | 9.265e-20 | 1.653e-15 | Up in Hypoxia-high | Other |
| **C1orf122** | 0.2719 | 0.688 | 0.641 | 9.815e-20 | 1.751e-15 | Up in Hypoxia-high | Other |
| **LAP3** | 0.1133 | 0.999 | 0.998 | 1.105e-19 | 1.972e-15 | Up in Hypoxia-high | Other |
| **HBEGF** | 0.4578 | 0.336 | 0.275 | 1.232e-19 | 2.199e-15 | Up in Hypoxia-high | Other |
| **PRRX1** | -0.2918 | 0.553 | 0.628 | 1.597e-19 | 2.849e-15 | Up in Hypoxia-low | Other |
| **FOS** | -0.1963 | 0.736 | 0.796 | 1.626e-19 | 2.902e-15 | Up in Hypoxia-low | Other |
| **ENG** | -0.2082 | 0.842 | 0.877 | 1.979e-19 | 3.531e-15 | Up in Hypoxia-low | Other |
| **STAB1** | -0.4826 | 0.219 | 0.293 | 2.149e-19 | 3.835e-15 | Up in Hypoxia-low | Other |
| **ADAMTSL4** | 0.5092 | 0.287 | 0.209 | 2.159e-19 | 3.852e-15 | Up in Hypoxia-high | Other |
| **RNF130** | 0.2533 | 0.801 | 0.742 | 3.671e-19 | 6.551e-15 | Up in Hypoxia-high | Other |
| **SELENOP** | -0.5907 | 0.2 | 0.273 | 4.026e-19 | 7.184e-15 | Up in Hypoxia-low | Other |
| **NFAM1** | 0.2116 | 0.839 | 0.783 | 4.47e-19 | 7.976e-15 | Up in Hypoxia-high | Other |
| **CDKN1A** | 0.2804 | 0.654 | 0.612 | 5.03e-19 | 8.975e-15 | Up in Hypoxia-high | Other |
| **SELL** | -0.4642 | 0.301 | 0.382 | 5.312e-19 | 9.479e-15 | Up in Hypoxia-low | Other |
| **SIRPA** | 0.3319 | 0.536 | 0.465 | 5.5e-19 | 9.814e-15 | Up in Hypoxia-high | Other |
| **ITGB7** | 0.3273 | 0.453 | 0.397 | 5.573e-19 | 9.943e-15 | Up in Hypoxia-high | Other |
| **CFH** | -0.2417 | 0.688 | 0.747 | 6.329e-19 | 1.129e-14 | Up in Hypoxia-low | Other |
| **VWA1** | -0.3775 | 0.352 | 0.419 | 6.999e-19 | 1.249e-14 | Up in Hypoxia-low | Other |
| **EPHB4** | -0.5674 | 0.145 | 0.211 | 7.21e-19 | 1.287e-14 | Up in Hypoxia-low | Other |
| **HIST1H4A** | -0.2604 | 0.732 | 0.791 | 7.703e-19 | 1.374e-14 | Up in Hypoxia-low | Other |
| **SFRP4** | -0.6477 | 0.205 | 0.232 | 1.029e-18 | 1.836e-14 | Up in Hypoxia-low | Other |
| **HLA-DPA1** | 0.1108 | 1 | 1 | 1.095e-18 | 1.955e-14 | Up in Hypoxia-high | Other |
| **DERL3** | -0.4639 | 0.297 | 0.368 | 1.209e-18 | 2.157e-14 | Up in Hypoxia-low | Other |
| **CD79A** | -0.552 | 0.265 | 0.333 | 1.512e-18 | 2.697e-14 | Up in Hypoxia-low | Other |
| **AHNAK** | 0.236 | 0.852 | 0.814 | 1.8e-18 | 3.211e-14 | Up in Hypoxia-high | Other |
| **PIK3AP1** | 0.2197 | 0.668 | 0.619 | 1.822e-18 | 3.252e-14 | Up in Hypoxia-high | Other |
| **SLC9A3R2** | -0.4506 | 0.27 | 0.334 | 2.329e-18 | 4.156e-14 | Up in Hypoxia-low | Other |
| **IGLV3-1** | -0.5399 | 0.859 | 0.892 | 3.052e-18 | 5.445e-14 | Up in Hypoxia-low | Other |
| **HIST1H3D** | -0.2498 | 0.751 | 0.805 | 3.072e-18 | 5.481e-14 | Up in Hypoxia-low | Other |
| **RAMP3** | -0.5264 | 0.204 | 0.278 | 3.377e-18 | 6.025e-14 | Up in Hypoxia-low | Other |
| **PIM2** | -0.3645 | 0.41 | 0.482 | 3.697e-18 | 6.597e-14 | Up in Hypoxia-low | Other |
| **FCMR** | -0.459 | 0.306 | 0.389 | 3.881e-18 | 6.924e-14 | Up in Hypoxia-low | Other |
| **CCN1** | -0.2642 | 0.618 | 0.688 | 4.079e-18 | 7.279e-14 | Up in Hypoxia-low | Other |
| **SLC31A2** | 0.1871 | 0.88 | 0.828 | 4.222e-18 | 7.533e-14 | Up in Hypoxia-high | Other |
| **CORO1A** | -0.1591 | 0.93 | 0.954 | 4.348e-18 | 7.758e-14 | Up in Hypoxia-low | Other |
| **LGALS2** | 0.2121 | 0.823 | 0.782 | 7.195e-18 | 1.284e-13 | Up in Hypoxia-high | Other |
| **SH3BGRL3** | 0.1876 | 0.979 | 0.972 | 7.295e-18 | 1.302e-13 | Up in Hypoxia-high | Other |
| **TAP1** | 0.1089 | 0.998 | 0.996 | 8.194e-18 | 1.462e-13 | Up in Hypoxia-high | Other |
| **GXYLT2** | -0.5445 | 0.161 | 0.226 | 1.084e-17 | 1.934e-13 | Up in Hypoxia-low | Other |

| | | | | | | | |
|---|---|---|---|---|---|---|---|
| **CD63** | 0.1084 | 1 | 1 | 1.1e-17 | 1.963e-13 | Up in Hypoxia-high | Other |
| **CCDC115** | 0.2668 | 0.637 | 0.561 | 1.124e-17 | 2.005e-13 | Up in Hypoxia-high | Other |
| **CD27** | -0.4494 | 0.274 | 0.347 | 1.215e-17 | 2.167e-13 | Up in Hypoxia-low | Other |
| **ID3** | -0.4307 | 0.281 | 0.355 | 1.258e-17 | 2.245e-13 | Up in Hypoxia-low | Other |
| **SARDH** | 0.4732 | 0.305 | 0.245 | 1.297e-17 | 2.315e-13 | Up in Hypoxia-high | Other |
| **ANTXR1** | -0.3365 | 0.364 | 0.431 | 1.456e-17 | 2.598e-13 | Up in Hypoxia-low | Other |
| **CYBA** | 0.1223 | 0.998 | 0.998 | 1.538e-17 | 2.745e-13 | Up in Hypoxia-high | Other |
| **FCN1** | 0.4076 | 0.391 | 0.305 | 1.659e-17 | 2.961e-13 | Up in Hypoxia-high | Other |
| **WNT5A** | 0.4843 | 0.273 | 0.204 | 1.949e-17 | 3.478e-13 | Up in Hypoxia-high | Other |
| **CDH5** | -0.442 | 0.205 | 0.276 | 2.05e-17 | 3.658e-13 | Up in Hypoxia-low | Other |
| **COL15A1** | -0.3122 | 0.405 | 0.48 | 2.732e-17 | 4.876e-13 | Up in Hypoxia-low | Other |
| **IGLV6-57** | -0.5518 | 0.567 | 0.637 | 3.435e-17 | 6.128e-13 | Up in Hypoxia-low | Other |
| **HIST1H1B** | -0.2533 | 0.72 | 0.78 | 3.477e-17 | 6.203e-13 | Up in Hypoxia-low | Other |
| **MYO1G** | 0.2403 | 0.629 | 0.566 | 3.579e-17 | 6.385e-13 | Up in Hypoxia-high | Other |
| **KCNN4** | 0.5293 | 0.238 | 0.173 | 3.655e-17 | 6.522e-13 | Up in Hypoxia-high | Other |
| **RAB13** | 0.1603 | 0.943 | 0.91 | 4.028e-17 | 7.188e-13 | Up in Hypoxia-high | Other |
| **SDCBP** | 0.0953 | 0.995 | 0.992 | 4.251e-17 | 7.585e-13 | Up in Hypoxia-high | Other |
| **KYNU** | 0.1624 | 0.878 | 0.838 | 4.462e-17 | 7.961e-13 | Up in Hypoxia-high | Other |
| **LPCAT1** | -0.2404 | 0.545 | 0.624 | 5.473e-17 | 9.765e-13 | Up in Hypoxia-low | Other |
| **TMEM38B** | 0.3298 | 0.355 | 0.282 | 5.702e-17 | 1.017e-12 | Up in Hypoxia-high | Other |
| **HERPUD1** | -0.2041 | 0.817 | 0.86 | 5.718e-17 | 1.02e-12 | Up in Hypoxia-low | Other |
| **IL32** | 0.0793 | 1 | 1 | 6.109e-17 | 1.09e-12 | Up in Hypoxia-high | Other |
| **C1R** | -0.0801 | 0.995 | 0.997 | 6.268e-17 | 1.118e-12 | Up in Hypoxia-low | Other |
| **HLA-A** | 0.0563 | 0.99 | 0.989 | 6.669e-17 | 1.19e-12 | Up in Hypoxia-high | Other |
| **IGHJ2** | -0.5776 | 0.368 | 0.426 | 7.815e-17 | 1.394e-12 | Up in Hypoxia-low | Other |
| **DDX5** | -0.133 | 0.988 | 0.99 | 9.881e-17 | 1.763e-12 | Up in Hypoxia-low | Other |
| **CXCL16** | 0.2476 | 0.67 | 0.602 | 1.11e-16 | 1.981e-12 | Up in Hypoxia-high | Other |
| **PYCARD** | 0.1888 | 0.888 | 0.853 | 1.368e-16 | 2.44e-12 | Up in Hypoxia-high | Other |
| **LTBP2** | -0.6143 | 0.123 | 0.183 | 1.622e-16 | 2.894e-12 | Up in Hypoxia-low | Other |
| **ITGAX** | 0.1588 | 0.981 | 0.967 | 1.623e-16 | 2.896e-12 | Up in Hypoxia-high | Other |
| **AXL** | -0.3689 | 0.345 | 0.417 | 1.711e-16 | 3.052e-12 | Up in Hypoxia-low | Other |
| **IGHD** | -0.6421 | 0.274 | 0.322 | 1.85e-16 | 3.302e-12 | Up in Hypoxia-low | Other |
| **CD53** | 0.1187 | 0.994 | 0.991 | 1.892e-16 | 3.376e-12 | Up in Hypoxia-high | Other |
| **SCPEP1** | 0.1264 | 0.989 | 0.983 | 1.929e-16 | 3.442e-12 | Up in Hypoxia-high | Other |
| **PLEKHB2** | 0.227 | 0.77 | 0.729 | 2.072e-16 | 3.697e-12 | Up in Hypoxia-high | Other |
| **TNF** | 0.4124 | 0.331 | 0.279 | 2.39e-16 | 4.265e-12 | Up in Hypoxia-high | Other |
| **SLCO2B1** | -0.2586 | 0.532 | 0.61 | 2.391e-16 | 4.266e-12 | Up in Hypoxia-low | Other |
| **PODN** | -0.6003 | 0.128 | 0.184 | 2.605e-16 | 4.648e-12 | Up in Hypoxia-low | Other |
| **IGLJ1** | -0.507 | 0.178 | 0.251 | 4.935e-16 | 8.806e-12 | Up in Hypoxia-low | Other |
| **HEXB** | 0.2034 | 0.818 | 0.781 | 5.784e-16 | 1.032e-11 | Up in Hypoxia-high | Other |
| **COL18A1** | -0.2485 | 0.536 | 0.581 | 5.812e-16 | 1.037e-11 | Up in Hypoxia-low | Other |
| **CD93** | -0.3162 | 0.356 | 0.413 | 7.311e-16 | 1.304e-11 | Up in Hypoxia-low | Other |
| **TTC3** | -0.2411 | 0.612 | 0.681 | 7.415e-16 | 1.323e-11 | Up in Hypoxia-low | Other |
| **FXYD2** | 0.4854 | 0.249 | 0.184 | 7.519e-16 | 1.342e-11 | Up in Hypoxia-high | Other |

| | | | | | | | |
|---|---|---|---|---|---|---|---|
| **DPT** | -0.2675 | 0.418 | 0.439 | 8.03e-16 | 1.433e-11 | Up in Hypoxia-low | Other |
| **FMOD** | -0.3207 | 0.324 | 0.385 | 9.754e-16 | 1.74e-11 | Up in Hypoxia-low | Other |
| **GBE1** | 0.6348 | 0.165 | 0.107 | 1.138e-15 | 2.031e-11 | Up in Hypoxia-high | Other |
| **GBP1** | 0.1372 | 0.881 | 0.864 | 1.209e-15 | 2.158e-11 | Up in Hypoxia-high | Other |
| **ADAM8** | 0.5137 | 0.219 | 0.154 | 1.318e-15 | 2.351e-11 | Up in Hypoxia-high | Other |
| **COX5B** | 0.1691 | 0.951 | 0.933 | 1.329e-15 | 2.372e-11 | Up in Hypoxia-high | Other |
| **HIST1H2BD** | -0.2714 | 0.515 | 0.596 | 1.414e-15 | 2.524e-11 | Up in Hypoxia-low | Other |
| **ABCG1** | 0.2248 | 0.681 | 0.642 | 1.461e-15 | 2.607e-11 | Up in Hypoxia-high | Other |
| **HIST1H4D** | -0.2261 | 0.785 | 0.833 | 1.464e-15 | 2.612e-11 | Up in Hypoxia-low | Other |
| **HIST1H4C** | -0.2664 | 0.727 | 0.781 | 1.504e-15 | 2.684e-11 | Up in Hypoxia-low | Other |
| **SDC1** | -0.5457 | 0.148 | 0.201 | 1.882e-15 | 3.357e-11 | Up in Hypoxia-low | Other |
| **LHFPL6** | -0.4872 | 0.204 | 0.265 | 1.91e-15 | 3.407e-11 | Up in Hypoxia-low | Other |
| **FSTL1** | -0.1489 | 0.899 | 0.918 | 2.203e-15 | 3.931e-11 | Up in Hypoxia-low | Other |
| **RNASE1** | 0.4022 | 0.41 | 0.359 | 3.268e-15 | 5.831e-11 | Up in Hypoxia-high | Other |
| **LGALS3BP** | 0.1207 | 0.997 | 0.994 | 3.692e-15 | 6.587e-11 | Up in Hypoxia-high | Other |
| **MYLK** | -0.3797 | 0.326 | 0.397 | 3.942e-15 | 7.033e-11 | Up in Hypoxia-low | Other |
| **C11orf96** | 0.36 | 0.374 | 0.336 | 4.01e-15 | 7.155e-11 | Up in Hypoxia-high | Other |
| **NOL3** | 0.5959 | 0.185 | 0.13 | 4.063e-15 | 7.25e-11 | Up in Hypoxia-high | Other |
| **LRRK2** | 0.208 | 0.639 | 0.601 | 4.232e-15 | 7.551e-11 | Up in Hypoxia-high | Other |
| **LAMA4** | -0.2034 | 0.713 | 0.758 | 4.345e-15 | 7.753e-11 | Up in Hypoxia-low | Other |
| **AP5B1** | 0.1755 | 0.81 | 0.746 | 4.385e-15 | 7.824e-11 | Up in Hypoxia-high | Other |
| **SPI1** | 0.1105 | 0.991 | 0.988 | 4.429e-15 | 7.902e-11 | Up in Hypoxia-high | Other |
| **OLFML3** | -0.4416 | 0.181 | 0.239 | 4.454e-15 | 7.947e-11 | Up in Hypoxia-low | Other |
| **PTGIS** | -0.555 | 0.139 | 0.19 | 4.463e-15 | 7.964e-11 | Up in Hypoxia-low | Other |
| **FBLIM1** | -0.29 | 0.375 | 0.442 | 4.554e-15 | 8.126e-11 | Up in Hypoxia-low | Other |
| **P4HB** | 0.1769 | 0.96 | 0.94 | 5.605e-15 | 1E-10 | Up in Hypoxia-high | Other |
| **FPR2** | 0.5073 | 0.242 | 0.178 | 5.612e-15 | 1.001e-10 | Up in Hypoxia-high | Other |
| **CPNE5** | -0.5141 | 0.128 | 0.189 | 5.677e-15 | 1.013e-10 | Up in Hypoxia-low | Other |
| **CRIP2** | -0.3333 | 0.354 | 0.413 | 5.979e-15 | 1.067e-10 | Up in Hypoxia-low | Other |
| **PSME2** | 0.1203 | 0.989 | 0.984 | 6.049e-15 | 1.079e-10 | Up in Hypoxia-high | Other |
| **NPC2** | 0.1053 | 0.995 | 0.995 | 8.441e-15 | 1.506e-10 | Up in Hypoxia-high | Other |
| **CTSH** | 0.0941 | 0.991 | 0.988 | 9.681e-15 | 1.727e-10 | Up in Hypoxia-high | Other |
| **NR2F2** | -0.4276 | 0.235 | 0.3 | 9.704e-15 | 1.732e-10 | Up in Hypoxia-low | Other |
| **SOX18** | -0.4361 | 0.197 | 0.256 | 1.18e-14 | 2.106e-10 | Up in Hypoxia-low | Other |
| **CLDN7** | 0.4222 | 0.273 | 0.205 | 1.238e-14 | 2.209e-10 | Up in Hypoxia-high | Other |
| **CSF2RB** | 0.1452 | 0.913 | 0.879 | 1.276e-14 | 2.278e-10 | Up in Hypoxia-high | Other |
| **PCDH7** | -0.4885 | 0.128 | 0.182 | 1.387e-14 | 2.474e-10 | Up in Hypoxia-low | Other |
| **PRKCD** | 0.2253 | 0.7 | 0.643 | 1.815e-14 | 3.239e-10 | Up in Hypoxia-high | Other |
| **VASH1** | -0.1621 | 0.865 | 0.887 | 1.838e-14 | 3.28e-10 | Up in Hypoxia-low | Other |
| **CISH** | 0.3673 | 0.348 | 0.284 | 1.861e-14 | 3.32e-10 | Up in Hypoxia-high | Other |
| **F2R** | -0.2982 | 0.359 | 0.425 | 1.968e-14 | 3.512e-10 | Up in Hypoxia-low | Other |
| **PRNP** | 0.1888 | 0.813 | 0.771 | 1.993e-14 | 3.556e-10 | Up in Hypoxia-high | Other |
| **CNN2** | -0.2409 | 0.637 | 0.689 | 2.025e-14 | 3.613e-10 | Up in Hypoxia-low | Other |
| **TAGLN2** | 0.1205 | 0.999 | 0.998 | 2.052e-14 | 3.661e-10 | Up in Hypoxia-high | Other |

| | | | | | | | |
|---|---|---|---|---|---|---|---|
| **SYNPO** | -0.4884 | 0.17 | 0.232 | 2.169e-14 | 3.869e-10 | Up in Hypoxia-low | Other |
| **FNDC1** | -0.4292 | 0.167 | 0.22 | 2.473e-14 | 4.413e-10 | Up in Hypoxia-low | Other |
| **INHBA** | 0.2595 | 0.744 | 0.71 | 3.237e-14 | 5.776e-10 | Up in Hypoxia-high | Other |
| **PSTPIP2** | 0.2675 | 0.51 | 0.448 | 3.363e-14 | 6E-10 | Up in Hypoxia-high | Other |
| **SLC40A1** | -0.6673 | 0.111 | 0.169 | 3.693e-14 | 6.589e-10 | Up in Hypoxia-low | Other |
| **IKZF1** | -0.1687 | 0.807 | 0.856 | 4.156e-14 | 7.415e-10 | Up in Hypoxia-low | Other |
| **LBH** | -0.1994 | 0.743 | 0.787 | 4.431e-14 | 7.906e-10 | Up in Hypoxia-low | Other |
| **FGL2** | 0.1611 | 0.935 | 0.938 | 4.693e-14 | 8.373e-10 | Up in Hypoxia-high | Other |
| **HSPB1** | -0.1188 | 0.985 | 0.989 | 5.1e-14 | 9.1e-10 | Up in Hypoxia-low | Other |
| **PXDN** | -0.3119 | 0.376 | 0.44 | 5.585e-14 | 9.966e-10 | Up in Hypoxia-low | Other |
| **HNRNPA3** | -0.136 | 0.935 | 0.958 | 5.736e-14 | 1.024e-09 | Up in Hypoxia-low | Other |
| **ACKR1** | -0.6194 | 0.133 | 0.189 | 6.649e-14 | 1.186e-09 | Up in Hypoxia-low | Other |
| **PLIN2** | 0.2889 | 0.516 | 0.464 | 7.606e-14 | 1.357e-09 | Up in Hypoxia-high | Other |
| **ABCA1** | 0.2324 | 0.48 | 0.429 | 1.172e-13 | 2.091e-09 | Up in Hypoxia-high | Other |
| **PIK3CB** | 0.2948 | 0.394 | 0.335 | 1.173e-13 | 2.092e-09 | Up in Hypoxia-high | Other |
| **SPNS2** | -0.6594 | 0.1 | 0.15 | 1.453e-13 | 2.593e-09 | Up in Hypoxia-low | Other |
| **TNFSF13** | 0.1795 | 0.849 | 0.802 | 1.492e-13 | 2.662e-09 | Up in Hypoxia-high | Other |
| **JAML** | 0.232 | 0.636 | 0.58 | 1.552e-13 | 2.768e-09 | Up in Hypoxia-high | Other |
| **ANGPTL2** | -0.2829 | 0.383 | 0.451 | 1.667e-13 | 2.974e-09 | Up in Hypoxia-low | Other |
| **LAMP1** | 0.1292 | 0.979 | 0.974 | 1.73e-13 | 3.088e-09 | Up in Hypoxia-high | Other |
| **LIMD2** | -0.1354 | 0.947 | 0.962 | 1.794e-13 | 3.201e-09 | Up in Hypoxia-low | Other |
| **ALDOC** | 0.515 | 0.207 | 0.15 | 1.794e-13 | 3.201e-09 | Up in Hypoxia-high | Other |
| **ANKRD22** | 0.1766 | 0.663 | 0.612 | 1.816e-13 | 3.241e-09 | Up in Hypoxia-high | Other |
| **ZCCHC24** | -0.3533 | 0.284 | 0.344 | 1.89e-13 | 3.372e-09 | Up in Hypoxia-low | Other |
| **SLC2A5** | 0.3523 | 0.382 | 0.314 | 1.97e-13 | 3.516e-09 | Up in Hypoxia-high | Other |
| **IRF1** | 0.1021 | 0.993 | 0.992 | 1.978e-13 | 3.53e-09 | Up in Hypoxia-high | Other |
| **EMILIN2** | 0.236 | 0.637 | 0.57 | 2.01e-13 | 3.587e-09 | Up in Hypoxia-high | Other |
| **SELE** | -0.6357 | 0.127 | 0.179 | 2.106e-13 | 3.759e-09 | Up in Hypoxia-low | Other |
| **C3AR1** | 0.5295 | 0.198 | 0.139 | 2.199e-13 | 3.923e-09 | Up in Hypoxia-high | Other |
| **C1S** | -0.1333 | 0.954 | 0.96 | 2.206e-13 | 3.936e-09 | Up in Hypoxia-low | Other |
| **S100A9** | 0.1395 | 0.954 | 0.933 | 2.423e-13 | 4.323e-09 | Up in Hypoxia-high | Other |
| **SSR4** | -0.1986 | 0.862 | 0.886 | 2.424e-13 | 4.326e-09 | Up in Hypoxia-low | Other |
| **C5AR1** | 0.2147 | 0.689 | 0.625 | 2.511e-13 | 4.481e-09 | Up in Hypoxia-high | Other |
| **MXI1** | 0.3578 | 0.35 | 0.285 | 2.86e-13 | 5.102e-09 | Up in Hypoxia-high | Other |
| **SIGLEC10** | 0.1625 | 0.865 | 0.821 | 3.03e-13 | 5.406e-09 | Up in Hypoxia-high | Other |
| **ICAM2** | -0.3817 | 0.285 | 0.356 | 3.083e-13 | 5.5e-09 | Up in Hypoxia-low | Other |
| **PTP4A3** | -0.4649 | 0.149 | 0.209 | 3.326e-13 | 5.935e-09 | Up in Hypoxia-low | Other |
| **ADAMTSL2** | -0.4872 | 0.148 | 0.191 | 3.484e-13 | 6.217e-09 | Up in Hypoxia-low | Other |
| **SPON2** | -0.2227 | 0.365 | 0.408 | 3.585e-13 | 6.397e-09 | Up in Hypoxia-low | Other |
| **TNIP1** | 0.1583 | 0.858 | 0.832 | 4.017e-13 | 7.167e-09 | Up in Hypoxia-high | Other |
| **MCAM** | -0.3389 | 0.29 | 0.359 | 4.34e-13 | 7.744e-09 | Up in Hypoxia-low | Other |
| **CD274** | 0.3089 | 0.374 | 0.318 | 4.721e-13 | 8.424e-09 | Up in Hypoxia-high | Other |
| **MSC** | 0.3536 | 0.375 | 0.316 | 4.922e-13 | 8.782e-09 | Up in Hypoxia-high | Other |
| **TSPO** | 0.1103 | 0.957 | 0.929 | 5.116e-13 | 9.128e-09 | Up in Hypoxia-high | Other |

| | | | | | | | |
|---|---|---|---|---|---|---|---|
| **EEF2** | -0.0552 | 1 | 1 | 5.143e-13 | 9.177e-09 | Up in Hypoxia-low | Other |
| **IFITM1** | -0.2361 | 0.73 | 0.777 | 5.978e-13 | 1.067e-08 | Up in Hypoxia-low | Other |
| **SLC22A3** | -0.527 | 0.155 | 0.208 | 6.037e-13 | 1.077e-08 | Up in Hypoxia-low | Other |
| **DRAM1** | 0.1672 | 0.757 | 0.728 | 6.411e-13 | 1.144e-08 | Up in Hypoxia-high | Other |
| **SYNPO2** | -0.6065 | 0.11 | 0.161 | 7.016e-13 | 1.252e-08 | Up in Hypoxia-low | Other |
| **ARGLU1** | -0.1513 | 0.857 | 0.888 | 7.365e-13 | 1.314e-08 | Up in Hypoxia-low | Other |
| **TFRC** | 0.1227 | 0.963 | 0.935 | 7.67e-13 | 1.369e-08 | Up in Hypoxia-high | Other |
| **NNMT** | -0.1535 | 0.79 | 0.815 | 8.395e-13 | 1.498e-08 | Up in Hypoxia-low | Other |
| **ACOT7** | 0.2625 | 0.616 | 0.577 | 9.207e-13 | 1.643e-08 | Up in Hypoxia-high | Other |
| **FCAR** | 0.4235 | 0.213 | 0.156 | 1.026e-12 | 1.832e-08 | Up in Hypoxia-high | Other |
| **TENT5C** | -0.3506 | 0.319 | 0.379 | 1.086e-12 | 1.937e-08 | Up in Hypoxia-low | Other |
| **CXCL13** | -0.4015 | 0.376 | 0.411 | 1.151e-12 | 2.054e-08 | Up in Hypoxia-low | Other |
| **IL1R1** | -0.2986 | 0.359 | 0.436 | 1.157e-12 | 2.064e-08 | Up in Hypoxia-low | Other |
| **LAMB3** | 0.4302 | 0.203 | 0.147 | 1.229e-12 | 2.193e-08 | Up in Hypoxia-high | Other |
| **BCAP31** | 0.1636 | 0.919 | 0.897 | 1.349e-12 | 2.407e-08 | Up in Hypoxia-high | Other |
| **FSTL3** | 0.4825 | 0.242 | 0.184 | 1.564e-12 | 2.79e-08 | Up in Hypoxia-high | Other |
| **CPXM1** | -0.3552 | 0.23 | 0.284 | 1.656e-12 | 2.955e-08 | Up in Hypoxia-low | Other |
| **ITGA4** | -0.356 | 0.328 | 0.402 | 1.681e-12 | 3E-08 | Up in Hypoxia-low | Other |
| **MYH9** | -0.0956 | 0.998 | 0.999 | 1.752e-12 | 3.125e-08 | Up in Hypoxia-low | Other |
| **HIC1** | -0.2725 | 0.283 | 0.313 | 1.855e-12 | 3.31e-08 | Up in Hypoxia-low | Other |
| **SLC38A1** | -0.3628 | 0.332 | 0.401 | 1.899e-12 | 3.389e-08 | Up in Hypoxia-low | Other |
| **MDFI** | -0.3745 | 0.197 | 0.248 | 1.9e-12 | 3.39e-08 | Up in Hypoxia-low | Other |
| **ADA2** | 0.1361 | 0.971 | 0.959 | 1.939e-12 | 3.46e-08 | Up in Hypoxia-high | Other |
| **ZFAND5** | 0.1532 | 0.949 | 0.926 | 2.144e-12 | 3.826e-08 | Up in Hypoxia-high | Other |
| **MDK** | -0.5235 | 0.154 | 0.212 | 2.203e-12 | 3.932e-08 | Up in Hypoxia-low | Other |
| **SPIB** | -0.4683 | 0.208 | 0.261 | 2.474e-12 | 4.414e-08 | Up in Hypoxia-low | Other |
| **SULT1C2** | 0.2442 | 0.564 | 0.491 | 2.627e-12 | 4.688e-08 | Up in Hypoxia-high | Other |
| **AC119396.1** | -0.5028 | 0.133 | 0.189 | 2.672e-12 | 4.768e-08 | Up in Hypoxia-low | Other |
| **C1orf198** | -0.3985 | 0.192 | 0.251 | 3.296e-12 | 5.88e-08 | Up in Hypoxia-low | Other |
| **ATP6V1B2** | 0.0862 | 0.993 | 0.987 | 3.516e-12 | 6.274e-08 | Up in Hypoxia-high | Other |
| **ANXA6** | -0.2083 | 0.566 | 0.614 | 3.632e-12 | 6.48e-08 | Up in Hypoxia-low | Other |
| **CIRBP** | -0.1248 | 0.979 | 0.985 | 3.837e-12 | 6.846e-08 | Up in Hypoxia-low | Other |
| **PTGES** | 0.4513 | 0.24 | 0.185 | 4.045e-12 | 7.218e-08 | Up in Hypoxia-high | Other |
| **NISCH** | -0.2679 | 0.415 | 0.477 | 4.345e-12 | 7.754e-08 | Up in Hypoxia-low | Other |
| **ETS1** | -0.1772 | 0.769 | 0.806 | 4.484e-12 | 8.001e-08 | Up in Hypoxia-low | Other |
| **PHLDB1** | -0.2077 | 0.528 | 0.565 | 4.777e-12 | 8.523e-08 | Up in Hypoxia-low | Other |
| **MS4A1** | -0.5711 | 0.182 | 0.234 | 4.987e-12 | 8.898e-08 | Up in Hypoxia-low | Other |
| **CD300E** | 0.4391 | 0.241 | 0.182 | 5.256e-12 | 9.379e-08 | Up in Hypoxia-high | Other |
| **RNF13** | 0.1369 | 0.753 | 0.704 | 5.553e-12 | 9.909e-08 | Up in Hypoxia-high | Other |
| **GUCY1A1** | -0.3744 | 0.223 | 0.288 | 5.929e-12 | 1.058e-07 | Up in Hypoxia-low | Other |
| **RAPGEF1** | 0.1335 | 0.873 | 0.839 | 5.981e-12 | 1.067e-07 | Up in Hypoxia-high | Other |
| **ADAMDEC1** | 0.1843 | 0.694 | 0.647 | 6.353e-12 | 1.134e-07 | Up in Hypoxia-high | Other |
| **SERPING1** | 0.0946 | 0.985 | 0.978 | 6.543e-12 | 1.167e-07 | Up in Hypoxia-high | Other |
| **LXN** | -0.4198 | 0.195 | 0.239 | 6.543e-12 | 1.167e-07 | Up in Hypoxia-low | Other |

| | | | | | | | |
|---|---|---|---|---|---|---|---|
| **TLE5** | -0.1398 | 0.944 | 0.963 | 6.707e-12 | 1.197e-07 | Up in Hypoxia-low | Other |
| **NR1H3** | 0.0972 | 0.963 | 0.954 | 7.502e-12 | 1.339e-07 | Up in Hypoxia-high | Other |
| **SOD3** | -0.183 | 0.612 | 0.643 | 7.676e-12 | 1.37e-07 | Up in Hypoxia-low | Other |
| **TRBC1** | -0.1769 | 0.785 | 0.827 | 7.823e-12 | 1.396e-07 | Up in Hypoxia-low | Other |
| **CYP2S1** | 0.3248 | 0.411 | 0.351 | 7.913e-12 | 1.412e-07 | Up in Hypoxia-high | Other |
| **TRDC** | -0.3932 | 0.219 | 0.287 | 8.263e-12 | 1.474e-07 | Up in Hypoxia-low | Other |
| **CXCR4** | -0.1437 | 0.823 | 0.852 | 8.956e-12 | 1.598e-07 | Up in Hypoxia-low | Other |
| **ITGA11** | -0.4516 | 0.118 | 0.168 | 1.022e-11 | 1.824e-07 | Up in Hypoxia-low | Other |
| **MARCKSL1** | -0.2238 | 0.574 | 0.617 | 1.053e-11 | 1.878e-07 | Up in Hypoxia-low | Other |
| **COL12A1** | -0.45 | 0.192 | 0.243 | 1.191e-11 | 2.126e-07 | Up in Hypoxia-low | Other |
| **TKT** | 0.1046 | 0.984 | 0.973 | 1.219e-11 | 2.176e-07 | Up in Hypoxia-high | Other |
| **TSPAN13** | -0.5418 | 0.137 | 0.192 | 1.226e-11 | 2.188e-07 | Up in Hypoxia-low | Other |
| **IGFLR1** | 0.1831 | 0.699 | 0.66 | 1.293e-11 | 2.308e-07 | Up in Hypoxia-high | Other |
| **TSPAN3** | 0.1673 | 0.759 | 0.711 | 1.357e-11 | 2.422e-07 | Up in Hypoxia-high | Other |
| **COL14A1** | -0.4768 | 0.124 | 0.169 | 1.446e-11 | 2.58e-07 | Up in Hypoxia-low | Other |
| **CTBS** | 0.4193 | 0.251 | 0.191 | 1.543e-11 | 2.753e-07 | Up in Hypoxia-high | Other |
| **GLIS2** | -0.404 | 0.189 | 0.244 | 1.817e-11 | 3.243e-07 | Up in Hypoxia-low | Other |
| **HNRNPDL** | -0.0958 | 0.999 | 0.997 | 1.852e-11 | 3.305e-07 | Up in Hypoxia-low | Other |
| **C1QTNF6** | -0.2818 | 0.302 | 0.36 | 1.894e-11 | 3.379e-07 | Up in Hypoxia-low | Other |
| **NRIP3** | 0.5383 | 0.147 | 0.101 | 2.007e-11 | 3.58e-07 | Up in Hypoxia-high | Other |
| **TM4SF1** | -0.5063 | 0.139 | 0.193 | 2.036e-11 | 3.634e-07 | Up in Hypoxia-low | Other |
| **NCSTN** | 0.1602 | 0.723 | 0.673 | 2.221e-11 | 3.963e-07 | Up in Hypoxia-high | Other |
| **MXRA8** | -0.2071 | 0.392 | 0.423 | 2.392e-11 | 4.269e-07 | Up in Hypoxia-low | Other |
| **MGST3** | 0.2096 | 0.694 | 0.654 | 2.505e-11 | 4.469e-07 | Up in Hypoxia-high | Other |
| **TNXB** | -0.5269 | 0.116 | 0.162 | 2.684e-11 | 4.788e-07 | Up in Hypoxia-low | Other |
| **SGK1** | 0.1242 | 0.93 | 0.907 | 2.698e-11 | 4.815e-07 | Up in Hypoxia-high | Other |
| **SCD** | 0.1467 | 0.79 | 0.759 | 3.019e-11 | 5.386e-07 | Up in Hypoxia-high | Other |
| **SHC1** | -0.2808 | 0.418 | 0.485 | 3.075e-11 | 5.486e-07 | Up in Hypoxia-low | Other |
| **PGF** | -0.4645 | 0.148 | 0.196 | 3.231e-11 | 5.766e-07 | Up in Hypoxia-low | Other |
| **NDP** | 0.5581 | 0.152 | 0.104 | 3.299e-11 | 5.886e-07 | Up in Hypoxia-high | Other |
| **ATP6AP1** | 0.1283 | 0.899 | 0.86 | 3.421e-11 | 6.104e-07 | Up in Hypoxia-high | Other |
| **CD79B** | -0.5284 | 0.168 | 0.219 | 3.43e-11 | 6.121e-07 | Up in Hypoxia-low | Other |
| **SEPTIN6** | -0.2387 | 0.509 | 0.571 | 3.732e-11 | 6.66e-07 | Up in Hypoxia-low | Other |
| **SCARF1** | 0.2152 | 0.604 | 0.54 | 3.891e-11 | 6.943e-07 | Up in Hypoxia-high | Other |
| **WFDC2** | 0.4578 | 0.207 | 0.155 | 4.015e-11 | 7.163e-07 | Up in Hypoxia-high | Other |
| **LRRC25** | 0.3152 | 0.386 | 0.32 | 4.194e-11 | 7.483e-07 | Up in Hypoxia-high | Other |
| **ANXA5** | 0.114 | 0.975 | 0.969 | 4.818e-11 | 8.598e-07 | Up in Hypoxia-high | Other |
| **PRSS23** | -0.2351 | 0.433 | 0.47 | 5.102e-11 | 9.103e-07 | Up in Hypoxia-low | Other |
| **UCP2** | -0.1859 | 0.773 | 0.815 | 5.594e-11 | 9.982e-07 | Up in Hypoxia-low | Other |
| **SELENOM** | -0.1428 | 0.788 | 0.826 | 5.759e-11 | 1.028e-06 | Up in Hypoxia-low | Other |
| **CCL3L1** | 0.3818 | 0.26 | 0.207 | 5.76e-11 | 1.028e-06 | Up in Hypoxia-high | Other |
| **APBB2** | -0.5051 | 0.14 | 0.191 | 5.775e-11 | 1.03e-06 | Up in Hypoxia-low | Other |
| **CLEC14A** | -0.49 | 0.146 | 0.194 | 5.937e-11 | 1.059e-06 | Up in Hypoxia-low | Other |
| **PECAM1** | -0.1768 | 0.732 | 0.782 | 6.645e-11 | 1.186e-06 | Up in Hypoxia-low | Other |

| | | | | | | | |
|---|---|---|---|---|---|---|---|
| **PPARGC1B** | 0.3104 | 0.358 | 0.295 | 6.841e-11 | 1.221e-06 | Up in Hypoxia-high | Other |
| **CTSL** | 0.1562 | 0.88 | 0.854 | 6.851e-11 | 1.222e-06 | Up in Hypoxia-high | Other |
| **PDGFRB** | -0.1649 | 0.694 | 0.733 | 7.135e-11 | 1.273e-06 | Up in Hypoxia-low | Other |
| **IL7R** | -0.2078 | 0.456 | 0.511 | 8.281e-11 | 1.478e-06 | Up in Hypoxia-low | Other |
| **EEF1B2** | -0.1312 | 0.962 | 0.968 | 8.852e-11 | 1.58e-06 | Up in Hypoxia-low | Other |
| **COL7A1** | 0.389 | 0.204 | 0.155 | 9.655e-11 | 1.723e-06 | Up in Hypoxia-high | Other |
| **ATP6V0E1** | 0.1713 | 0.793 | 0.75 | 9.872e-11 | 1.761e-06 | Up in Hypoxia-high | Other |
| **SLC5A3** | 0.1809 | 0.472 | 0.427 | 1.012e-10 | 1.806e-06 | Up in Hypoxia-high | Other |
| **SNX29** | -0.247 | 0.4 | 0.463 | 1.078e-10 | 1.923e-06 | Up in Hypoxia-low | Other |
| **PRRC2B** | -0.1288 | 0.902 | 0.923 | 1.087e-10 | 1.939e-06 | Up in Hypoxia-low | Other |
| **PLS3** | -0.354 | 0.217 | 0.269 | 1.155e-10 | 2.061e-06 | Up in Hypoxia-low | Other |
| **BHLHE41** | 0.3156 | 0.313 | 0.256 | 1.169e-10 | 2.086e-06 | Up in Hypoxia-high | Other |
| **HIST2H2AB** | -0.2771 | 0.463 | 0.523 | 1.219e-10 | 2.174e-06 | Up in Hypoxia-low | Other |
| **ORMDL3** | -0.2248 | 0.527 | 0.582 | 1.261e-10 | 2.25e-06 | Up in Hypoxia-low | Other |
| **NOP53** | -0.1321 | 0.95 | 0.966 | 1.307e-10 | 2.332e-06 | Up in Hypoxia-low | Other |
| **KIAA1217** | -0.4113 | 0.157 | 0.208 | 1.395e-10 | 2.489e-06 | Up in Hypoxia-low | Other |
| **ACTG2** | -0.4639 | 0.132 | 0.181 | 1.519e-10 | 2.71e-06 | Up in Hypoxia-low | Other |
| **PARVA** | -0.4024 | 0.173 | 0.218 | 1.763e-10 | 3.146e-06 | Up in Hypoxia-low | Other |
| **EIF4B** | -0.1129 | 0.971 | 0.983 | 1.823e-10 | 3.253e-06 | Up in Hypoxia-low | Other |
| **BTG2** | -0.1751 | 0.684 | 0.721 | 1.921e-10 | 3.428e-06 | Up in Hypoxia-low | Other |
| **UGP2** | 0.2212 | 0.59 | 0.543 | 1.971e-10 | 3.517e-06 | Up in Hypoxia-high | Other |
| **CRELD2** | -0.1944 | 0.595 | 0.642 | 1.998e-10 | 3.565e-06 | Up in Hypoxia-low | Other |
| **CD200** | -0.3853 | 0.169 | 0.225 | 2.054e-10 | 3.665e-06 | Up in Hypoxia-low | Other |
| **CLEC12A** | 0.4279 | 0.212 | 0.16 | 2.145e-10 | 3.827e-06 | Up in Hypoxia-high | Other |
| **COX4I1** | 0.109 | 0.98 | 0.976 | 2.169e-10 | 3.87e-06 | Up in Hypoxia-high | Other |
| **IQGAP1** | 0.1 | 0.961 | 0.948 | 2.355e-10 | 4.203e-06 | Up in Hypoxia-high | Other |
| **CLPTM1L** | -0.1705 | 0.717 | 0.759 | 2.363e-10 | 4.216e-06 | Up in Hypoxia-low | Other |
| **GNG11** | -0.509 | 0.113 | 0.157 | 2.367e-10 | 4.224e-06 | Up in Hypoxia-low | Other |
| **CSF1R** | 0.1333 | 0.976 | 0.965 | 2.378e-10 | 4.242e-06 | Up in Hypoxia-high | Other |
| **HIST1H3H** | -0.3022 | 0.468 | 0.516 | 2.497e-10 | 4.456e-06 | Up in Hypoxia-low | Other |
| **GYS1** | 0.3368 | 0.31 | 0.25 | 2.568e-10 | 4.583e-06 | Up in Hypoxia-high | Other |
| **HIST1H2BG** | -0.3092 | 0.408 | 0.465 | 2.627e-10 | 4.688e-06 | Up in Hypoxia-low | Other |
| **LAPTM4B** | -0.4027 | 0.143 | 0.192 | 3.092e-10 | 5.517e-06 | Up in Hypoxia-low | Other |
| **FCRL3** | -0.5412 | 0.126 | 0.173 | 3.192e-10 | 5.696e-06 | Up in Hypoxia-low | Other |
| **SEPTIN1** | -0.186 | 0.638 | 0.69 | 3.955e-10 | 7.058e-06 | Up in Hypoxia-low | Other |
| **POLR2L** | -0.15 | 0.829 | 0.859 | 4.086e-10 | 7.291e-06 | Up in Hypoxia-low | Other |
| **P4HA2** | 0.3798 | 0.296 | 0.24 | 4.322e-10 | 7.712e-06 | Up in Hypoxia-high | Other |
| **VOPP1** | -0.1113 | 0.814 | 0.862 | 4.477e-10 | 7.988e-06 | Up in Hypoxia-low | Other |
| **TMBIM1** | 0.1641 | 0.776 | 0.736 | 4.664e-10 | 8.322e-06 | Up in Hypoxia-high | Other |
| **CPVL** | -0.2097 | 0.617 | 0.668 | 4.928e-10 | 8.793e-06 | Up in Hypoxia-low | Other |
| **NIBAN3** | -0.4718 | 0.156 | 0.211 | 4.929e-10 | 8.795e-06 | Up in Hypoxia-low | Other |
| **ECE1** | -0.2673 | 0.366 | 0.424 | 5.202e-10 | 9.282e-06 | Up in Hypoxia-low | Other |
| **DCHS1** | -0.412 | 0.141 | 0.182 | 5.71e-10 | 1.019e-05 | Up in Hypoxia-low | Other |
| **MS4A7** | 0.1423 | 0.65 | 0.645 | 5.918e-10 | 1.056e-05 | Up in Hypoxia-high | Other |

| | | | | | | | |
|---|---|---|---|---|---|---|---|
| **PSMB10** | 0.1128 | 0.977 | 0.97 | 6.025e-10 | 1.075e-05 | Up in Hypoxia-high | Other |
| **MS4A4A** | 0.3279 | 0.22 | 0.167 | 6.474e-10 | 1.155e-05 | Up in Hypoxia-high | Other |
| **HIST1H4E** | -0.1858 | 0.731 | 0.761 | 7.098e-10 | 1.267e-05 | Up in Hypoxia-low | Other |
| **CTNNB1** | -0.1129 | 0.907 | 0.915 | 8.036e-10 | 1.434e-05 | Up in Hypoxia-low | Other |
| **FCGR1B** | 0.1396 | 0.78 | 0.744 | 8.835e-10 | 1.576e-05 | Up in Hypoxia-high | Other |
| **GSTO1** | 0.1054 | 0.916 | 0.888 | 9.83e-10 | 1.754e-05 | Up in Hypoxia-high | Other |
| **APOBR** | 0.2146 | 0.542 | 0.485 | 1.064e-09 | 1.899e-05 | Up in Hypoxia-high | Other |
| **FGF7** | -0.4302 | 0.175 | 0.228 | 1.074e-09 | 1.916e-05 | Up in Hypoxia-low | Other |
| **LILRB3** | 0.3121 | 0.274 | 0.218 | 1.075e-09 | 1.919e-05 | Up in Hypoxia-high | Other |
| **CCDC69** | -0.2643 | 0.392 | 0.46 | 1.083e-09 | 1.933e-05 | Up in Hypoxia-low | Other |
| **CORO1C** | 0.1231 | 0.936 | 0.917 | 1.097e-09 | 1.958e-05 | Up in Hypoxia-high | Other |
| **HIST1H2BH** | -0.3273 | 0.392 | 0.454 | 1.131e-09 | 2.018e-05 | Up in Hypoxia-low | Other |
| **CELSR1** | 0.3794 | 0.204 | 0.154 | 1.135e-09 | 2.025e-05 | Up in Hypoxia-high | Other |
| **FAP** | -0.3576 | 0.227 | 0.278 | 1.159e-09 | 2.068e-05 | Up in Hypoxia-low | Other |
| **CYSTM1** | 0.2794 | 0.397 | 0.343 | 1.218e-09 | 2.173e-05 | Up in Hypoxia-high | Other |
| **CLIC1** | 0.078 | 0.992 | 0.987 | 1.228e-09 | 2.192e-05 | Up in Hypoxia-high | Other |
| **FCGR2A** | 0.1591 | 0.625 | 0.585 | 1.235e-09 | 2.203e-05 | Up in Hypoxia-high | Other |
| **USP22** | -0.1301 | 0.853 | 0.876 | 1.251e-09 | 2.233e-05 | Up in Hypoxia-low | Other |
| **HEG1** | -0.3078 | 0.226 | 0.28 | 1.274e-09 | 2.273e-05 | Up in Hypoxia-low | Other |
| **PDLIM3** | -0.2249 | 0.419 | 0.472 | 1.282e-09 | 2.287e-05 | Up in Hypoxia-low | Other |
| **CLDN5** | -0.4155 | 0.154 | 0.19 | 1.29e-09 | 2.302e-05 | Up in Hypoxia-low | Other |
| **SMAD3** | -0.3444 | 0.252 | 0.312 | 1.346e-09 | 2.401e-05 | Up in Hypoxia-low | Other |
| **RIPOR2** | -0.2995 | 0.346 | 0.404 | 1.359e-09 | 2.424e-05 | Up in Hypoxia-low | Other |
| **IFI44** | -0.3348 | 0.395 | 0.457 | 1.419e-09 | 2.532e-05 | Up in Hypoxia-low | Other |
| **KCNJ8** | -0.4783 | 0.116 | 0.162 | 1.453e-09 | 2.592e-05 | Up in Hypoxia-low | Other |
| **BIN1** | -0.2304 | 0.414 | 0.474 | 1.594e-09 | 2.844e-05 | Up in Hypoxia-low | Other |
| **CCL4L2** | 0.2861 | 0.378 | 0.323 | 1.652e-09 | 2.947e-05 | Up in Hypoxia-high | Other |
| **IGHV6-1** | -0.4705 | 0.27 | 0.33 | 1.655e-09 | 2.952e-05 | Up in Hypoxia-low | Other |
| **ARHGAP9** | -0.1863 | 0.643 | 0.689 | 1.655e-09 | 2.953e-05 | Up in Hypoxia-low | Other |
| **EID1** | -0.1228 | 0.902 | 0.916 | 1.946e-09 | 3.473e-05 | Up in Hypoxia-low | Other |
| **SVIL** | -0.3771 | 0.153 | 0.202 | 2.088e-09 | 3.726e-05 | Up in Hypoxia-low | Other |
| **CD22** | -0.6293 | 0.1 | 0.144 | 2.143e-09 | 3.824e-05 | Up in Hypoxia-low | Other |
| **FCRL1** | -0.4902 | 0.147 | 0.2 | 2.214e-09 | 3.95e-05 | Up in Hypoxia-low | Other |
| **REXO2** | -0.1753 | 0.581 | 0.634 | 2.281e-09 | 4.07e-05 | Up in Hypoxia-low | Other |
| **ATP8A1** | -0.413 | 0.171 | 0.225 | 2.284e-09 | 4.075e-05 | Up in Hypoxia-low | Other |
| **LEF1** | -0.4015 | 0.189 | 0.245 | 2.455e-09 | 4.38e-05 | Up in Hypoxia-low | Other |
| **COL5A1** | -0.166 | 0.838 | 0.854 | 2.468e-09 | 4.404e-05 | Up in Hypoxia-low | Other |
| **HDAC7** | -0.2452 | 0.428 | 0.485 | 2.529e-09 | 4.512e-05 | Up in Hypoxia-low | Other |
| **LPCAT2** | 0.2866 | 0.313 | 0.263 | 2.557e-09 | 4.563e-05 | Up in Hypoxia-high | Other |
| **HSPG2** | -0.1519 | 0.828 | 0.839 | 2.596e-09 | 4.632e-05 | Up in Hypoxia-low | Other |
| **SLC41A2** | 0.3502 | 0.202 | 0.157 | 2.66e-09 | 4.746e-05 | Up in Hypoxia-high | Other |
| **FOSL2** | 0.2225 | 0.579 | 0.529 | 2.671e-09 | 4.766e-05 | Up in Hypoxia-high | Other |
| **CCR7** | -0.3734 | 0.215 | 0.273 | 2.683e-09 | 4.787e-05 | Up in Hypoxia-low | Other |
| **SEPTIN2** | -0.117 | 0.895 | 0.909 | 2.708e-09 | 4.832e-05 | Up in Hypoxia-low | Other |

| | | | | | | | |
|---|---|---|---|---|---|---|---|
| **FKBP11** | -0.3006 | 0.278 | 0.336 | 2.819e-09 | 5.031e-05 | Up in Hypoxia-low | Other |
| **FAH** | 0.2679 | 0.342 | 0.285 | 2.93e-09 | 5.229e-05 | Up in Hypoxia-high | Other |
| **NFE2L3** | 0.408 | 0.22 | 0.168 | 3.008e-09 | 5.367e-05 | Up in Hypoxia-high | Other |
| **THBD** | -0.2445 | 0.311 | 0.343 | 3.297e-09 | 5.883e-05 | Up in Hypoxia-low | Other |
| **MCF2L** | -0.4491 | 0.12 | 0.161 | 3.447e-09 | 6.151e-05 | Up in Hypoxia-low | Other |
| **SLC39A14** | 0.241 | 0.527 | 0.485 | 3.523e-09 | 6.286e-05 | Up in Hypoxia-high | Other |
| **HIST1H4I** | -0.1683 | 0.668 | 0.72 | 3.564e-09 | 6.359e-05 | Up in Hypoxia-low | Other |
| **RAMP2** | -0.449 | 0.129 | 0.174 | 3.589e-09 | 6.405e-05 | Up in Hypoxia-low | Other |
| **SQSTM1** | 0.0428 | 1 | 1 | 3.601e-09 | 6.425e-05 | Up in Hypoxia-high | Other |
| **SPR** | 0.2992 | 0.317 | 0.26 | 3.767e-09 | 6.721e-05 | Up in Hypoxia-high | Other |
| **KLRC3** | -0.4066 | 0.136 | 0.186 | 3.774e-09 | 6.733e-05 | Up in Hypoxia-low | Other |
| **PTPRS** | -0.4465 | 0.112 | 0.153 | 3.774e-09 | 6.734e-05 | Up in Hypoxia-low | Other |
| **TCF7** | -0.3595 | 0.221 | 0.276 | 3.967e-09 | 7.078e-05 | Up in Hypoxia-low | Other |
| **ATOX1** | 0.0731 | 0.976 | 0.966 | 4.22e-09 | 7.53e-05 | Up in Hypoxia-high | Other |
| **GAL3ST4** | -0.4523 | 0.134 | 0.183 | 4.266e-09 | 7.611e-05 | Up in Hypoxia-low | Other |
| **PDGFRA** | -0.2149 | 0.453 | 0.496 | 4.648e-09 | 8.294e-05 | Up in Hypoxia-low | Other |
| **IGFBP6** | 0.4659 | 0.172 | 0.131 | 4.78e-09 | 8.53e-05 | Up in Hypoxia-high | Other |
| **APLNR** | -0.3646 | 0.178 | 0.223 | 4.928e-09 | 8.794e-05 | Up in Hypoxia-low | Other |
| **DKK 3,00** | -0.4789 | 0.117 | 0.154 | 4.953e-09 | 8.838e-05 | Up in Hypoxia-low | Other |
| **NOTCH3** | -0.1876 | 0.348 | 0.361 | 5.014e-09 | 8.947e-05 | Up in Hypoxia-low | Other |
| **SORT1** | 0.2101 | 0.377 | 0.323 | 5.036e-09 | 8.986e-05 | Up in Hypoxia-high | Other |
| **TALDO1** | 0.1044 | 0.912 | 0.879 | 5.167e-09 | 9.219e-05 | Up in Hypoxia-high | Other |
| **ISG20** | -0.1911 | 0.566 | 0.63 | 5.314e-09 | 9.482e-05 | Up in Hypoxia-low | Other |
| **SLC2A4RG** | -0.2164 | 0.468 | 0.511 | 5.34e-09 | 9.529e-05 | Up in Hypoxia-low | Other |
| **LAMA5** | -0.3552 | 0.152 | 0.197 | 6.393e-09 | 0.0001141 | Up in Hypoxia-low | Other |
| **TMED2** | -0.1356 | 0.897 | 0.923 | 6.488e-09 | 0.0001158 | Up in Hypoxia-low | Other |
| **MMP14** | 0.0746 | 0.997 | 0.994 | 6.491e-09 | 0.0001158 | Up in Hypoxia-high | Other |
| **IL33** | -0.369 | 0.158 | 0.202 | 6.622e-09 | 0.0001181 | Up in Hypoxia-low | Other |
| **OLR1** | 0.1733 | 0.587 | 0.544 | 6.699e-09 | 0.0001195 | Up in Hypoxia-high | Other |
| **CMTM6** | 0.1031 | 0.872 | 0.838 | 7.333e-09 | 0.0001308 | Up in Hypoxia-high | Other |
| **LAMB1** | -0.1595 | 0.484 | 0.535 | 7.8e-09 | 0.0001392 | Up in Hypoxia-low | Other |
| **S1PR3** | 0.2588 | 0.426 | 0.365 | 7.979e-09 | 0.0001424 | Up in Hypoxia-high | Other |
| **NUAK1** | -0.3847 | 0.122 | 0.162 | 8.516e-09 | 0.0001519 | Up in Hypoxia-low | Other |
| **PMEPA1** | -0.1663 | 0.459 | 0.481 | 8.848e-09 | 0.0001579 | Up in Hypoxia-low | Other |
| **TWF2** | 0.1903 | 0.59 | 0.558 | 8.908e-09 | 0.000159 | Up in Hypoxia-high | Other |
| **ARHGAP23** | -0.3443 | 0.174 | 0.22 | 9.009e-09 | 0.0001607 | Up in Hypoxia-low | Other |
| **HIST1H2AB** | -0.3877 | 0.219 | 0.263 | 9.014e-09 | 0.0001608 | Up in Hypoxia-low | Other |
| **GBP4** | 0.1075 | 0.892 | 0.881 | 9.099e-09 | 0.0001624 | Up in Hypoxia-high | Other |
| **PLXDC1** | -0.4056 | 0.106 | 0.148 | 9.183e-09 | 0.0001639 | Up in Hypoxia-low | Other |
| **NRROS** | 0.2546 | 0.408 | 0.359 | 9.56e-09 | 0.0001706 | Up in Hypoxia-high | Other |
| **ENAH** | -0.3899 | 0.108 | 0.147 | 1.006e-08 | 0.0001795 | Up in Hypoxia-low | Other |
| **ST6GAL1** | -0.2996 | 0.329 | 0.385 | 1.009e-08 | 0.00018 | Up in Hypoxia-low | Other |
| **NCEH1** | 0.2337 | 0.469 | 0.417 | 1.014e-08 | 0.0001809 | Up in Hypoxia-high | Other |
| **AVPI1** | 0.3472 | 0.242 | 0.19 | 1.025e-08 | 0.0001829 | Up in Hypoxia-high | Other |

| | | | | | | | |
|---|---|---|---|---|---|---|---|
| **FNBP1** | -0.1366 | 0.798 | 0.841 | 1.033e-08 | 0.0001844 | Up in Hypoxia-low | Other |
| **CNDP2** | 0.0839 | 0.992 | 0.99 | 1.087e-08 | 0.0001939 | Up in Hypoxia-high | Other |
| **RGCC** | 0.297 | 0.291 | 0.251 | 1.097e-08 | 0.0001958 | Up in Hypoxia-high | Other |
| **DNASE1L3** | -0.4802 | 0.142 | 0.193 | 1.106e-08 | 0.0001973 | Up in Hypoxia-low | Other |
| **TCL1A** | -0.6221 | 0.126 | 0.169 | 1.165e-08 | 0.0002079 | Up in Hypoxia-low | Other |
| **CNBP** | 0.119 | 0.99 | 0.986 | 1.245e-08 | 0.0002221 | Up in Hypoxia-high | Other |
| **RASGRP2** | -0.329 | 0.242 | 0.295 | 1.295e-08 | 0.000231 | Up in Hypoxia-low | Other |
| **MCM7** | -0.2444 | 0.416 | 0.473 | 1.335e-08 | 0.0002382 | Up in Hypoxia-low | Other |
| **DEPTOR** | -0.395 | 0.136 | 0.17 | 1.443e-08 | 0.0002575 | Up in Hypoxia-low | Other |
| **PCSK7** | -0.1814 | 0.549 | 0.604 | 1.522e-08 | 0.0002716 | Up in Hypoxia-low | Other |
| **RNF19B** | 0.1773 | 0.606 | 0.578 | 1.606e-08 | 0.0002866 | Up in Hypoxia-high | Other |
| **ACSS1** | -0.231 | 0.448 | 0.506 | 1.618e-08 | 0.0002888 | Up in Hypoxia-low | Other |
| **ALDH2** | 0.0778 | 0.983 | 0.972 | 1.636e-08 | 0.0002918 | Up in Hypoxia-high | Other |
| **CPE** | 0.2537 | 0.416 | 0.387 | 1.728e-08 | 0.0003083 | Up in Hypoxia-high | Other |
| **ACP2** | 0.1376 | 0.782 | 0.748 | 1.779e-08 | 0.0003175 | Up in Hypoxia-high | Other |
| **SMAD6** | -0.4103 | 0.15 | 0.185 | 1.795e-08 | 0.0003203 | Up in Hypoxia-low | Other |
| **SAMD3** | -0.4621 | 0.155 | 0.205 | 1.825e-08 | 0.0003255 | Up in Hypoxia-low | Other |
| **TNFAIP2** | 0.0879 | 0.949 | 0.929 | 1.855e-08 | 0.000331 | Up in Hypoxia-high | Other |
| **VAMP5** | 0.084 | 0.986 | 0.979 | 1.958e-08 | 0.0003494 | Up in Hypoxia-high | Other |
| **MAGED1** | -0.2575 | 0.358 | 0.415 | 1.977e-08 | 0.0003528 | Up in Hypoxia-low | Other |
| **CDC25B** | -0.2579 | 0.369 | 0.427 | 2.017e-08 | 0.0003599 | Up in Hypoxia-low | Other |
| **STXBP2** | 0.1663 | 0.71 | 0.674 | 2.087e-08 | 0.0003724 | Up in Hypoxia-high | Other |
| **SET** | -0.1293 | 0.863 | 0.887 | 2.088e-08 | 0.0003726 | Up in Hypoxia-low | Other |
| **ALPL** | -0.2628 | 0.218 | 0.262 | 2.176e-08 | 0.0003882 | Up in Hypoxia-low | Other |
| **ACVR1B** | 0.2227 | 0.416 | 0.363 | 2.193e-08 | 0.0003913 | Up in Hypoxia-high | Other |
| **SPARCL1** | -0.3136 | 0.197 | 0.243 | 2.2e-08 | 0.0003926 | Up in Hypoxia-low | Other |
| **C1RL** | 0.3276 | 0.265 | 0.215 | 2.238e-08 | 0.0003993 | Up in Hypoxia-high | Other |
| **CD72** | -0.2483 | 0.401 | 0.461 | 2.353e-08 | 0.0004198 | Up in Hypoxia-low | Other |
| **LUC7L3** | -0.1721 | 0.687 | 0.725 | 2.409e-08 | 0.0004298 | Up in Hypoxia-low | Other |
| **SEPTIN8** | -0.2809 | 0.286 | 0.336 | 2.492e-08 | 0.0004446 | Up in Hypoxia-low | Other |
| **PNISR** | -0.1302 | 0.776 | 0.815 | 2.54e-08 | 0.0004531 | Up in Hypoxia-low | Other |
| **GSDME** | 0.292 | 0.304 | 0.255 | 2.777e-08 | 0.0004955 | Up in Hypoxia-high | Other |
| **C17orf58** | 0.3711 | 0.213 | 0.165 | 2.784e-08 | 0.0004967 | Up in Hypoxia-high | Other |
| **EPHB2** | -0.3297 | 0.187 | 0.231 | 2.864e-08 | 0.0005111 | Up in Hypoxia-low | Other |
| **LTBP3** | -0.1 | 0.834 | 0.841 | 2.885e-08 | 0.0005148 | Up in Hypoxia-low | Other |
| **EGFL7** | -0.3886 | 0.132 | 0.174 | 2.924e-08 | 0.0005218 | Up in Hypoxia-low | Other |
| **VAC14** | 0.1571 | 0.552 | 0.505 | 2.931e-08 | 0.0005229 | Up in Hypoxia-high | Other |
| **ZNF358** | -0.278 | 0.316 | 0.37 | 2.963e-08 | 0.0005287 | Up in Hypoxia-low | Other |
| **ITGA6** | -0.4223 | 0.151 | 0.192 | 2.986e-08 | 0.0005327 | Up in Hypoxia-low | Other |
| **RRAS2** | -0.4564 | 0.102 | 0.144 | 3.085e-08 | 0.0005505 | Up in Hypoxia-low | Other |
| **PRKAR1B** | -0.4619 | 0.108 | 0.147 | 3.117e-08 | 0.0005561 | Up in Hypoxia-low | Other |
| **PDLIM5** | -0.1693 | 0.576 | 0.627 | 3.183e-08 | 0.000568 | Up in Hypoxia-low | Other |
| **SELPLG** | 0.1905 | 0.625 | 0.579 | 3.219e-08 | 0.0005743 | Up in Hypoxia-high | Other |
| **IFI16** | -0.1479 | 0.871 | 0.901 | 3.224e-08 | 0.0005753 | Up in Hypoxia-low | Other |

| | | | | | | | |
|---|---|---|---|---|---|---|---|
| **ATP5F1B** | 0.0928 | 0.996 | 0.995 | 3.225e-08 | 0.0005754 | Up in Hypoxia-high | Other |
| **ZBTB7A** | 0.1492 | 0.843 | 0.819 | 3.244e-08 | 0.0005788 | Up in Hypoxia-high | Other |
| **ACVRL1** | 0.1984 | 0.607 | 0.554 | 3.287e-08 | 0.0005865 | Up in Hypoxia-high | Other |
| **SLC22A15** | 0.2688 | 0.281 | 0.234 | 3.292e-08 | 0.0005874 | Up in Hypoxia-high | Other |
| **HSPA1A** | 0.1566 | 0.845 | 0.84 | 3.349e-08 | 0.0005976 | Up in Hypoxia-high | Other |
| **GSTP1** | -0.084 | 0.996 | 0.996 | 3.363e-08 | 6E-04 | Up in Hypoxia-low | Other |
| **ITGB5** | -0.237 | 0.318 | 0.354 | 3.414e-08 | 0.0006092 | Up in Hypoxia-low | Other |
| **HIST1H2AE** | -0.2228 | 0.509 | 0.56 | 3.45e-08 | 0.0006155 | Up in Hypoxia-low | Other |
| **NDRG2** | -0.2917 | 0.234 | 0.278 | 3.566e-08 | 0.0006362 | Up in Hypoxia-low | Other |
| **MAP3K7CL** | 0.172 | 0.285 | 0.258 | 4.054e-08 | 0.0007233 | Up in Hypoxia-high | Other |
| **FDX1** | 0.1335 | 0.73 | 0.679 | 4.184e-08 | 0.0007465 | Up in Hypoxia-high | Other |
| **CCL4** | 0.2676 | 0.31 | 0.279 | 4.362e-08 | 0.0007783 | Up in Hypoxia-high | Other |
| **NID1** | -0.3162 | 0.211 | 0.254 | 4.433e-08 | 0.000791 | Up in Hypoxia-low | Other |
| **TPD52** | -0.3892 | 0.131 | 0.176 | 4.496e-08 | 0.0008023 | Up in Hypoxia-low | Other |
| **PDPN** | 0.171 | 0.695 | 0.668 | 4.539e-08 | 0.0008099 | Up in Hypoxia-high | Other |
| **CD40** | 0.1253 | 0.892 | 0.874 | 4.871e-08 | 0.0008692 | Up in Hypoxia-high | Other |
| **IGF2R** | 0.2053 | 0.547 | 0.501 | 4.971e-08 | 0.000887 | Up in Hypoxia-high | Other |
| **HP1BP3** | -0.1377 | 0.808 | 0.842 | 5.091e-08 | 0.0009084 | Up in Hypoxia-low | Other |
| **ATP6AP2** | 0.1883 | 0.613 | 0.571 | 5.329e-08 | 0.0009509 | Up in Hypoxia-high | Other |
| **FARP1** | -0.2941 | 0.228 | 0.276 | 5.555e-08 | 0.0009912 | Up in Hypoxia-low | Other |
| **GPX4** | 0.0602 | 0.988 | 0.982 | 5.697e-08 | 0.001017 | Up in Hypoxia-high | Other |
| **MCRIP2** | 0.3566 | 0.222 | 0.178 | 5.87e-08 | 0.001047 | Up in Hypoxia-high | Other |
| **MALT1** | -0.3624 | 0.232 | 0.283 | 6.291e-08 | 0.001123 | Up in Hypoxia-low | Other |
| **RAB5IF** | 0.0859 | 0.948 | 0.932 | 6.351e-08 | 0.001133 | Up in Hypoxia-high | Other |
| **MARCKS** | -0.0727 | 0.975 | 0.981 | 6.372e-08 | 0.001137 | Up in Hypoxia-low | Other |
| **ATF4** | -0.2145 | 0.525 | 0.582 | 6.432e-08 | 0.001148 | Up in Hypoxia-low | Other |
| **IGHV5-10-1** | -0.6968 | 0.165 | 0.15 | 6.535e-08 | 0.001166 | Up in Hypoxia-low | Other |
| **SND1** | -0.113 | 0.884 | 0.912 | 6.578e-08 | 0.001174 | Up in Hypoxia-low | Other |
| **ACO2** | 0.1607 | 0.742 | 0.71 | 6.701e-08 | 0.001196 | Up in Hypoxia-high | Other |
| **SH3PXD2A** | -0.141 | 0.537 | 0.57 | 7.049e-08 | 0.001258 | Up in Hypoxia-low | Other |
| **C1QB** | 0.0547 | 1 | 1 | 7.279e-08 | 0.001299 | Up in Hypoxia-high | Other |
| **MAN2A1** | -0.2465 | 0.341 | 0.401 | 7.416e-08 | 0.001323 | Up in Hypoxia-low | Other |
| **KLRK1** | -0.4114 | 0.152 | 0.199 | 7.437e-08 | 0.001327 | Up in Hypoxia-low | Other |
| **EHD2** | -0.1771 | 0.45 | 0.479 | 7.654e-08 | 0.001366 | Up in Hypoxia-low | Other |
| **JAG2** | -0.3523 | 0.159 | 0.197 | 7.894e-08 | 0.001408 | Up in Hypoxia-low | Other |
| **HIST1H3G** | -0.2755 | 0.4 | 0.458 | 8.087e-08 | 0.001443 | Up in Hypoxia-low | Other |
| **CCT3** | -0.1182 | 0.855 | 0.87 | 8.212e-08 | 0.001465 | Up in Hypoxia-low | Other |
| **NUCB1** | 0.1067 | 0.971 | 0.964 | 8.216e-08 | 0.001466 | Up in Hypoxia-high | Other |
| **H2AFY** | 0.1101 | 0.802 | 0.759 | 8.351e-08 | 0.00149 | Up in Hypoxia-high | Other |
| **COL5A2** | -0.2032 | 0.595 | 0.642 | 8.36e-08 | 0.001492 | Up in Hypoxia-low | Other |
| **NIBAN1** | 0.2269 | 0.433 | 0.382 | 8.398e-08 | 0.001498 | Up in Hypoxia-high | Other |
| **ABHD2** | 0.1503 | 0.822 | 0.793 | 8.437e-08 | 0.001505 | Up in Hypoxia-high | Other |
| **ANAPC5** | -0.2639 | 0.368 | 0.424 | 8.675e-08 | 0.001548 | Up in Hypoxia-low | Other |
| **MAP1B** | -0.3253 | 0.171 | 0.205 | 9.258e-08 | 0.001652 | Up in Hypoxia-low | Other |

| | | | | | | | |
|---|---|---|---|---|---|---|---|
| **GREM1** | -0.3828 | 0.201 | 0.247 | 9.444e-08 | 0.001685 | Up in Hypoxia-low | Other |
| **NET1** | -0.3692 | 0.152 | 0.196 | 9.616e-08 | 0.001716 | Up in Hypoxia-low | Other |
| **RNASEK** | 0.0878 | 0.896 | 0.864 | 9.85e-08 | 0.001758 | Up in Hypoxia-high | Other |
| **TRA2A** | -0.1146 | 0.786 | 0.828 | 1.06e-07 | 0.001891 | Up in Hypoxia-low | Other |
| **TOP2B** | -0.1449 | 0.594 | 0.653 | 1.1e-07 | 0.001962 | Up in Hypoxia-low | Other |
| **HIST1H2BN** | -0.1682 | 0.635 | 0.683 | 1.185e-07 | 0.002114 | Up in Hypoxia-low | Other |
| **TMEM150B** | 0.2521 | 0.339 | 0.29 | 1.218e-07 | 0.002174 | Up in Hypoxia-high | Other |
| **RBM47** | 0.1871 | 0.552 | 0.498 | 1.233e-07 | 0.0022 | Up in Hypoxia-high | Other |
| **HIST1H2AD** | -0.2077 | 0.567 | 0.603 | 1.246e-07 | 0.002223 | Up in Hypoxia-low | Other |
| **TMBIM6** | 0.0911 | 0.998 | 0.992 | 1.283e-07 | 0.002289 | Up in Hypoxia-high | Other |
| **GADD45G** | 0.2334 | 0.411 | 0.368 | 1.296e-07 | 0.002313 | Up in Hypoxia-high | Other |
| **MFAP2** | -0.3374 | 0.161 | 0.203 | 1.333e-07 | 0.002379 | Up in Hypoxia-low | Other |
| **B4GALT2** | -0.4631 | 0.104 | 0.14 | 1.395e-07 | 0.002489 | Up in Hypoxia-low | Other |
| **FASTK** | -0.2128 | 0.411 | 0.465 | 1.404e-07 | 0.002505 | Up in Hypoxia-low | Other |
| **CYTH4** | 0.1529 | 0.566 | 0.517 | 1.477e-07 | 0.002636 | Up in Hypoxia-high | Other |
| **TMEM173** | -0.1959 | 0.501 | 0.551 | 1.491e-07 | 0.00266 | Up in Hypoxia-low | Other |
| **RBM12** | -0.2393 | 0.392 | 0.452 | 1.537e-07 | 0.002743 | Up in Hypoxia-low | Other |
| **COL9A2** | 0.3854 | 0.186 | 0.145 | 1.646e-07 | 0.002937 | Up in Hypoxia-high | Other |
| **MTFP1** | 0.3487 | 0.227 | 0.181 | 1.693e-07 | 0.003021 | Up in Hypoxia-high | Other |
| **CLDN23** | 0.3789 | 0.182 | 0.141 | 1.704e-07 | 0.00304 | Up in Hypoxia-high | Other |
| **TNS2** | -0.2769 | 0.212 | 0.259 | 1.733e-07 | 0.003092 | Up in Hypoxia-low | Other |
| **IGHV1-46** | -0.7648 | 0.226 | 0.255 | 1.754e-07 | 0.00313 | Up in Hypoxia-low | Other |
| **EPB41L3** | 0.1631 | 0.657 | 0.621 | 1.767e-07 | 0.003153 | Up in Hypoxia-high | Other |
| **PFKFB4** | 0.4262 | 0.141 | 0.105 | 1.817e-07 | 0.003242 | Up in Hypoxia-high | Other |
| **LGMN** | -0.1642 | 0.813 | 0.831 | 1.896e-07 | 0.003383 | Up in Hypoxia-low | Other |
| **SNTB1** | 0.1881 | 0.382 | 0.333 | 1.917e-07 | 0.00342 | Up in Hypoxia-high | Other |
| **AGTRAP** | 0.1845 | 0.536 | 0.484 | 1.928e-07 | 0.003441 | Up in Hypoxia-high | Other |
| **TNFRSF14** | 0.1295 | 0.861 | 0.827 | 1.935e-07 | 0.003453 | Up in Hypoxia-high | Other |
| **TDO2** | -0.3887 | 0.116 | 0.156 | 1.937e-07 | 0.003456 | Up in Hypoxia-low | Other |
| **GCH1** | 0.1075 | 0.65 | 0.616 | 1.95e-07 | 0.003479 | Up in Hypoxia-high | Other |
| **GRINA** | 0.0714 | 0.995 | 0.992 | 1.955e-07 | 0.003487 | Up in Hypoxia-high | Other |
| **IRF4** | -0.3065 | 0.265 | 0.315 | 2.014e-07 | 0.003593 | Up in Hypoxia-low | Other |
| **TRIOBP** | -0.1694 | 0.522 | 0.568 | 2.07e-07 | 0.003694 | Up in Hypoxia-low | Other |
| **NPDC1** | -0.2379 | 0.356 | 0.397 | 2.089e-07 | 0.003728 | Up in Hypoxia-low | Other |
| **SHE** | -0.4049 | 0.11 | 0.147 | 2.111e-07 | 0.003767 | Up in Hypoxia-low | Other |
| **IGFBP5** | -0.2507 | 0.349 | 0.389 | 2.125e-07 | 0.003792 | Up in Hypoxia-low | Other |
| **MSR1** | 0.2485 | 0.404 | 0.35 | 2.18e-07 | 0.00389 | Up in Hypoxia-high | Other |
| **EGR1** | -0.3215 | 0.309 | 0.364 | 2.198e-07 | 0.003922 | Up in Hypoxia-low | Other |
| **PSAP** | 0.0395 | 1 | 1 | 2.22e-07 | 0.003961 | Up in Hypoxia-high | Other |
| **MN1** | -0.327 | 0.137 | 0.157 | 2.224e-07 | 0.003968 | Up in Hypoxia-low | Other |
| **CHIT1** | -0.0917 | 0.516 | 0.484 | 2.313e-07 | 0.004128 | Up in Hypoxia-low | Other |
| **ZEB2** | 0.0991 | 0.845 | 0.828 | 2.439e-07 | 0.004352 | Up in Hypoxia-high | Other |
| **VAV1** | 0.1069 | 0.802 | 0.795 | 2.514e-07 | 0.004486 | Up in Hypoxia-high | Other |
| **H3F3B** | 0.1407 | 0.888 | 0.877 | 2.563e-07 | 0.004574 | Up in Hypoxia-high | Other |

| | | | | | | | |
|---|---|---|---|---|---|---|---|
| **XCR1** | -0.4194 | 0.137 | 0.18 | 2.605e-07 | 0.004648 | Up in Hypoxia-low | Other |
| **GATA6** | -0.4146 | 0.122 | 0.158 | 2.627e-07 | 0.004687 | Up in Hypoxia-low | Other |
| **PTPN6** | 0.1348 | 0.858 | 0.83 | 2.641e-07 | 0.004713 | Up in Hypoxia-high | Other |
| **JUNB** | 0.0879 | 0.945 | 0.941 | 2.719e-07 | 0.004851 | Up in Hypoxia-high | Other |
| **FKBP9** | -0.2484 | 0.33 | 0.379 | 2.723e-07 | 0.004858 | Up in Hypoxia-low | Other |
| **TIFAB** | 0.2048 | 0.362 | 0.316 | 2.747e-07 | 0.004901 | Up in Hypoxia-high | Other |
| **KMO** | 0.2684 | 0.198 | 0.165 | 2.838e-07 | 0.005064 | Up in Hypoxia-high | Other |
| **CD34** | -0.3847 | 0.141 | 0.181 | 2.882e-07 | 0.005143 | Up in Hypoxia-low | Other |
| **RBM6** | -0.2122 | 0.436 | 0.495 | 2.938e-07 | 0.005242 | Up in Hypoxia-low | Other |
| **SHISA5** | -0.1743 | 0.694 | 0.73 | 3.002e-07 | 0.005357 | Up in Hypoxia-low | Other |
| **COX8A** | 0.0966 | 0.967 | 0.957 | 3.035e-07 | 0.005415 | Up in Hypoxia-high | Other |
| **APOL3** | 0.0882 | 0.94 | 0.924 | 3.039e-07 | 0.005422 | Up in Hypoxia-high | Other |
| **PLK3** | 0.1964 | 0.48 | 0.431 | 3.094e-07 | 0.005521 | Up in Hypoxia-high | Other |
| **PTAFR** | 0.1368 | 0.704 | 0.657 | 3.108e-07 | 0.005546 | Up in Hypoxia-high | Other |
| **HIST1H2BI** | -0.3363 | 0.272 | 0.323 | 3.227e-07 | 0.005759 | Up in Hypoxia-low | Other |
| **SLC3A2** | 0.1206 | 0.877 | 0.855 | 3.232e-07 | 0.005766 | Up in Hypoxia-high | Other |
| **RASA2** | -0.1552 | 0.617 | 0.665 | 3.282e-07 | 0.005857 | Up in Hypoxia-low | Other |
| **ATP6V1A** | 0.1184 | 0.876 | 0.849 | 3.412e-07 | 0.006087 | Up in Hypoxia-high | Other |
| **P2RX7** | 0.1785 | 0.445 | 0.41 | 3.432e-07 | 0.006123 | Up in Hypoxia-high | Other |
| **STX4** | 0.1402 | 0.631 | 0.587 | 3.565e-07 | 0.006362 | Up in Hypoxia-high | Other |
| **SCARA5** | 0.3797 | 0.206 | 0.161 | 3.589e-07 | 0.006404 | Up in Hypoxia-high | Other |
| **SRSF11** | -0.1185 | 0.816 | 0.846 | 3.638e-07 | 0.006491 | Up in Hypoxia-low | Other |
| **TMEM51** | 0.2783 | 0.328 | 0.278 | 3.777e-07 | 0.006739 | Up in Hypoxia-high | Other |
| **OAS2** | -0.2138 | 0.682 | 0.728 | 3.791e-07 | 0.006764 | Up in Hypoxia-low | Other |
| **GLIPR2** | 0.1436 | 0.628 | 0.589 | 3.82e-07 | 0.006816 | Up in Hypoxia-high | Other |
| **SLC25A24** | 0.2673 | 0.327 | 0.286 | 4.21e-07 | 0.007512 | Up in Hypoxia-high | Other |
| **CYB5A** | 0.152 | 0.649 | 0.601 | 4.255e-07 | 0.007592 | Up in Hypoxia-high | Other |
| **GPR84** | 0.2468 | 0.28 | 0.246 | 4.368e-07 | 0.007794 | Up in Hypoxia-high | Other |
| **PA2G4** | -0.0929 | 0.904 | 0.905 | 4.406e-07 | 0.007862 | Up in Hypoxia-low | Other |
| **ADAMTS1** | -0.3598 | 0.192 | 0.234 | 4.496e-07 | 0.008022 | Up in Hypoxia-low | Other |
| **RIPK2** | 0.2636 | 0.205 | 0.166 | 4.512e-07 | 0.00805 | Up in Hypoxia-high | Other |
| **HIST1H2BO** | -0.258 | 0.381 | 0.437 | 4.528e-07 | 0.00808 | Up in Hypoxia-low | Other |
| **CYB561A3** | -0.1717 | 0.596 | 0.643 | 4.61e-07 | 0.008226 | Up in Hypoxia-low | Other |
| **NECTIN4** | 0.3222 | 0.169 | 0.131 | 4.665e-07 | 0.008323 | Up in Hypoxia-high | Other |
| **BCL11A** | -0.4121 | 0.148 | 0.193 | 4.679e-07 | 0.008349 | Up in Hypoxia-low | Other |
| **EMP3** | 0.1228 | 0.865 | 0.845 | 4.75e-07 | 0.008475 | Up in Hypoxia-high | Other |
| **ATF3** | 0.2681 | 0.287 | 0.249 | 5.152e-07 | 0.009193 | Up in Hypoxia-high | Other |
| **NDUFB5** | 0.2788 | 0.21 | 0.168 | 5.199e-07 | 0.009276 | Up in Hypoxia-high | Other |
| **MEF2B** | -0.3574 | 0.179 | 0.218 | 5.566e-07 | 0.009931 | Up in Hypoxia-low | Other |
| **HCK** | 0.1882 | 0.465 | 0.417 | 5.579e-07 | 0.009954 | Up in Hypoxia-high | Other |
| **RNF144B** | 0.148 | 0.566 | 0.533 | 5.611e-07 | 0.01001 | Up in Hypoxia-high | Other |
| **PGGHG** | -0.1933 | 0.449 | 0.501 | 5.616e-07 | 0.01002 | Up in Hypoxia-low | Other |
| **HIST1H3I** | -0.1899 | 0.544 | 0.579 | 5.668e-07 | 0.01011 | Up in Hypoxia-low | Other |
| **CSGALNAC** | -0.2922 | 0.171 | 0.214 | 5.774e-07 | 0.0103 | Up in Hypoxia-low | Other |

| | | | | | | | |
|---|---|---|---|---|---|---|---|
| **SORL1** | -0.1817 | 0.423 | 0.475 | 5.878e-07 | 0.01049 | Up in Hypoxia-low | Other |
| **ADM** | 0.3575 | 0.157 | 0.126 | 6.044e-07 | 0.01078 | Up in Hypoxia-high | Other |
| **KLF16** | 0.3137 | 0.235 | 0.189 | 6.402e-07 | 0.01142 | Up in Hypoxia-high | Other |
| **TMEM37** | -0.3545 | 0.125 | 0.161 | 6.471e-07 | 0.01155 | Up in Hypoxia-low | Other |
| **PCM1** | -0.2293 | 0.398 | 0.457 | 6.51e-07 | 0.01162 | Up in Hypoxia-low | Other |
| **GLA** | 0.2294 | 0.357 | 0.31 | 6.711e-07 | 0.01197 | Up in Hypoxia-high | Other |
| **ZNF428** | -0.2067 | 0.375 | 0.419 | 6.813e-07 | 0.01216 | Up in Hypoxia-low | Other |
| **MUC1** | 0.3647 | 0.164 | 0.128 | 6.961e-07 | 0.01242 | Up in Hypoxia-high | Other |
| **SRM** | -0.2309 | 0.354 | 0.407 | 7.031e-07 | 0.01255 | Up in Hypoxia-low | Other |
| **STX7** | -0.1755 | 0.586 | 0.636 | 7.094e-07 | 0.01266 | Up in Hypoxia-low | Other |
| **RAB20** | 0.2004 | 0.336 | 0.304 | 7.177e-07 | 0.01281 | Up in Hypoxia-high | Other |
| **RABGAP1L** | -0.2422 | 0.369 | 0.426 | 7.223e-07 | 0.01289 | Up in Hypoxia-low | Other |
| **ARHGEF17** | -0.3463 | 0.155 | 0.187 | 7.375e-07 | 0.01316 | Up in Hypoxia-low | Other |
| **MFSD12** | 0.1378 | 0.792 | 0.76 | 7.465e-07 | 0.01332 | Up in Hypoxia-high | Other |
| **DNPH1** | -0.1426 | 0.623 | 0.653 | 7.71e-07 | 0.01376 | Up in Hypoxia-low | Other |
| **DUT** | -0.1585 | 0.598 | 0.649 | 7.724e-07 | 0.01378 | Up in Hypoxia-low | Other |
| **PBX1** | -0.3688 | 0.146 | 0.184 | 7.73e-07 | 0.01379 | Up in Hypoxia-low | Other |
| **NKG7** | 0.1164 | 0.626 | 0.634 | 7.879e-07 | 0.01406 | Up in Hypoxia-high | Other |
| **ARPC5** | 0.1156 | 0.965 | 0.953 | 7.959e-07 | 0.0142 | Up in Hypoxia-high | Other |
| **DARS** | 0.2944 | 0.202 | 0.159 | 7.972e-07 | 0.01422 | Up in Hypoxia-high | Other |
| **IL3RA** | 0.1904 | 0.427 | 0.385 | 7.986e-07 | 0.01425 | Up in Hypoxia-high | Other |
| **WDR82** | -0.1318 | 0.762 | 0.799 | 8.259e-07 | 0.01474 | Up in Hypoxia-low | Other |
| **SRSF3** | -0.0844 | 0.97 | 0.975 | 8.346e-07 | 0.01489 | Up in Hypoxia-low | Other |
| **LIMA1** | -0.3392 | 0.161 | 0.202 | 8.493e-07 | 0.01515 | Up in Hypoxia-low | Other |
| **HLA-E** | 0.0563 | 1 | 1 | 8.69e-07 | 0.01551 | Up in Hypoxia-high | Other |
| **NCKAP1** | -0.2354 | 0.302 | 0.34 | 8.853e-07 | 0.0158 | Up in Hypoxia-low | Other |
| **PDLIM7** | -0.1365 | 0.636 | 0.656 | 8.867e-07 | 0.01582 | Up in Hypoxia-low | Other |
| **ARHGAP45** | -0.0998 | 0.83 | 0.861 | 9.02e-07 | 0.01609 | Up in Hypoxia-low | Other |
| **ARHGEF1** | -0.1161 | 0.882 | 0.899 | 9.395e-07 | 0.01676 | Up in Hypoxia-low | Other |
| **ANXA11** | 0.0874 | 0.967 | 0.957 | 9.677e-07 | 0.01727 | Up in Hypoxia-high | Other |
| **HDLBP** | -0.1041 | 0.916 | 0.935 | 9.961e-07 | 0.01777 | Up in Hypoxia-low | Other |
| **IL10RA** | 0.0824 | 0.859 | 0.831 | 1.015e-06 | 0.01811 | Up in Hypoxia-high | Other |
| **IFI44L** | -0.3908 | 0.325 | 0.363 | 1.016e-06 | 0.01813 | Up in Hypoxia-low | Other |
| **PPM1G** | -0.1405 | 0.629 | 0.673 | 1.048e-06 | 0.0187 | Up in Hypoxia-low | Other |
| **CIAO2B** | 0.17 | 0.612 | 0.573 | 1.049e-06 | 0.01872 | Up in Hypoxia-high | Other |
| **NME4** | -0.2937 | 0.272 | 0.32 | 1.05e-06 | 0.01874 | Up in Hypoxia-low | Other |
| **NAP1L4** | -0.1414 | 0.582 | 0.633 | 1.069e-06 | 0.01908 | Up in Hypoxia-low | Other |
| **MAP1S** | 0.2799 | 0.316 | 0.265 | 1.072e-06 | 0.01913 | Up in Hypoxia-high | Other |
| **HNRNPUL1** | -0.0727 | 0.994 | 0.992 | 1.079e-06 | 0.01926 | Up in Hypoxia-low | Other |
| **SLC2A6** | 0.2362 | 0.44 | 0.391 | 1.117e-06 | 0.01992 | Up in Hypoxia-high | Other |
| **PLCG2** | -0.2988 | 0.264 | 0.314 | 1.183e-06 | 0.02112 | Up in Hypoxia-low | Other |
| **PPIB** | -0.059 | 0.986 | 0.987 | 1.238e-06 | 0.0221 | Up in Hypoxia-low | Other |
| **GCN1** | -0.1944 | 0.428 | 0.483 | 1.252e-06 | 0.02234 | Up in Hypoxia-low | Other |
| **PRKD2** | -0.2051 | 0.383 | 0.43 | 1.271e-06 | 0.02268 | Up in Hypoxia-low | Other |

| | | | | | | | |
|---|---|---|---|---|---|---|---|
| **BCL6** | 0.1114 | 0.712 | 0.68 | 1.282e-06 | 0.02287 | Up in Hypoxia-high | Other |
| **M6PR** | 0.0874 | 0.94 | 0.924 | 1.318e-06 | 0.02351 | Up in Hypoxia-high | Other |
| **SRC** | 0.1676 | 0.621 | 0.582 | 1.355e-06 | 0.02418 | Up in Hypoxia-high | Other |
| **RNF126** | -0.1838 | 0.406 | 0.452 | 1.409e-06 | 0.02513 | Up in Hypoxia-low | Other |
| **PCMTD2** | -0.2373 | 0.298 | 0.345 | 1.428e-06 | 0.02548 | Up in Hypoxia-low | Other |
| **TPST1** | -0.3 | 0.144 | 0.183 | 1.472e-06 | 0.02626 | Up in Hypoxia-low | Other |
| **ACOT11** | 0.4029 | 0.135 | 0.101 | 1.481e-06 | 0.02642 | Up in Hypoxia-high | Other |
| **CLEC4E** | 0.2936 | 0.195 | 0.157 | 1.498e-06 | 0.02673 | Up in Hypoxia-high | Other |
| **FKBP10** | -0.1526 | 0.496 | 0.533 | 1.514e-06 | 0.02702 | Up in Hypoxia-low | Other |
| **CNIH4** | 0.3627 | 0.165 | 0.125 | 1.516e-06 | 0.02705 | Up in Hypoxia-high | Other |
| **CARHSP1** | -0.364 | 0.174 | 0.209 | 1.596e-06 | 0.02848 | Up in Hypoxia-low | Other |
| **VAMP8** | 0.0745 | 0.965 | 0.951 | 1.6e-06 | 0.02855 | Up in Hypoxia-high | Other |
| **ZNF366** | -0.3839 | 0.136 | 0.171 | 1.618e-06 | 0.02887 | Up in Hypoxia-low | Other |
| **CYC1** | 0.1474 | 0.783 | 0.751 | 1.683e-06 | 0.03003 | Up in Hypoxia-high | Other |
| **GSDMB** | -0.2634 | 0.256 | 0.306 | 1.73e-06 | 0.03086 | Up in Hypoxia-low | Other |
| **LAMB2** | -0.1599 | 0.502 | 0.547 | 1.731e-06 | 0.03089 | Up in Hypoxia-low | Other |
| **STEAP3** | 0.176 | 0.574 | 0.526 | 1.742e-06 | 0.03108 | Up in Hypoxia-high | Other |
| **RBFOX2** | -0.2707 | 0.167 | 0.206 | 1.783e-06 | 0.03182 | Up in Hypoxia-low | Other |
| **PPP2R5C** | -0.1547 | 0.701 | 0.739 | 1.817e-06 | 0.03241 | Up in Hypoxia-low | Other |
| **ZRANB2** | -0.2053 | 0.443 | 0.498 | 1.818e-06 | 0.03244 | Up in Hypoxia-low | Other |
| **MYC** | 0.133 | 0.804 | 0.766 | 1.819e-06 | 0.03246 | Up in Hypoxia-high | Other |
| **ORM1** | 0.3421 | 0.119 | 0.105 | 1.834e-06 | 0.03273 | Up in Hypoxia-high | Other |
| **TRIO** | -0.2591 | 0.167 | 0.208 | 1.846e-06 | 0.03294 | Up in Hypoxia-low | Other |
| **GRB2** | 0.1172 | 0.761 | 0.731 | 1.882e-06 | 0.03358 | Up in Hypoxia-high | Other |
| **NKD2** | -0.2653 | 0.131 | 0.163 | 1.883e-06 | 0.0336 | Up in Hypoxia-low | Other |
| **TCIRG1** | 0.098 | 0.971 | 0.964 | 1.913e-06 | 0.03413 | Up in Hypoxia-high | Other |
| **PHLDB2** | -0.3421 | 0.144 | 0.184 | 1.954e-06 | 0.03487 | Up in Hypoxia-low | Other |
| **COX5A** | 0.094 | 0.983 | 0.976 | 1.978e-06 | 0.0353 | Up in Hypoxia-high | Other |
| **S100A16** | -0.241 | 0.24 | 0.275 | 1.998e-06 | 0.03564 | Up in Hypoxia-low | Other |
| **NAIP** | -0.1861 | 0.518 | 0.568 | 2.002e-06 | 0.03573 | Up in Hypoxia-low | Other |
| **MMP19** | 0.3345 | 0.199 | 0.159 | 2.013e-06 | 0.03593 | Up in Hypoxia-high | Other |
| **PCSK6** | 0.3888 | 0.187 | 0.152 | 2.029e-06 | 0.03621 | Up in Hypoxia-high | Other |
| **CD19** | -0.3912 | 0.138 | 0.177 | 2.033e-06 | 0.03627 | Up in Hypoxia-low | Other |
| **ANPEP** | 0.1981 | 0.362 | 0.322 | 2.041e-06 | 0.03642 | Up in Hypoxia-high | Other |
| **ARHGAP4** | -0.1581 | 0.529 | 0.575 | 2.055e-06 | 0.03667 | Up in Hypoxia-low | Other |
| **FBXW4** | -0.2359 | 0.341 | 0.389 | 2.128e-06 | 0.03797 | Up in Hypoxia-low | Other |
| **CREG1** | 0.0583 | 0.888 | 0.865 | 2.164e-06 | 0.03861 | Up in Hypoxia-high | Other |
| **RILP** | 0.2012 | 0.364 | 0.315 | 2.17e-06 | 0.03872 | Up in Hypoxia-high | Other |
| **ACO1** | 0.2086 | 0.477 | 0.426 | 2.181e-06 | 0.03892 | Up in Hypoxia-high | Other |
| **USP34** | -0.2279 | 0.383 | 0.439 | 2.346e-06 | 0.04185 | Up in Hypoxia-low | Other |
| **TBC1D4** | -0.3211 | 0.174 | 0.22 | 2.359e-06 | 0.04209 | Up in Hypoxia-low | Other |
| **TCF7L2** | 0.1758 | 0.346 | 0.301 | 2.427e-06 | 0.0433 | Up in Hypoxia-high | Other |
| **DIDO1** | -0.1545 | 0.601 | 0.638 | 2.507e-06 | 0.04474 | Up in Hypoxia-low | Other |
| **CPM** | 0.1224 | 0.611 | 0.555 | 2.511e-06 | 0.04481 | Up in Hypoxia-high | Other |

| | | | | | | | |
|---|---|---|---|---|---|---|---|
| **SPCS3** | -0.1374 | 0.714 | 0.756 | 2.522e-06 | 0.04499 | Up in Hypoxia-low | Other |
| **MAN2B1** | 0.0763 | 0.944 | 0.929 | 2.545e-06 | 0.04542 | Up in Hypoxia-high | Other |
| **KLRG1** | -0.4059 | 0.13 | 0.171 | 2.575e-06 | 0.04595 | Up in Hypoxia-low | Other |
| **RUBCNL** | -0.3751 | 0.186 | 0.232 | 2.634e-06 | 0.047 | Up in Hypoxia-low | Other |
| **MYBL2** | -0.4836 | 0.109 | 0.147 | 2.638e-06 | 0.04706 | Up in Hypoxia-low | Other |
| **WBP2** | 0.1327 | 0.663 | 0.635 | 2.669e-06 | 0.04762 | Up in Hypoxia-high | Other |
| **ALDH1B1** | -0.2513 | 0.169 | 0.206 | 2.795e-06 | 0.04987 | Up in Hypoxia-low | Other |
| **TEAD4** | -0.4095 | 0.105 | 0.139 | 2.799e-06 | 0.04994 | Up in Hypoxia-low | Other |
| **PILRB** | -0.208 | 0.462 | 0.516 | 2.799e-06 | 0.04994 | Up in Hypoxia-low | Other |

**Supplementary Table 5.** Distribution of SPP1-high and SPP1-low spots by hypoxia status across all 14,226 spatial transcriptomic spots.

| Hypoxia status | SPP1-high spots | SPP1-low spots | Row total |
|---|---|---|---|
| Hypoxia-high | **1,228 (36.2%)** | 2,166 (63.8%) | 3,394 (100%) |
| Hypoxia-low | 1,334 (12.3%) | 9,498 (87.7%) | 10,832 (100%) |

Chi-squared $\chi^2 = 995.26$, df = 1, $p < 2.2 \times 10^{-16}$ (Pearson chi-squared test with Yates' continuity correction). Odds ratio = 4.04 (95% CI: 3.69–4.42, Fisher's exact test mid-p method, $p = 2.23 \times 10^{-195}$).

SPP1-high: spots with SCT-normalised SPP1 expression exceeding both the absolute floor of 1.099 normalised units (global median of expressing spots) and the within-granuloma 75th percentile.

Hypoxia-high: spots with AddModuleScore exceeding both zero (background level) and the within-granuloma 75th percentile. Values shown as n (row %).

Supplementary Table 6

| Sample_Granuloma_ID | Patient | Total_spots | Dominant_cluster | Pct_cluster3 | N_hypoxia_high | Pct_hypoxia_high | Median_hypoxia | N_spp1_high | Pct_spp1_high | Median_SPP1_expr | Pct_both_high | Spearman_r | Cor_padj | N_spots_corr | Hypoxia_category | Cor_group |
|---|---|---|---|---|---|---|---|---|---|---|---|---|---|---|---|---|
| **Patient_1Granuloma_A** | Patient_1 | 1513 | 1 | 12.1 | 368 | 24.3 | -0.0704 | 107 | 7.1 | 0 | 1.7 | 0.0688 | 0.01603 | 1513 | Sufficient for analysis | Weak positive, significant |
| **Patient_1Granuloma_B** | Patient_1 | 63 | 5 | 0 | 5 | 7.9 | -0.1743 | 3 | 4.8 | 0 | 0 | -0.3472 | 0.01225 | 63 | Sufficient for analysis | Negative, significant |
| **Patient_1Granuloma_C** | Patient_1 | 1047 | 1 | 10.4 | 188 | 18 | -0.0897 | 78 | 7.4 | 0 | 1.4 | 0.0731 | 0.03603 | 1047 | Sufficient for analysis | Weak positive, significant |
| **Patient_2_1Granuloma_A** | Patient_2 | 762 | 2 | 6.2 | 191 | 25.1 | -0.0527 | 128 | 16.8 | 0.6931 | 6.2 | 0.2132 | 1.186e-08 | 762 | Sufficient for analysis | Weak positive, significant |
| **Patient_2_1Granuloma_B** | Patient_2 | 322 | 2 | 5.6 | 61 | 18.9 | -0.093 | 18 | 5.6 | 0.6931 | 0.9 | 0.0417 | 0.5696 | 322 | Sufficient for analysis | Weak positive, NS |
| **Patient_2_1Granuloma_C** | Patient_2 | 82 | 4 | 0 | 17 | 20.7 | -0.0985 | 6 | 7.3 | 0.6931 | 1.2 | 0.2346 | 0.0616 | 82 | Sufficient for analysis | Weak positive, NS |
| **Patient_2_1Granuloma_D** | Patient_2 | 20 | 5 | 0 | 1 | 5 | -0.1206 | 0 | 0 | 0 | 0 | 0 | 1 | 20 | Insufficient hypoxia-high spots | Negative, NS |
| **Patient_2_1Granuloma_E** | Patient_2 | 14 | 5 | 0 | 1 | 7.1 | -0.1318 | 0 | 0 | 0 | 0 | 0.3239 | 0.3785 | 14 | Insufficient hypoxia-high spots | Strong positive, underpowered |
| **Patient_2_1Granuloma_F** | Patient_2 | 729 | 7 | 6.9 | 182 | 25 | -0.0317 | 154 | 21.1 | 0.6931 | 7.7 | 0.2498 | 4.664e-11 | 729 | Sufficient for analysis | Weak positive, significant |
| **Patient_2_1Granuloma_G** | Patient_2 | 130 | 2 | 0 | 33 | 25.4 | -0.0458 | 22 | 16.9 | 1.0986 | 5.4 | 0.0627 | 0.5858 | 130 | Sufficient for analysis | Weak positive, NS |
| **Patient_2_2Granuloma_A** | Patient_2 | 858 | 5 | 13.2 | 215 | 25.1 | -0.0551 | 148 | 17.2 | 0.6931 | 9.4 | 0.3164 | 1.574e-20 | 858 | Sufficient for analysis | Strong positive, significant |
| **Patient_2_2Granuloma_B** | Patient_2 | 418 | 1 | 6.2 | 86 | 20.6 | -0.0844 | 13 | 3.1 | 0 | 1.7 | 0.1581 | 0.003093 | 418 | Sufficient for analysis | Weak positive, significant |
| **Patient_2_2Granuloma_C** | Patient_2 | 121 | 4 | 1.7 | 18 | 14.9 | -0.0952 | 11 | 9.1 | 0.6931 | 1.7 | 0.0952 | 0.4269 | 121 | Sufficient for analysis | Weak positive, NS |
| **Patient_2_2Granuloma_D** | Patient_2 | 15 | 4 | 0 | 3 | 20 | -0.1028 | 2 | 13.3 | 0.6931 | 6.7 | 0.0723 | 0.8865 | 15 | Insufficient hypoxia-high spots | Weak positive, NS |
| **Patient_2_2Granuloma_E** | Patient_2 | 707 | 8 | 8.5 | 177 | 25 | -0.0301 | 172 | 24.3 | 0.6931 | 8.6 | 0.2506 | 7.49e-11 | 707 | Sufficient for analysis | Weak positive, significant |
| **Patient_3_1Granuloma_0** | Patient_3 | 11 | 1 | 9.1 | 3 | 27.3 | -0.0759 | 3 | 27.3 | 1.7918 | 27.3 | 0.656 | 0.05321 | 11 | Insufficient hypoxia-high spots | Strong positive, underpowered |
| **Patient_3_1Granuloma_A** | Patient_3 | 54 | 3 | 33.3 | 14 | 25.9 | 0.1241 | 14 | 25.9 | 2.3502 | 14.8 | 0.7341 | 1.335e-09 | 54 | Sufficient for analysis | Strong positive, significant |
| **Patient_3_1Granuloma_B** | Patient_3 | 45 | 4 | 13.3 | 11 | 24.4 | -0.0507 | 11 | 24.4 | 1.6094 | 13.3 | 0.4321 | 0.007288 | 45 | Sufficient for analysis | Strong positive, significant |
| **Patient_3_1Granuloma_C** | Patient_3 | 12 | 4 | 0 | 3 | 25 | 7E-04 | 3 | 25 | 1.7006 | 8.3 | 0.3746 | 0.3533 | 12 | Insufficient hypoxia-high spots | Strong positive, underpowered |
| **Patient_3_1Granuloma_D** | Patient_3 | 41 | 4 | 19.5 | 10 | 24.4 | 0.0112 | 9 | 22 | 1.7918 | 7.3 | 0.25 | 0.197 | 41 | Sufficient for analysis | Weak positive, NS |
| **Patient_3_1Granuloma_E** | Patient_3 | 299 | 3 | 30.1 | 75 | 25.1 | 0.0611 | 75 | 25.1 | 2.3026 | 14 | 0.5612 | 2.852e-25 | 299 | Sufficient for analysis | Strong positive, significant |
| **Patient_3_1Granuloma_F** | Patient_3 | 15 | 2 | 0 | 4 | 26.7 | 0.0278 | 4 | 26.7 | 2.4849 | 6.7 | 0.0215 | 0.9718 | 15 | Insufficient hypoxia-high spots | Weak positive, NS |
| **Patient_3_1Granuloma_H** | Patient_3 | 271 | 1 | 32.1 | 68 | 25.1 | 0.0634 | 67 | 24.7 | 2.7081 | 18.1 | 0.7394 | 4.79e-47 | 271 | Sufficient for analysis | Strong positive, significant |
| **Patient_3_1Granuloma_I** | Patient_3 | 24 | 1 | 4.2 | 6 | 25 | 0.0684 | 5 | 20.8 | 2.5249 | 12.5 | 0.5902 | 0.006 | 24 | Sufficient for analysis | Strong positive, significant |
| **Patient_3_1Granuloma_J** | Patient_3 | 53 | 1 | 13.2 | 13 | 24.5 | 0.0378 | 12 | 22.6 | 2.7081 | 15.1 | 0.7863 | 2.022e-11 | 53 | Sufficient for analysis | Strong positive, significant |
| **Patient_3_1Granuloma_L** | Patient_3 | 562 | 1 | 31.7 | 141 | 25.1 | 0.071 | 140 | 24.9 | 2.6391 | 18.9 | 0.7911 | 7.713e-120 | 562 | Sufficient for analysis | Strong positive, significant |
| **Patient_3_1Granuloma_M** | Patient_3 | 15 | 5 | 0 | 0 | 0 | -0.1742 | 2 | 13.3 | 1.3863 | 0 | -0.2178 | 0.5559 | 15 | Hypoxia-negative | Negative, NS |
| **Patient_3_1Granuloma_P** | Patient_3 | 30 | 7 | 10 | 8 | 26.7 | -0.0134 | 7 | 23.3 | 2.2499 | 13.3 | 0.2478 | 0.3112 | 30 | Sufficient for analysis | Weak positive, NS |
| **Patient_3_1Granuloma_Q** | Patient_3 | 18 | 1 | 5.6 | 5 | 27.8 | -0.0014 | 5 | 27.8 | 2.828 | 22.2 | 0.8393 | 4.449e-05 | 18 | Sufficient for analysis | Strong positive, significant |
| **Patient_3_1Granuloma_S** | Patient_3 | 65 | 4 | 13.8 | 16 | 24.6 | -0.0441 | 16 | 24.6 | 1.7918 | 13.8 | 0.4027 | 0.002403 | 65 | Sufficient for analysis | Strong positive, significant |
| **Patient_3_1Granuloma_T** | Patient_3 | 26 | 7 | 0 | 7 | 26.9 | -0.0422 | 7 | 26.9 | 2.3979 | 3.8 | 0.1615 | 0.5559 | 26 | Sufficient for analysis | Weak positive, NS |
| **Patient_3_1Granuloma_U** | Patient_3 | 16 | 1 | 12.5 | 4 | 25 | 0.0288 | 4 | 25 | 2.562 | 12.5 | 0.2415 | 0.49 | 16 | Insufficient hypoxia-high spots | Weak positive, NS |
| **Patient_3_1Granuloma_V** | Patient_3 | 37 | 1 | 21.6 | 9 | 24.3 | 0.0284 | 9 | 24.3 | 2.5649 | 18.9 | 0.7055 | 3.821e-06 | 37 | Sufficient for analysis | Strong positive, significant |
| **Patient_3_1Granuloma_W** | Patient_3 | 59 | 4 | 0 | 15 | 25.4 | -0.054 | 13 | 22 | 1.6094 | 6.8 | 0.0393 | 0.8688 | 59 | Sufficient for analysis | Weak positive, NS |
| **Patient_3_1Granuloma_X** | Patient_3 | 14 | 4 | 0 | 1 | 7.1 | -0.0835 | 3 | 21.4 | 1.3863 | 0 | -0.0359 | 0.9506 | 14 | Insufficient hypoxia-high spots | Negative, NS |
| **Patient_3_2Granuloma_A** | Patient_3 | 506 | 3 | 41.1 | 127 | 25.1 | 0.0746 | 126 | 24.9 | 2.6391 | 18 | 0.7266 | 7.061e-83 | 506 | Sufficient for analysis | Strong positive, significant |
| **Patient_3_2Granuloma_B** | Patient_3 | 226 | 1 | 28.8 | 57 | 25.2 | 0.043 | 53 | 23.5 | 2.3979 | 15.5 | 0.6768 | 1.248e-30 | 226 | Sufficient for analysis | Strong positive, significant |
| **Patient_3_2Granuloma_C** | Patient_3 | 49 | 7 | 0 | 12 | 24.5 | 0.0153 | 10 | 20.4 | 1.0986 | 8.2 | 0.1786 | 0.3466 | 49 | Sufficient for analysis | Weak positive, NS |
| **Patient_3_2Granuloma_D** | Patient_3 | 654 | 1 | 25.8 | 164 | 25.1 | 0.0424 | 160 | 24.5 | 2.5649 | 17.9 | 0.7508 | 4.569e-118 | 654 | Sufficient for analysis | Strong positive, significant |
| **Patient_3_2Granuloma_E** | Patient_3 | 81 | 6 | 17.3 | 20 | 24.7 | 0.0752 | 20 | 24.7 | 2.4849 | 14.8 | 0.451 | 7.157e-05 | 81 | Sufficient for analysis | Strong positive, significant |
| **Patient_3_2Granuloma_F** | Patient_3 | 12 | 5 | 0 | 0 | 0 | -0.2055 | 2 | 16.7 | 1.3863 | 0 | 0.6478 | 0.04404 | 12 | Hypoxia-negative | Strong positive, significant |
| **Patient_3_2Granuloma_G** | Patient_3 | 17 | 2 | 0 | 4 | 23.5 | -0.0069 | 4 | 23.5 | 2.4849 | 0 | -0.4416 | 0.1341 | 17 | Insufficient hypoxia-high spots | Negative, NS |
| **Patient_3_2Granuloma_H** | Patient_3 | 46 | 2 | 10.9 | 12 | 26.1 | 0.0226 | 12 | 26.1 | 2.7726 | 8.7 | 0.0883 | 0.6715 | 46 | Sufficient for analysis | Weak positive, NS |
| **Patient_3_2Granuloma_I** | Patient_3 | 40 | 4 | 27.5 | 10 | 25 | 0.0489 | 10 | 25 | 2.6736 | 20 | 0.7194 | 6.758e-07 | 40 | Sufficient for analysis | Strong positive, significant |
| **Patient_3_2Granuloma_J** | Patient_3 | 10 | 4 | 0 | 0 | 0 | -0.1443 | 2 | 20 | 1.3863 | 0 | 0.1111 | 0.8688 | 10 | Hypoxia-negative | Weak positive, NS |
| **Patient_3_2Granuloma_K** | Patient_3 | 15 | 4 | 0 | 4 | 26.7 | -0.0306 | 4 | 26.7 | 0.6931 | 6.7 | 0.3261 | 0.3533 | 15 | Insufficient hypoxia-high spots | Strong positive, underpowered |
| **Patient_3_2Granuloma_L** | Patient_3 | 34 | 5 | 14.7 | 9 | 26.5 | -0.0408 | 9 | 26.5 | 1.0986 | 8.8 | 0.1633 | 0.49 | 34 | Sufficient for analysis | Weak positive, NS |
| **Patient_3_2Granuloma_M** | Patient_3 | 54 | 3 | 61.1 | 14 | 25.9 | 0.147 | 14 | 25.9 | 3.155 | 13 | 0.6353 | 9.233e-07 | 54 | Sufficient for analysis | Strong positive, significant |
| **Patient_3_2Granuloma_N** | Patient_3 | 17 | 5 | 0 | 4 | 23.5 | -0.1013 | 3 | 17.6 | 1.0986 | 5.9 | -0.0489 | 0.9297 | 17 | Insufficient hypoxia-high spots | Negative, NS |
| **Patient_4_1Granuloma_A** | Patient_4 | 1889 | 7 | 18.9 | 472 | 25 | 0.0467 | 403 | 21.3 | 1.6094 | 11.3 | 0.3331 | 5.164e-49 | 1889 | Sufficient for analysis | Strong positive, significant |
| **Patient_4_1Granuloma_B** | Patient_4 | 30 | 6 | 20 | 8 | 26.7 | 0.069 | 8 | 26.7 | 2.191 | 16.7 | 0.585 | 0.001955 | 30 | Sufficient for analysis | Strong positive, significant |
| **Patient_4_1Granuloma_C** | Patient_4 | 14 | 4 | 7.1 | 4 | 28.6 | 0.0354 | 3 | 21.4 | 1.3863 | 21.4 | 0.3661 | 0.321 | 14 | Insufficient hypoxia-high spots | Strong positive, underpowered |
| **Patient_4_1Granuloma_D** | Patient_4 | 79 | 6 | 13.9 | 20 | 25.3 | 0.0822 | 20 | 25.3 | 1.7918 | 11.4 | 0.2988 | 0.01603 | 79 | Sufficient for analysis | Weak positive, significant |
| **Patient_4_1Granuloma_E** | Patient_4 | 69 | 3 | 52.2 | 17 | 24.6 | 0.1401 | 17 | 24.6 | 2.3026 | 11.6 | 0.4887 | 6.447e-05 | 69 | Sufficient for analysis | Strong positive, significant |
| **Patient_4_1Granuloma_F** | Patient_4 | 62 | 1 | 8.1 | 16 | 25.8 | 0.0156 | 15 | 24.2 | 1.3863 | 8.1 | 0.117 | 0.49 | 62 | Sufficient for analysis | Weak positive, NS |
| **Patient_4_2Granuloma_A** | Patient_4 | 1700 | 2 | 1.9 | 425 | 25 | 0.016 | 364 | 21.4 | 1.7918 | 7.4 | 0.1438 | 1.186e-08 | 1700 | Sufficient for analysis | Weak positive, significant |
| **Patient_4_2Granuloma_B** | Patient_4 | 63 | 6 | 1.6 | 16 | 25.4 | 0.024 | 12 | 19 | 1.7918 | 3.2 | 0.0189 | 0.9462 | 63 | Sufficient for analysis | Weak positive, NS |
| **Patient_4_2Granuloma_C** | Patient_4 | 24 | 6 | 0 | 3 | 12.5 | -0.1283 | 3 | 12.5 | 1.0986 | 0 | 0.0685 | 0.8688 | 24 | Insufficient hypoxia-high spots | Weak positive, NS |
| **Patient_4_2Granuloma_D** | Patient_4 | 36 | 1 | 11.1 | 9 | 25 | 0.0357 | 9 | 25 | 2.3026 | 13.9 | 0.4176 | 0.02333 | 36 | Sufficient for analysis | Strong positive, significant |
| **Patient_4_2Granuloma_E** | Patient_4 | 31 | 2 | 0 | 8 | 25.8 | 0.0491 | 8 | 25.8 | 2.0794 | 6.5 | 0.0037 | 1 | 31 | Sufficient for analysis | Weak positive, NS |

Supplementary Table 7

| Antigen | Catalogue Number | Panel | Host Species | Clone | Isotype | Manufacturer | Dilution | Permeabilization | Secondary |
|---|---|---|---|---|---|---|---|---|---|
| **CD163** | OTI2B12 | Myeloid | mouse | Monoclonal | IgG2a | Origene | 1;100 | No | Goat anti-Mouse IgG2a Cross-Adsorbed Secondary Antibody, Alexa Fluor™ 633/ |
| **FCGR1A(CD64)** | OTI1A8 | Myeloid | Mouse | Monoclonal | IgG1 | origene | 1;100 | No | Goat anti-Mouse IgG1 Cross-Adsorbed Secondary Antibody, Alexa Fluor™ 647 |
| **IDO1** | UMAB126 | Myeloid | Mouse | Monoclonal | IgG1 | Origene | 1;100 | Yes | Goat anti-Mouse IgG1 Cross-Adsorbed Secondary Antibody, Alexa Fluor™ 647 |
| **GLUT1** | 6513-MSM5 | Hypoxia | Mouse | Monoclonal | IgG2b | NeoBiotechnologies | 1:50 | No | Goat anti-Mouse IgG2b Cross-Adsorbed Secondary Antibody, Alexa Fluor™ 488 |
| **HIF1A** | 3091-MSM1 | Hypoxia | Mouse | Monoclonal | IgG2b | NeoBiotechnologies | 1;100 | Yes | Goat anti-Mouse IgG2b Cross-Adsorbed Secondary Antibody, Alexa Fluor™ 488 |
| **SLC16A3** | PA5-54413 | Hypoxia | Rabbit | Polyclonal | IgG | Invitrogen | 1:50 | No | Donkey anti-Rabbit IgG (H+L) Highly Cross-Adsorbed Secondary Antibody, Alexa Fluor™ 546 |
| **CA9** | PA5-47268 | Hypoxia | Goat | Polyclonal | IgG | Invitrogen | 15ug/ml | No | Donkey anti-Goat IgG (H+L) Cross-Adsorbed Secondary Antibody, Alexa Fluor™ 680 |
| **SPP1** | PA5-141129 | Hypoxia | Rabbit | Polyclonal | IgG | Invitrogen | 1:50 | Yes | Donkey anti-Rabbit IgG (H+L) Highly Cross-Adsorbed Secondary Antibody, Alexa Fluor™ 546 |
| **CD4** | ab133616 | Lymphoid | Rabbit | Monoclonal | IgG | Abcam | 1;100 | No | Donkey anti-Rabbit IgG (H+L) Highly Cross-Adsorbed Secondary Antibody, Alexa Fluor™ 546 |
| **CD3** | ab16669 | Lymphoid | Rabbit | Monoclonal | IgG | Abcam | 1;100 | No | Donkey anti-Rabbit IgG (H+L) Highly Cross-Adsorbed Secondary Antibody, Alexa Fluor™ 546 |
| **CD8** | IR623 | Lymphoid | mouse | Monoclonal | IgG1 | Dako | ready to use | No | Goat anti-Mouse IgG1 Cross-Adsorbed Secondary Antibody, Alexa Fluor™ 647 |
